# Long read multi-omics sequencing reveals DNA-to-RNA evolutionary remodeling trajectories of recurrent astrocytoma

**DOI:** 10.64898/2026.07.12.738108

**Authors:** Sherif Rashad, Daisuke Ando, Shota Yamashita, Yoshiteru Shimoda, Masayuki Kanamori, Hidenori Endo, Kuniyasu Niizuma

## Abstract

Astrocytoma recurrence is shaped by genetic, epigenetic and transcriptional evolution, but how DNA remodeling is propagated into full-length RNA isoforms and coding consequences remains poorly resolved. Here we applied paired PacBio HiFi whole-genome sequencing and Kinnex full-length RNA sequencing to matched primary and recurrent astrocytoma from six patients, with matched blood controls. Recurrent tumors preserved core glioma driver identity while acquiring patient-specific remodeling across somatic variants, copy number, structural variation, loss of heterozygosity, haplotype imbalance and DNA methylation. Long-read transcriptomics revealed extensive recurrence-associated gene-expression, isoform-usage, differential transcript-usage and predicted ORF/protein-fate remodeling beyond gene-level expression. A denominator-aware integration framework showed that copy-number changes provide broad RNA-dosage links, whereas expressed somatic variants, allele/haplotype-specific transcript usage, methylation-linked isoform remodeling and structural variants generate more focused DNA–RNA–ORF chains. These findings establish longitudinal long-read multi-omics as a framework for prioritizing patient-specific remodeling trajectories and identifying targets for precision medicine in recurrent astrocytoma.

## Introduction

Gliomas are the most common malignant primary brain tumors in adults and remain a major cause of cancer-related morbidity and mortality in neuro-oncology^1^. Although molecular classification and standard treatment have improved the diagnosis and management of diffuse gliomas, recurrence remains a central clinical problem^2,3^. Recurrent gliomas are not simple regrowth of the original tumor. Instead, they emerge through longitudinal remodeling across genetic, epigenetic, transcriptional, microenvironmental, and post-transcriptional layers^4–8^. Large-scale and longitudinal studies have shown that glioma progression is shaped by somatic evolution, treatment-associated selection, DNA methylation remodeling, transcriptional state transitions, and interactions with the tumor microenvironment^4–7^. Nonetheless, connecting these regulatory layers within the same patient remains difficult, especially when the molecular consequences of DNA remodeling extend beyond gene-level expression. Likewise, Recent longitudinal studies have begun to define disease-specific routes of IDH-mutant astrocytoma progression. Several studies have shown progressive genomic, epigenomic, transcriptomic, and proteomic remodeling changes that drive IDH mutant astrocytoma recurrence, cell state change, and immune evasiveness^8–11^. However, these studies do not resolve how these alterations are expressed through full-length transcript structures, allele- or haplotype-specific isoform usage, and isoform-specific coding consequences within the same patient.

Despite these advances, an important technical gap remains. Short-read DNA and RNA sequencing have been indispensable for defining glioma/astrocytoma driver alterations, copy-number states, mutational processes, gene-expression programs, and DNA methylation classes. However, short reads provide only fragmented views of many molecular features that are central to tumor evolution, including complex structural variants, allele-specific events, long-range haplotypes, phased methylation states, full-length transcript isoforms, fusion transcripts, and isoform-specific coding consequences. As a result, most studies still analyze genome, epigenome, transcriptome, and predicted protein-output changes as partially separate layers. This limits the ability to ask a mechanistic question that is central to recurrence biology: which genomic and epigenomic alterations are reflected at patient-matched RNA, isoform, allele-specific, and predicted open reading frames (ORF)/protein-fate levels?

Long-read sequencing provides an opportunity to address this gap. Long-read RNA sequencing can capture full-length transcripts, enabling direct characterization of isoform diversity, alternative transcript usage, fusion transcripts, and coding-potential changes that are difficult to infer from short-read junction counts alone^12,13^. Recent cancer studies have shown that long-read transcriptomics can uncover extensive previously unannotated isoform diversity, identify cancer-associated transcript structures, resolve isoform-level expression programs, and reveal genomic alterations or fusions that may be misinterpreted by short-read RNA-seq^12,13^. Long-read DNA sequencing, although adopted more slowly in cancer because of cost, input requirements, and analytical complexity, can resolve structural variants, haplotypes, methylation landscapes, repetitive-element insertions, and complex rearrangements that are often inaccessible or ambiguous with short-read data^14,15^. These properties make long-read sequencing particularly well suited for studying recurrence, where DNA remodeling, epigenetic state changes, and transcriptome reprogramming may converge on patient-specific molecular outcomes.

Despite these advances, the full potential of paired long-read DNA and RNA sequencing remains underused in cancer. Many long-read RNA studies annotate novel isoforms and predict open reading frames but rarely connect these isoforms to patient-matched DNA alterations, allele or haplotype context, nonsense-mediated decay (NMD) potential, frameshift status, or altered proteoform structure. Conversely, long-read DNA studies can resolve complex genome architecture and methylation states, but their transcriptomic consequences are often inferred indirectly, combined with short read sequencing which cannot capture many of the complex phenomena resolved by long read DNA sequencing^15^, or not examined with matched full-length RNA data^14,15^. Thus, a key unresolved problem is not only whether recurrent gliomas acquire DNA and RNA alterations, but how often these alterations form coherent DNA-to-RNA-to-ORF remodeling chains.

Here, we apply paired long-read DNA and RNA sequencing to matched primary and recurrent astrocytomas from six patients, with matched blood samples used as germline controls for DNA analysis. We first define the recurrent astrocytoma DNA landscape using long-read genome sequencing, including somatic variants, structural rearrangements, copy-number alterations, loss of heterozygosity, and DNA methylation remodeling. We then use long-read RNA sequencing to characterize recurrence-associated changes in gene expression, isoform diversity, transcript usage, junction architecture, and predicted ORF/protein fate. Finally, we integrate the DNA and RNA layers to determine how genomic, and epigenomic remodeling is reflected at patient-matched RNA, isoform, allele/haplotype-specific, and predicted protein-output levels. This analysis reveals that recurrent astrocytoma remodeling is frequently expressed through isoform and ORF-level consequences that are not captured by gene-level expression alone and provides a denominator-aware framework for prioritizing high-confidence DNA-RNA-ORF remodeling chains in individual patients, providing basis for precision oncology and patient-specific therapeutics.

## Results

### Cohort, study design, and sequencing quality control

We selected six patients with matched primary and recurrent astrocytoma specimens collected during routine surgical resection, together with matched blood samples for germline DNA control. Samples were selected after RNA and DNA quality screening to ensure suitability for long-read genome and transcriptome sequencing (Figure 1a). The cohort included four males and two females, with age at primary tumor presentation ranging from 24 to 45 years. All patients underwent debulking surgery. Except one patient, none had received either radiotherapy or chemotherapy before recurrence. One patient had undergone postoperative concomitant radio therapy with temozolomide. Four patients showed increased tumor grade at recurrence, and all tumors carried IDH1 mutation. Additional clinical and molecular features are summarized in Supplementary Table 1. Integrated diagnoses were made in accordance with the 5th edition of the WHO Classification of Tumors of the Central Nervous System^3^.

**Figure 1.**
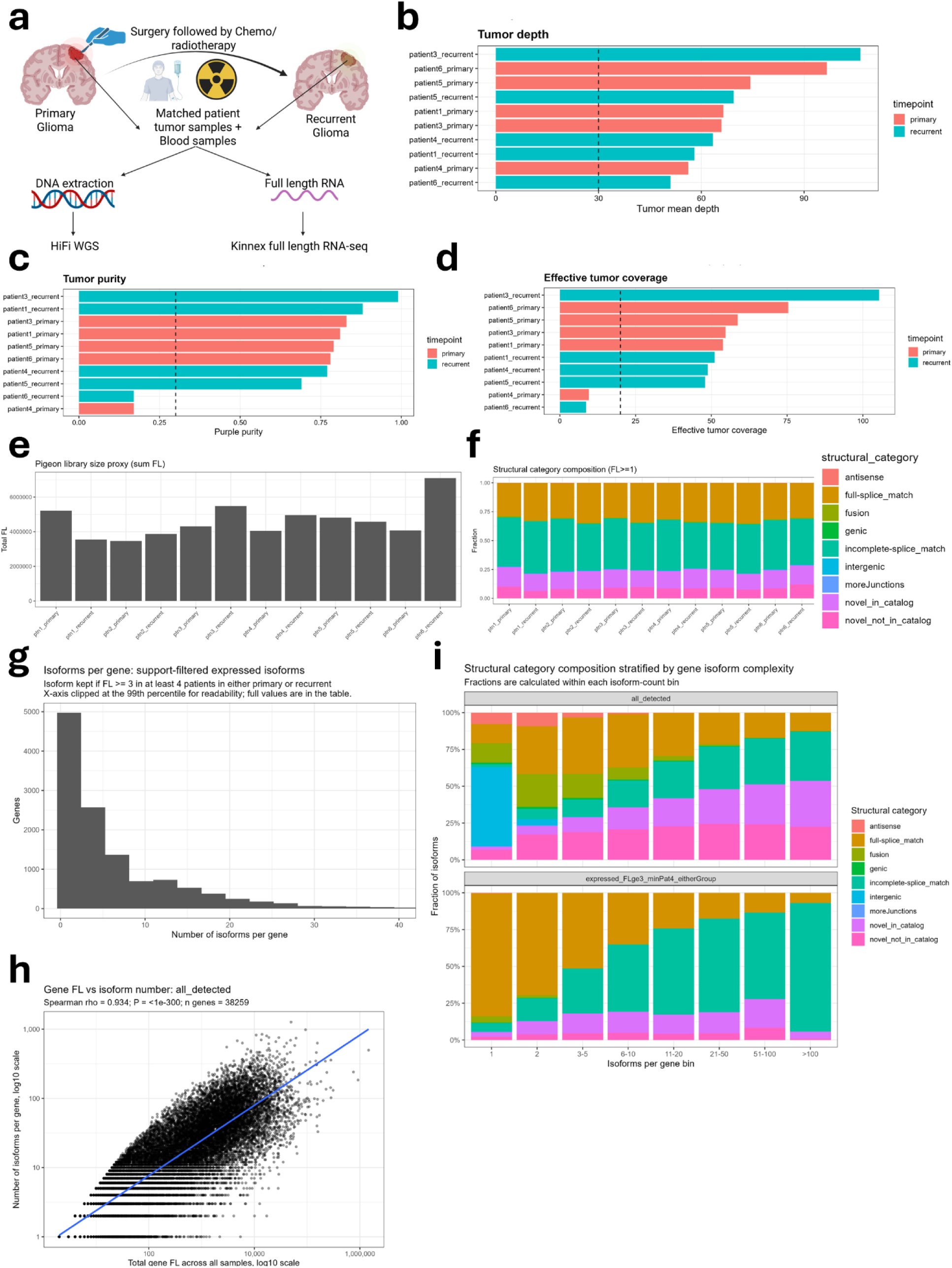
Longitudinal long-read DNA/RNA cohort design and sequencing quality control. **a,** Study design. Matched primary and recurrent glioma specimens were collected from six patients, together with matched blood samples for germline DNA control. Tumor and blood DNA were profiled using PacBio HiFi whole-genome sequencing, while matched tumor RNA was profiled using Kinnex full-length RNA sequencing. **b,** Mean tumor sequencing depth across tumor DNA samples. The recurrent sample from patient 2 had low sequencing depth and was excluded from DNA-dependent analyses. **c,** Tumor purity estimated from DNA sequencing. **d,** Effective tumor coverage after accounting for tumor purity. **e,** Processed full-length RNA read support per sample used for isoform-level analyses. **f,** Pigeon structural-category composition across RNA samples, showing broadly comparable transcript annotation profiles across the cohort. **g,** Distribution of the number of detected isoforms per gene among support-filtered expressed isoforms. **h,** Relationship between gene-level full-length read support and the number of detected isoforms per gene. **i,** Structural-category composition stratified by gene-level isoform complexity, showing increased representation of non-canonical transcript categories among genes with higher isoform counts.

We performed PacBio HiFi whole-genome sequencing on tumor and matched blood DNA, and Kinnex full-length RNA sequencing on matched tumor RNA (Figure 1a). Tumor sequencing depth was sufficient for DNA-dependent analyses in five of six recurrent-primary pairs, with most tumor samples reaching approximately 60× or higher mean depth (Figure 1b; Supplementary Table 1). The recurrent sample from patient 2 had substantially lower tumor depth because of a technical sequencing issue and therefore that patient was excluded from DNA-dependent analyses, while retained for RNA-based analyses (Supplementary table 1). Tumor purity and effective tumor coverage were acceptable for most samples, although lower purity in patient 4 primary and patient 6 recurrent tumors was noted and considered during downstream interpretation (Figure 1c,d).

Kinnex full-length RNA sequencing was performed at 10 million reads per sample and produced 4-8 million mapped and processed full-length (FL) reads per sample, providing sufficient depth for isoform-level analyses across the cohort (Figure 1e). Pigeon transcript classification showed broadly comparable structural-category composition across primary and recurrent samples, without evidence of a major sample-level technical shift in transcript annotation classes (Figure 1f). Across genes, the number of detected isoforms was highly variable: most genes were represented by a small number of isoforms, whereas a subset showed extensive isoform diversity (Figure 1g). Gene-level full-length read support was strongly associated with the number of detected isoforms, indicating that read depth contributes to isoform discovery and must be considered when interpreting isoform richness (Figure 1h). Genes with higher isoform counts also showed a larger fraction of non-canonical transcript categories, consistent with increased transcript-structure diversity among highly complex genes (Figure 1i).

Together, these quality-control analyses established a longitudinal long-read DNA/RNA cohort suitable for downstream analysis of recurrent astrocytoma genome remodeling, transcriptome remodeling, and patient-matched DNA-RNA integration. Because patient 2 recurrent tumor lacked sufficient DNA coverage, DNA-dependent recurrence analyses were restricted to the five DNA-evaluable patient pairs, whereas RNA analyses included all six patients.

### Long-read DNA sequencing reveals multi-layer genome remodeling in recurrent astrocytoma

We next used matched HiFi whole-genome sequencing to define the somatic DNA landscape of primary and recurrent astrocytomas. Across the paired cases, somatic SNV/indel analysis showed shared, primary-only, and recurrent-only variants, indicating clonal relatedness between primary and recurrent tumors together with ongoing patient-specific evolution (Fig. 2a). A driver-focused SNV/indel heatmap confirmed the expected glioma genetic backbone, including recurrent involvement of canonical glioma genes such as IDH1, TP53, ATRX, CIC, and other cancer-associated loci, while also showing patient-specific driver-context events (Fig. 2b). However, paired SNV/indel burden, median variant allele fraction, and read depth did not show a uniform recurrence-associated increase across patients (Fig. 2c). Variant-level VAF and depth distributions supported the reliability of the small-variant calls, while SBS96 context shifts showed patient-specific mutational-context changes rather than one shared recurrence-wide mutational pattern (Supplementary Fig. 1a,b). Thus, although small variants provide evidence of tumor lineage and patient-specific evolution, they do not by themselves explain the recurrent DNA phenotype.

**Figure 2.**
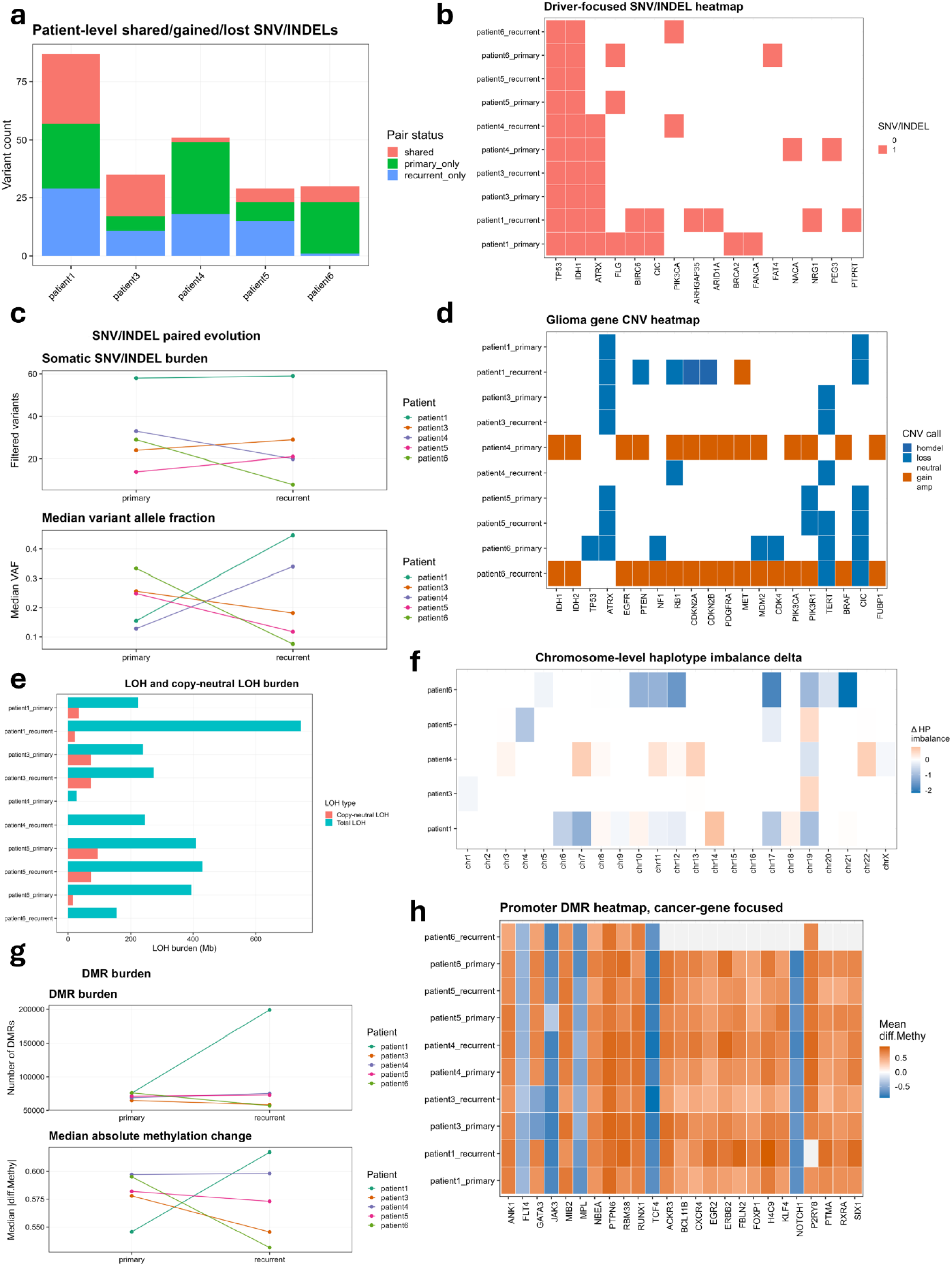
Long-read DNA sequencing reveals multi-layer small-variant, copy-number, allelic, and methylation remodeling during astrocytoma recurrence. **a,** Patient-level distribution of shared, primary-only, and recurrent-only somatic SNV/indel events across paired evaluable DNA samples. **b,** Driver-focused SNV/indel heatmap showing presence or absence of somatic SNV/indel events in recurrently altered glioma/cancer-associated genes across primary and recurrent samples. **c,** Paired SNV/indel evolution summary showing somatic SNV/indel burden, median variant allele fraction, and median read depth in primary and recurrent tumors. **d,** Glioma gene copy-number heatmap showing gain, loss, and homozygous deletion-like copy-number states across selected glioma-associated genes. **e,** LOH and copy-neutral LOH burden across primary and recurrent tumors. **f,** Chromosome-level recurrent-primary change in haplotype imbalance, summarized per patient and chromosome. **g,** DMR burden and median absolute methylation change across paired primary and recurrent tumors. **h,** Cancer-gene-focused promoter DMR heatmap showing mean differential methylation across selected cancer-associated genes and samples.

We therefore examined larger-scale copy-number and allelic remodeling. Gene copy-number profiling revealed patient-specific gains, losses, and homozygous deletion-like states across canonical glioma-associated genes, including receptor tyrosine kinase, PI3K pathway, cell-cycle, and tumor suppressor loci (Fig. 2d). Loss of heterozygosity (LOH) analysis further showed substantial allelic remodeling across samples, with total LOH exceeding copy-neutral LOH in most tumors and marked variation between primary and recurrent disease (Fig. 2e). Chromosome-level haplotype imbalance analysis identified recurrent-primary shifts affecting selected chromosomes rather than a uniform genome-wide direction of change (Fig. 2f). Additional haplotype-aware analyses showed chromosome-level dominant haplotype switching, ranked chromosome-level haplotype imbalance shifts, and patient-specific changes in CN-LOH, genome-wide haplotype imbalance, ploidy, purity, subclonal burden, and total LOH (Supplementary Fig. 1c–e). These results indicate that astrocytoma recurrence is accompanied by allele-aware copy-number remodeling, including changes in LOH and haplotype imbalance that are not fully captured by gene-level CNV calls alone.

Methylation analysis revealed a third major layer of DNA remodeling. Differentially methylated region burden varied strongly across patients, with patient 1 showing a prominent recurrent increase in DMR number and median absolute methylation change, whereas other patients showed more modest or opposite shifts (Fig. 2g). Cancer-gene-focused promoter DMR analysis revealed broad promoter methylation differences across primary and recurrent tumors, including both hypermethylated and hypomethylated states across genes involved in signaling, transcriptional regulation, and tumor biology (Fig. 2h). A focused analysis of shared promoter genes with the strongest primary-recurrent methylation shifts further highlighted patient-specific promoter methylation remodeling (Supplementary Fig. 2a). DMRs were detected across genomic contexts, with the largest absolute burden in intronic regions, while promoter and exon-associated DMRs provided more directly interpretable regulatory contexts (Supplementary Fig. 2b). Signal-level methylation summaries supported the DMR calls: promoter bigWig signal shifts differed by haplotype context, DMR-derived methylation differences showed high agreement with bigWig signal, and most promoter regions with bigWig support overlapped haplotype-resolved regions rather than no-overlap regions (Supplementary Fig. 2c–e). Across samples, promoter methylation signal shifted between primary and recurrent tumors, absolute promoter methylation change varied by patient, DMR-derived methylation estimates correlated strongly with bigWig signal, and genome-wide DMR signal showed patient-specific primary-recurrent remodeling (Supplementary Fig. 2f–i). Together, these data show that recurrent astrocytomas undergo broad, patient-specific DNA methylation remodeling that can be linked to promoter and haplotype contexts.

### Structural, copy-number, and methylation alterations define recurrently remodeled genomic substrates

We then focused on structural genome remodeling, which represented one of the richest long-read DNA signals. SV burden varied markedly across patients, with some tumors showing reduced SV burden at recurrence and others showing persistent or increased structural alteration (Fig. 3a). Representative circos plots for patient 1 illustrated extensive primary structural rearrangement and a remodeled recurrent structural landscape, supporting the idea that recurrence is not simply a linear accumulation of identical structural events (Fig. 3b). At the chromosome level, structural complexity was concentrated in selected patient-chromosome combinations rather than distributed evenly across the cohort (Fig. 3c). SV type composition showed that deletions, insertions, and duplications formed the major classes of detected SVs, with lower relative contributions from inversions and breakend-like events in most samples (Fig. 3d). These findings point to patient-specific structural genome remodeling as a major substrate of recurrent astrocytoma evolution.

**Figure 3.**
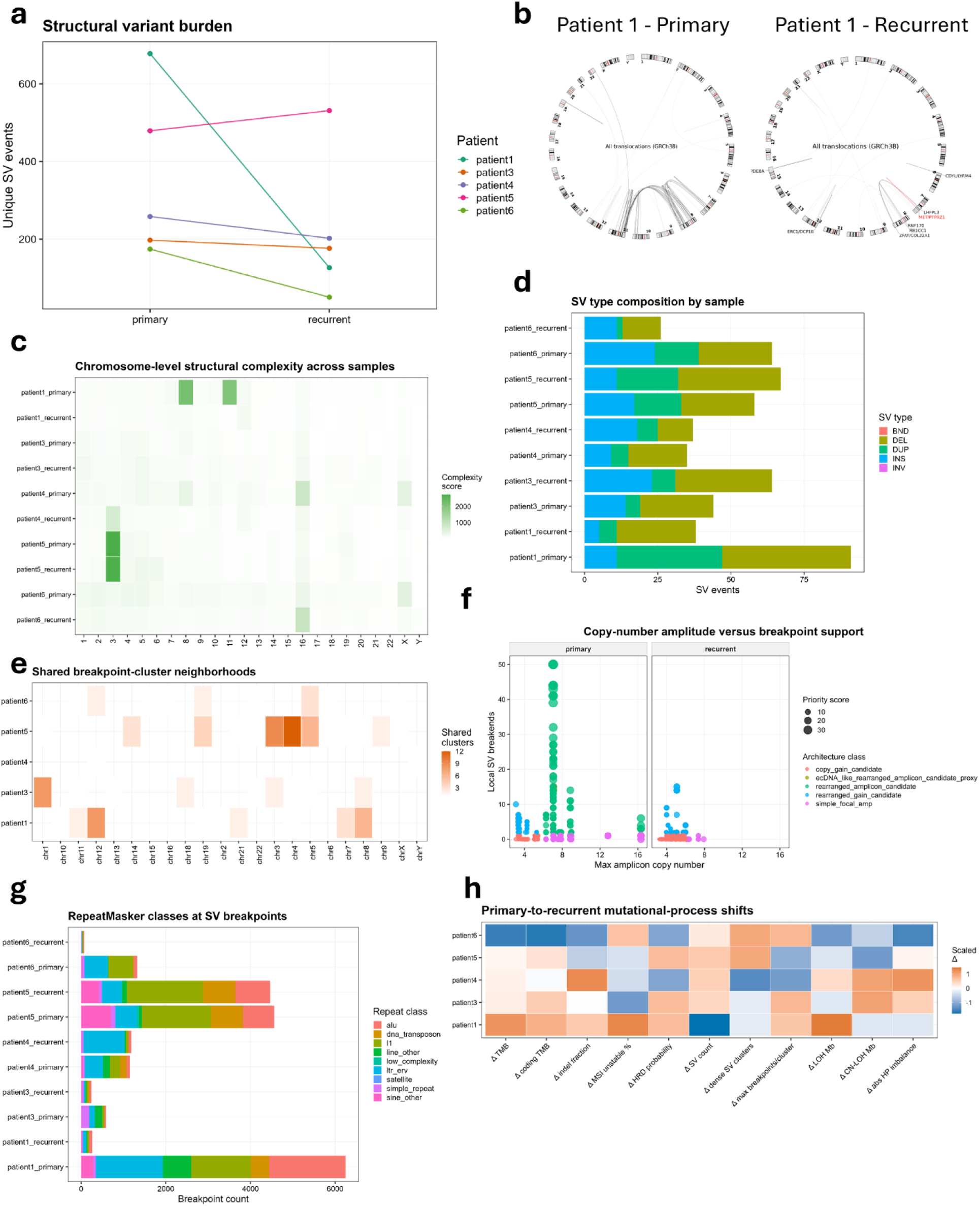
Long-read DNA sequencing identifies structural rearrangement, focal amplicon, and repeat-associated genome remodeling during astrocytoma recurrence. **a,** Structural variant burden across paired primary and recurrent tumors. **b,** Representative circos plots showing translocation/rearrangement architecture in patient 1 primary and recurrent tumors. **c,** Chromosome-level structural complexity across samples, showing patient- and chromosome-specific concentration of complex rearrangement signal. **d,** SV type composition by sample, including deletion, duplication, insertion, inversion, and breakend-like classes. **e,** Shared breakpoint-cluster neighborhoods across patients and chromosomes, summarizing regional structural instability rather than exact breakpoint identity. **f,** Copy-number amplitude versus local breakpoint support for focal amplicon-like regions, stratified by timepoint and structural architecture class. **g,** RepeatMasker classes at SV breakpoints, showing the repeat contexts intersected by structural rearrangement breakpoints. **h,** Primary-to-recurrent mutational-process shifts across patients.

To determine whether recurrent disease preserved the same structural loci or instead remodeled related genomic neighborhoods, we examined breakpoint-cluster sharing. Breakpoint-cluster neighborhoods showed recurrently involved chromosomal regions across patient pairs, suggesting that some structurally unstable regions persist or re-emerge even when exact breakpoints are not retained (Fig. 3e). Exact private breakpoint events were highly patient- and timepoint-specific, with large private SV burdens in selected primary or recurrent samples (Supplementary Fig. 3a). Complex rearrangement candidate classes, including clustered rearrangement, dense breakpoint cluster, rearranged amplicon candidate, and chromothripsis-like candidate classes, further showed that structural architecture varied across samples (Supplementary Fig. 3b). These labels should be interpreted as structural proxy classes rather than formally validated mechanisms, but they show that recurrent astrocytoma’s structural remodeling often occurs at the level of regional genome architecture rather than isolated SV events alone.

We next evaluated focal amplicon and rearranged copy-number architecture. Copy-number amplitude plotted against local SV breakpoint support identified amplified regions with variable degrees of structural support, including regions with high copy number and local breakpoint enrichment (Fig. 3f). Refined driver/onco-amplicon gene copy-number calls highlighted patient-specific focal copy-number gains involving cancer-relevant loci, providing a direct link between structural genome remodeling and oncogenic dosage architecture (Supplementary Fig. 3c). CCG-focused SV heatmaps further showed that SVs intersected selected cancer-associated genes in a patient-specific manner (Supplementary Fig. 3d). These results indicate that copy-number remodeling in recurrent astrocytoma includes not only broad CNV states but also structurally complex focal amplicon-like architectures.

We also examined repeat, mobile-element, and telomere-associated contexts as supporting annotations of the structural landscape. RepeatMasker annotation showed that SV breakpoints frequently intersected repeat contexts, including Alu, LINE/L1, LTR/ERV, DNA transposon, simple-repeat, low-complexity, and related repeat classes (Fig. 3g). RepeatMasker annotation of somatic indels showed related but not identical repeat-class distributions, and primary-to-recurrent mobile-element/repeat-context shifts were patient-specific rather than uniform (Supplementary Fig. 3e,f). Telomere-proximal breakage and telomere-window LOH only partially overlapped, suggesting that local telomere-proximal structural breakage and broader telomeric allelic imbalance are related but non-identical features of recurrence-associated genome instability (Supplementary Fig. 3g). These repeat and telomere analyses should be interpreted as genomic-context associations rather than definitive evidence of active mobile-element insertion or telomere-maintenance mechanism switching.

Finally, primary-to-recurrent mutational-process shifts were heterogeneous across patients, without evidence for one shared recurrence-wide repair phenotype (Fig. 3h). Overall, long-read DNA sequencing showed that recurrent astrocytomas preserve core driver identity while undergoing extensive patient-specific remodeling across small variants, CNV, LOH, haplotype imbalance, promoter methylation, SVs, focal amplicon architecture, repeat/telomere-associated genomic contexts, and mutational-process features. The dominant DNA-side conclusion is therefore not a single universal recurrence mechanism, but a heterogeneous remodeling landscape in which each recurrent tumor combines different genomic and epigenomic routes to generate a remodeled disease state. This multi-layer DNA substrate provides the foundation for the downstream long-read RNA and DNA-RNA integration analyses, where these genomic alterations can be connected to gene-expression, isoform, fusion, and ORF-level consequences.

### Long-read transcriptome sequencing reveals global changes in gene programs during astrocytoma recurrence

After defining the DNA remodeling landscape, we next asked whether recurrent tumors showed corresponding transcriptomic state changes using long-read RNA sequencing. We first analyzed gene-level expression by summing full-length (FL) Pigeon read counts per gene and fitting a paired primary-versus-recurrent model. This conventional gene-level analysis identified both upregulated and downregulated genes in recurrence (Fig. 4a). Over-representation analysis of recurrently upregulated genes highlighted mitochondrial and respiratory programs, including proton transmembrane transport, oxidative phosphorylation, electron transport chain, aerobic electron transport, ATP synthesis-coupled electron transport, and mitochondrial ATP synthesis-coupled electron transport (Fig. 4b). To complement single-gene differential expression, we performed sample-level program scoring using ssGSEA across Hallmark, Reactome, Oncogenic, and Immunologic gene-set collections. This analysis revealed patient-paired shifts in multiple tumor-state and microenvironment-related programs, including hypoxia, reactive oxygen species, epithelial–mesenchymal transition, UV-response-associated programs, interferon and inflammatory signatures, PDGF/ERK-related programs, IL15 response, and Rho GTPase signaling (Supplementary Fig. 4a–k). Because many program-level changes were nominal and did not uniformly survive global multiple-testing correction, we interpret these results as a paired tumor-state remodeling landscape rather than as a set of independently validated pathway claims. Together, this data shows that recurrent astrocytomas exhibit broad expression and pathway-state changes, but this analysis alone cannot resolve whether recurrence also remodels the structure and coding output of transcripts.

**Figure 4.**
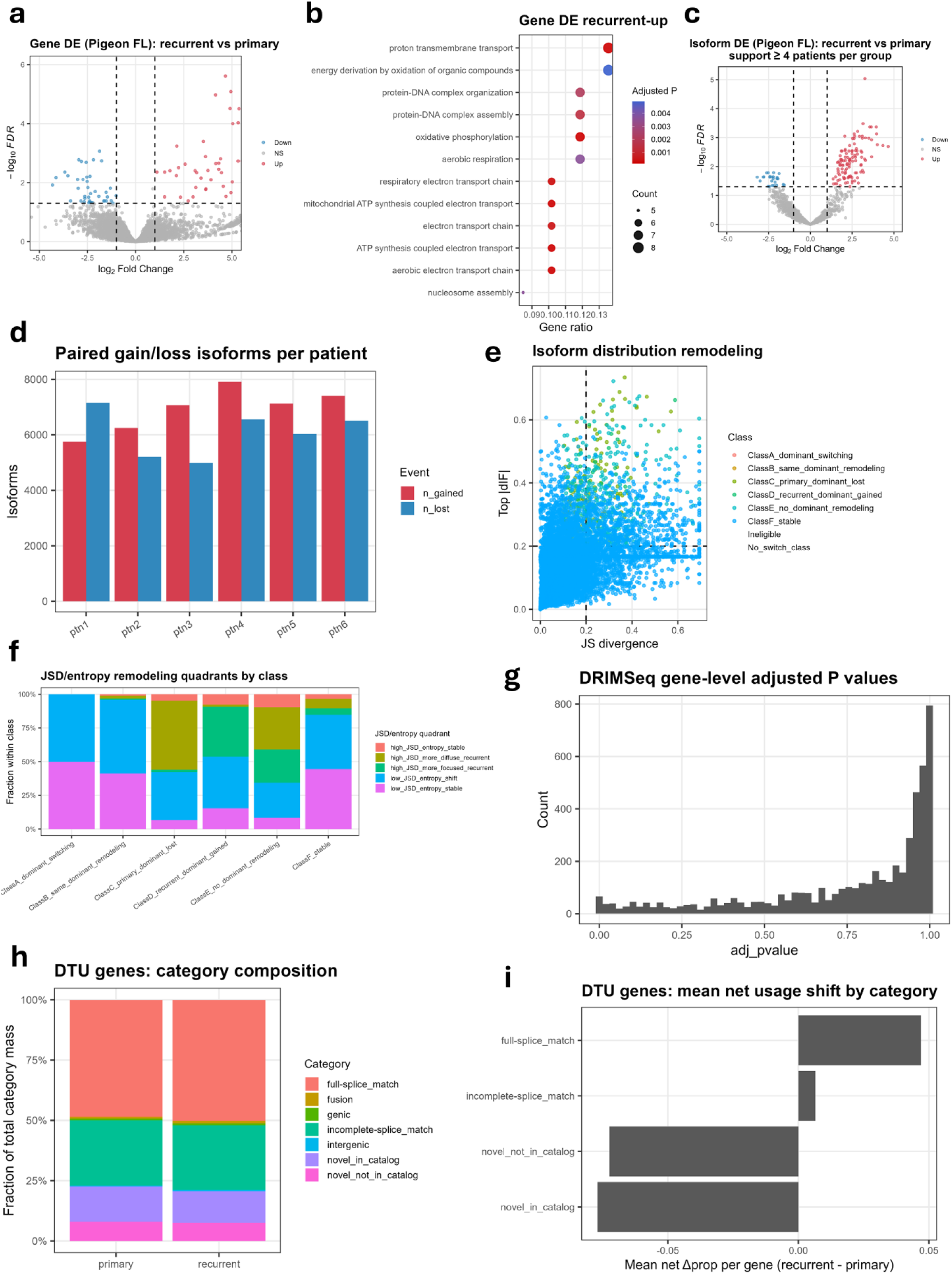
Long-read transcriptome sequencing reveals gene-program and isoform-usage remodeling during astrocytoma recurrence. **a,** Gene-level differential expression volcano plot comparing recurrent versus primary glioma using full-length Pigeon read counts summed per gene and modeled with a paired design. **b,** Gene Ontology biological process over-representation analysis of recurrently upregulated genes, highlighting mitochondrial respiration, electron transport, oxidative phosphorylation, proton transmembrane transport, and ATP synthesis-associated programs. **c,** Isoform-level differential expression volcano plot comparing recurrent versus primary tumors using support-filtered full-length isoform counts. **d,** Paired recurrent gain and loss of expressed isoforms per patient using high-confidence isoform support filters. **e,** Gene-level isoform remodeling map based on Jensen–Shannon divergence and maximum absolute delta isoform fraction. Dashed lines indicate the high-confidence thresholds used for remodeling calls: JSD ≥ 0.1 and max |dIF| ≥ 0.2, with patient support ≥4. **f,** JSD–entropy quadrant composition across A–F isoform remodeling classes. JSD captures overall primary-recurrent isoform-distribution divergence, whereas recurrent-primary Shannon entropy shift captures whether recurrent isoform usage becomes more diffuse, more focused, or remains similarly distributed. **g,** DRIMSeq gene-level adjusted P-value distribution from paired differential transcript usage analysis. **h,** Gene-weighted Pigeon structural-category composition among DTU genes in primary and recurrent tumors. **i,** Mean net recurrent-primary DTU usage shift by Pigeon structural category, showing category-level directionality of transcript usage changes.

### Long-read transcriptome sequencing reveals extensive isoform remodeling independent of apparent gene expression changes

We therefore next examined isoform-level remodeling directly. Differential isoform expression analysis using paired primary-vs-recurrent DEseq2-based analysis revealed sets of up and downregulated isoforms in recurrent tumors (Fig. 4c). However, these changes cannot be completely separated from global changes in gene expression, and they fail in capturing the complexity of gene isoform remodeling beyond simple changes in expression. Therefore, next we examined isoform gain/loss in recurrent tumors, restricting our criteria to isoforms with at least 5 FL reads that are present in at least 4 patient samples in either primary or recurrent tumors, and that represent at least 10% of the total gene FL reads. The paired isoform gain/loss analysis showed that each patient acquired and lost thousands of detected isoforms between primary and recurrent tumors, indicating extensive turnover of the expressed isoform repertoire across recurrence (Fig. 4d). Structural-category enrichment analysis of high-confidence gained and lost isoforms showed that recurrently gained isoforms were enriched in full-splice match and incomplete-splice match categories, whereas lost isoforms enriched in novel isoforms either in catalog or not in catalog (Supplementary Fig. 5a). These findings indicate that recurrence-associated transcriptome remodeling cannot be reduced to simple changes in isoform abundance.

To capture gene-level isoform remodeling more comprehensively, we used to independent statistical approaches. In the first approach, we classified each gene using a model based on isoform fractions rather than absolute abundance alone. First, we defined the dominant isoform per gene across primary and recurrent group (see methods). This framework allowed us to separate genes into six remodeling classes: Class A, dominant isoform switching, where a dominant isoform is present in both groups but changes identity; Class B, same-dominant remodeling, where the dominant isoform is preserved but secondary isoforms remodel; Class C, primary-dominant lost, where a dominant primary isoform is no longer dominant in recurrence; Class D, recurrent-dominant gained, where a dominant isoform emerges in recurrence; Class E, no-dominant remodeling, where no clear dominant isoform exists in either group but the isoform distribution changes strongly; and Class F, stable or not high-confidence remodeled genes. Most genes were classified as Class F (i.e., stable with no major isoform changes) and Class A had the least number of genes (Supplementary Fig. 5b). To identify genes within each class that showed statistical divergence from primary to recurrent tumors, we calculated, for each gene, the isoform fraction of each transcript within the gene, compared the primary and recurrent isoform-fraction distributions using Jensen–Shannon divergence (JSD), and quantified the largest individual isoform usage shift using maximum absolute delta isoform fraction, max |dIF| (see methods). High-confidence remodeling required JSD ≥ 0.1, max |dIF| ≥ 0.2, and support in at least four patients. This model distinguished a small subset of high-confidence remodeled genes from a large background of stable genes (Fig. 4e; Supplementary Fig. 5c–d).

We then extended this model by adding Shannon entropy, which measures how concentrated or diffuse isoform usage is within each gene, providing an additional layer of complexity that allows the capture of complex isoform remodeling events. A positive recurrent-primary entropy shift indicates that recurrent isoform usage becomes more diffuse across multiple isoforms, whereas a negative shift indicates that recurrent isoform usage becomes more focused toward fewer isoforms. The JSD–entropy framework separated genes with high distributional divergence but stable entropy from genes in which recurrence produced either a more diffuse or more focused isoform landscape (Fig. 4f; Supplementary Fig. 5e,f). The JSD–entropy quadrant analysis showed that remodeling classes differ not only in whether their isoform distributions diverge, but also in how recurrence changes isoform complexity (Fig. 4f). For example, many Class A and Class B genes remained in low-JSD bins, indicating that dominant isoform identity or secondary isoform structure can change without producing the largest global distributional divergence. In contrast, Classes C, D, and E contained larger fractions of high-JSD genes, consistent with stronger remodeling when dominant isoforms are lost, gained, or absent. Entropy further separated high-JSD genes into cases where recurrence produced a more diffuse isoform landscape, a more focused isoform landscape, or a large usage shift with relatively stable complexity. Thus, the JSD–dIF–entropy model captures three related but distinct dimensions of isoform remodeling: distributional divergence, largest isoform-fraction shift, and change in isoform-usage complexity.

Genes with high JSD (JSD ≥ 0.2) were enriched for biological processes related to extracellular matrix organization, extracellular structure organization, cilium movement, cilium-dependent motility, and microtubule/axoneme-associated processes, suggesting that strong isoform redistribution is not random but concentrates in genes linked to structural and motility-associated biology (Supplementary Fig. 5g). Representative isoform-usage plots, including RBM10, CFAP410, JUN, and EWSR1, illustrate how this combined framework captures several remodeling modes, including dominant switching, secondary isoform redistribution, and recurrence-associated changes in transcript structural category usage (Supplementary Fig. 6a–d).

The second approach used Dirichlet-multinomial modeling with DRIMSeq^16^ to identify genes with statistically significant differential transcript usage while accounting for patient pairing. The distribution of gene-level adjusted P values indicated a subset of genes with significant or near-significant transcript-usage remodeling (Fig. 4g). Enrichment analysis of DTU-significant genes (271 genes with FDR ≤ 0.1) highlighted RNA splicing, mRNA splicing via spliceosome, RNA localization, and related post-transcriptional processes, consistent with recurrence-associated remodeling of RNA-processing programs (Supplementary Fig. 7a).

DTU gene structural-category composition was broadly similar between primary and recurrent tumors, indicating that the most important signal was not a total replacement of transcript annotation classes, but selective redistribution within and across transcript structures (Fig. 4h). At the isoform level, DTU events involved both increased and decreased usage of full-splice match, incomplete-splice match, and novel-in-catalog isoforms, showing that recurrent transcript usage changes were distributed across annotated and partially novel transcript structures (Fig. 4i, Supplementary Fig. 7b). When DTU-significant genes were overlaid onto the Class A–F isoform-remodeling framework, many DTU genes still fell into the stable Class F category, while smaller subsets mapped to Class A, Class C, and Class D remodeling classes, indicating that many apparently stable genes show significant remodeling of their transcript usage (Supplementary Fig. 7c). This distinction is important: Dirichlet-multinomial modeling detects statistically significant transcript-usage shifts, whereas the JSD/dIF/entropy model prioritizes larger effect-size remodeling of gene-level isoform architecture. The two approaches are therefore complementary rather than redundant. Paired isoform trajectory examples, including DELE1 and SLC25A6, showed reciprocal gain and loss of specific isoforms across patients, illustrating how transcript usage remodeling can change dominant isoform architecture even when the same gene remains globally expressed in both primary and recurrent tumors (Supplementary Fig. 7d,e).

Overall, the RNA analysis shows that recurrent astrocytomas undergo transcriptome remodeling at multiple resolutions. Conventional gene-level analysis captures recurrence-associated expression and program shifts, but long-read RNA sequencing reveals a deeper layer of isoform repertoire turnover, dominant and secondary isoform remodeling, entropy-defined changes in transcript usage complexity, and statistically significant differential transcript usage. These data establish that recurrence-associated RNA remodeling is not simply a change in gene abundance. Instead, recurrent tumors reorganize the structure and usage of expressed transcripts, providing the substrate for the ORF/protein-fate analysis and DNA-RNA integration analyses that follow.

### Isoform remodeling is linked to distinct ORF/protein consequences not captured by traditional sequencing approaches

Having established that recurrent astrocytomas undergo extensive isoform remodeling, we next asked whether these transcript changes were predicted to alter coding potential or protein-output state. We used GeneMark-hmm to predict open reading frames (ORFs) from support-filtered Iso-Seq transcripts^17^. To do so, we extracted the FL transcript sequences from each sample, and used GeneMark to predict ORFs from each transcript, limiting the prediction to one ORF per transcript for simplification. Next, we “lifted” the ORF coordinates, which were in the transcript space, back to genome coordinates to properly classify the ORFs compared to the Gencode canonical annotation transcripts as well as to annotate consequence features such as nonsense mediated decay (NMD) potential, frameshift potential, or truncation states. As expected, predicted ORF detection was substantially higher among protein-coding transcripts than among noncoding transcripts in both primary and recurrent tumors, supporting the validity of the ORF annotation framework while also showing that a subset of noncoding-classified transcripts carried predicted ORFs (Fig. 5a).

**Figure 5.**
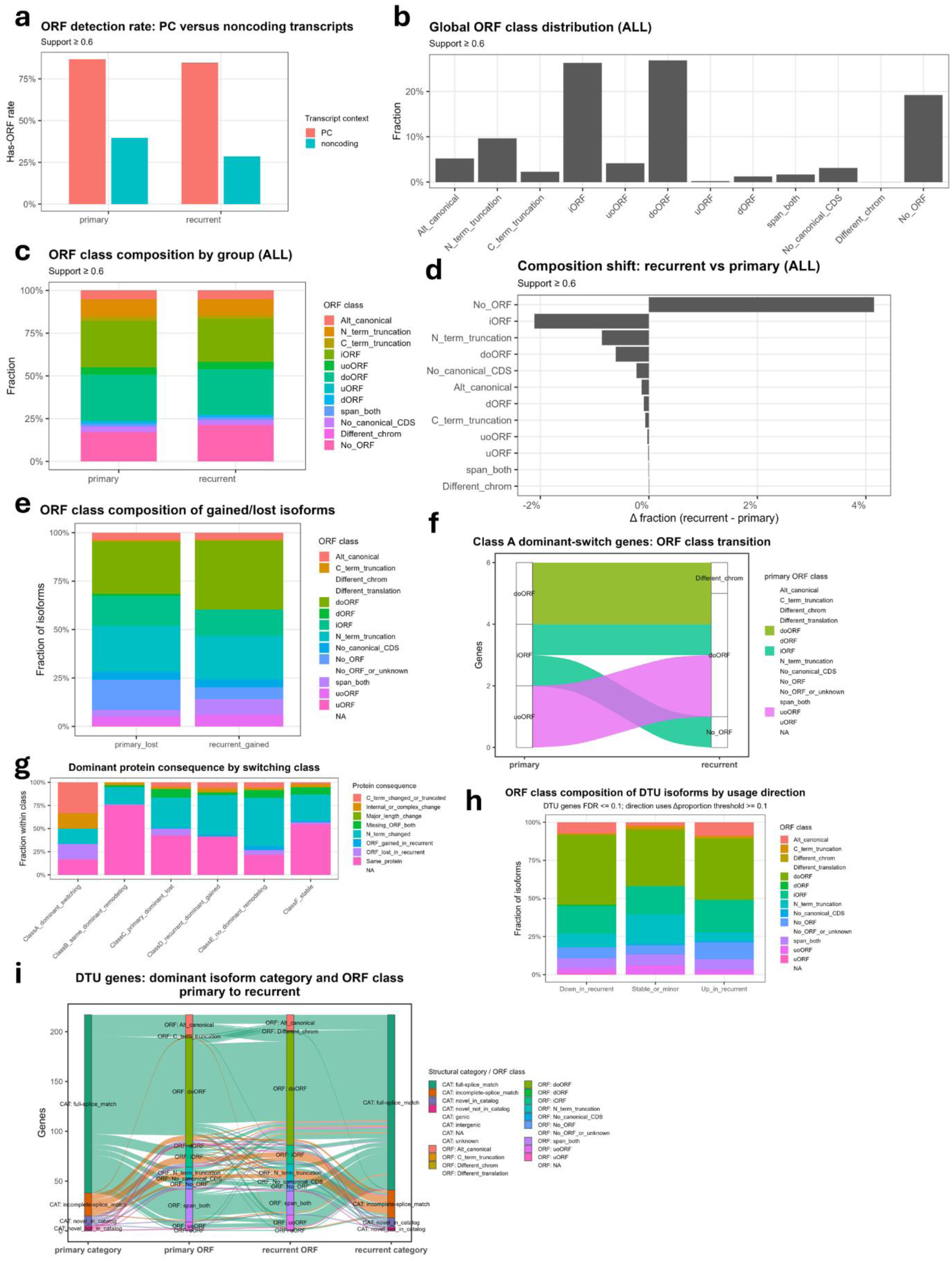
Isoform remodeling is linked to predicted ORF and protein-output consequences during astrocytoma recurrence. **a,** Predicted ORF detection rate among protein-coding and noncoding transcript contexts in primary and recurrent tumors. **b,** Global distribution of predicted ORF classes across all support-filtered transcripts. ORF classes include canonical or alternative canonical ORFs, N- or C-terminal truncation-like ORFs, internal ORFs, upstream or downstream ORF configurations, transcripts without canonical CDS annotation, and transcripts without a predicted ORF. **c,** Predicted ORF-class composition in primary and recurrent tumors across all support-filtered transcripts. **d,** Recurrent-primary shift in global ORF-class fraction across all support-filtered transcripts. Positive values indicate relative enrichment in recurrent tumors; negative values indicate relative depletion. **e,** ORF-class composition of high-confidence primary-lost and recurrent-gained isoforms. **f,** Predicted ORF-class transitions among Class A dominant-switch genes, comparing the dominant isoform ORF class in primary and recurrent tumors. **g,** Dominant predicted protein-output consequence across A–F isoform remodeling classes. Consequences include same predicted protein, ORF gain or loss in recurrence, N- or C-terminal change, internal or complex change, major length change, and missing ORF in both groups. **h,** ORF-class composition of DTU-associated isoforms stratified by recurrent usage direction. **i,** Alluvial plot linking dominant transcript structural category and dominant predicted ORF class in DTU genes from primary to recurrent tumors.

Across all support-filtered isoforms, the ORF landscape was diverse and included canonical or alternative canonical ORFs, N- or C-terminal truncation-like ORFs, internal ORFs, upstream or downstream ORF configurations, transcripts without canonical CDS annotation, and transcripts with no predicted ORF (Fig. 5b). The most abundant classes were iORF and doORF followed by N-terminal truncated ORFs and canonical ORFs (Fig. 5b). The overall ORF-class composition was broadly similar between primary and recurrent tumors, indicating that recurrence did not produce a wholesale shift in the global coding-class composition of all detected transcripts/ORFs (Fig. 5c). Nevertheless, direct recurrent-primary comparison revealed measurable class-level shifts, including an increase in transcripts without predicted ORFs and decreases in selected ORF classes such as internal ORFs and N-terminal truncation-like classes in recurrent tumors (Fig. 5d). Patient-stratified ORF-class composition showed that this global pattern was not driven by a single outlier sample, although patient-level differences in class proportions were present (Supplementary Fig. 8a). Transcript length and exon count distributions differed across ORF classes (Supplementary Fig. 8b-c). Additionally, frameshifting and NMD rates were enriched in biologically plausible ORF categories (For example; Frameshifting was not observed in canonical ORFs but was observed in iORF, uoORF, doORF, and N-terminal truncated ORFs), supporting the internal consistency of the predicted ORF annotations (Supplementary Fig. 8d–e).

We then asked whether the isoforms gained or lost during recurrence were associated with distinct ORF states. High-confidence primary-lost and recurrent-gained isoforms both spanned multiple predicted ORF classes, showing that recurrence-associated isoform turnover affects transcripts with a range of predicted coding outputs rather than only noncoding or low-confidence transcript models (Fig. 5e). ORF-class enrichment analysis further showed that recurrently gained and primary-lost isoforms differed in their relative enrichment for selected ORF classes, indicating that isoform turnover changes the predicted coding-state composition of the expressed transcript repertoire (Supplementary Fig. 9a). For example, ORFs spanning both start and stop codons into untranslated regions (span-both) were more frequently gained in recurrent tumors as well as doORFs and uoORFs. These findings extend the gain/loss analysis from Figure 4 by showing that recurrent isoform turnover is not only structural, but also potentially coding-relevant.

Next, we focused on genes with dominant isoform switching, because these provide the clearest examples in which the main expressed transcript of a gene changes between primary and recurrent tumors. Among Class A dominant-switch genes, primary and recurrent dominant isoforms frequently differed in predicted ORF class, including transitions among internal ORF, downstream-overlapping ORF, upstream-overlapping ORF, no-ORF, and other noncanonical ORF states (Fig. 5f). We then generalized this analysis across the A–F isoform-remodeling classes by assigning dominant protein-output consequences. Although many genes retained the same predicted protein-output category, remodeled classes contained substantial fractions of genes with predicted ORF gain, ORF loss, N-terminal change, C-terminal change or truncation, internal or complex change, and major length change (Fig. 5g). This shows that the A–F remodeling classes are not merely transcript-label changes; rather, they correspond to predicted changes in coding architecture and candidate proteoform output.

We also connected ORF consequences to the independent differential transcript usage analysis. Among DTU-associated isoforms, ORF-class composition differed by usage direction, indicating that isoforms increased, decreased, or only minimally changed in recurrence carried different predicted ORF-state distributions (Fig. 5h). DTU-associated genes also showed linked transitions between dominant transcript structural category and dominant ORF class from primary to recurrence, demonstrating that transcript-usage remodeling can couple structural isoform changes to predicted coding-state changes (Fig. 5i). Top ORF-class switches among DTU isoforms showed that recurrent up- and down-shifted isoforms were not symmetric; different ORF transitions contributed to increased versus decreased usage in recurrent tumors (Supplementary Fig. 9b). Stratifying DTU isoforms by both Pigeon structural category and ORF class further showed that full-splice match, incomplete-splice match, and novel-in-catalog isoforms all contributed to predicted ORF remodeling, rather than a single transcript annotation class dominating the result (Supplementary Fig. 9c). Nonetheless, as most significant DTU changes were annotated as full-splice matches, this class remained the most abundant and carried the most changes in ORF remodeling. Importantly, this indicates that even with stable isoform architecture in remodeled genes, the remodeling potentially leads to translational and proteome consequences.

Overall, these analyses show that recurrence-associated isoform remodeling has predicted coding consequences at several levels. Globally, primary and recurrent tumors retain broadly similar ORF-class composition, but specific gained/lost isoforms, dominant isoform switches, and DTU events are associated with changes in predicted ORF class, NMD or frameshift potential, and protein-output category. Thus, long-read RNA sequencing provides a transcript-to-ORF bridge that is not available from gene-level expression alone. These predicted ORF consequences do not replace proteomic validation, but they provide a principled framework for prioritizing isoform remodeling events most likely to affect protein output and for integrating RNA remodeling with DNA alterations in the following sections.

### An integrative framework maps DNA remodeling to RNA and predicted ORF consequences

Having defined DNA remodeling, RNA isoform remodeling, and predicted ORF consequences separately, we next asked how often these layers could be connected within patient-matched primary and recurrent tumors. Because many DNA alterations occur in genes that are not detected or not quantifiable in long-read RNA, we constructed a denominator-aware framework that separated the global DNA event universe from progressively RNA-aware subsets.

At the DNA level, copy-number alterations represented the largest event family, followed by DMR/methylation events, SV-linked RNA-architecture support, SV/rearrangement events, SNV/indel events, compound cis events, and multi-layer DNA events (Fig. 6a). At the RNA level, the recurrent RNA alteration universe was dominated by gene-expression changes and isoform-usage remodeling, followed by predicted ORF/protein-fate consequences, expressed somatic ALT RNA, differential transcript usage, SV-linked RNA architecture, allele-specific expression or allele-specific transcript usage (ASE/ASTU), and isoform-specific mutant expression (Fig. 6b). Methylation–RNA coupling was tracked separately from the core RNA alteration denominator to avoid counting prior DNA–RNA support itself as an independent RNA alteration class.

**Figure 6.**
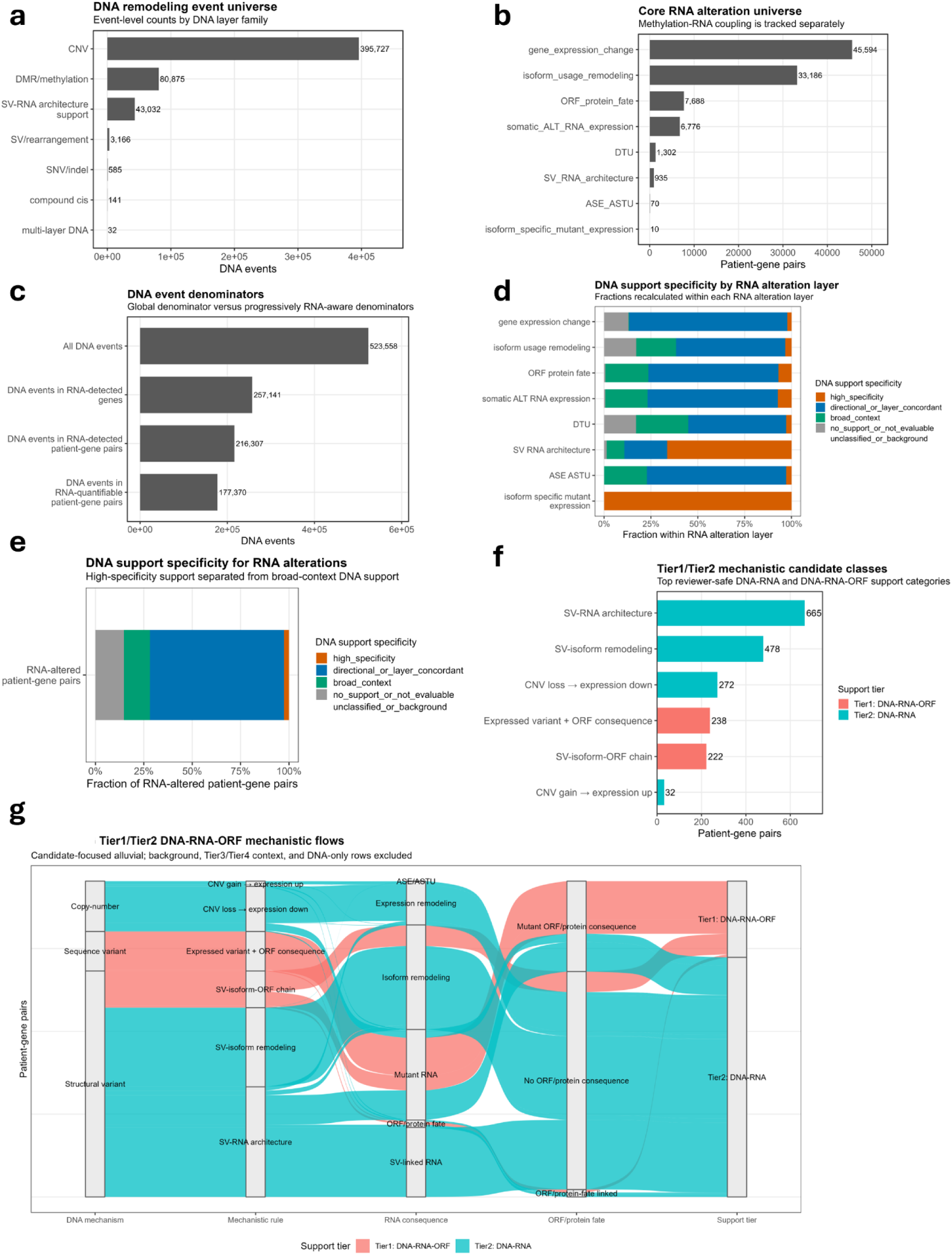
Denominator-aware integration of DNA remodeling with RNA and predicted ORF consequences. **a,** DNA remodeling event universe summarized by DNA layer family. DNA event families include CNV, DMR/methylation, SV-linked RNA-architecture support, SV/rearrangement, SNV/indel, compound cis events, and multi-layer DNA events. **b,** Core RNA alteration universe summarized by RNA alteration layer. RNA layers include gene-expression change, isoform-usage remodeling, predicted ORF/protein-fate consequence, expressed somatic ALT RNA, differential transcript usage, SV-linked RNA architecture, ASE/ASTU, and isoform-specific mutant expression. Methylation–RNA coupling was tracked separately as prior DNA–RNA support rather than counted as a core RNA alteration class. **c,** DNA event denominators used for RNA-aware integration. Bars show the global DNA event universe and progressively stricter RNA-aware subsets: DNA events in RNA-detected genes, DNA events in RNA-detected patient-gene pairs, and DNA events in RNA-quantifiable patient-gene pairs. **d,** DNA support specificity by RNA alteration layer. Fractions are recalculated within each RNA alteration layer and stratified as high-specificity DNA–RNA/ORF support, directional/layer-concordant DNA support, broad DNA context, or no support/not DNA-evaluable. **e,** Global DNA support specificity among RNA-altered patient-gene pairs, separating high-specificity support from directional/layer-concordant support, broad DNA context, and no support/not DNA-evaluable categories. **f,** High-confidence mechanistic candidate classes, showing the top DNA–RNA and DNA–RNA–ORF support categories. Categories include SV-linked RNA architecture, SV-linked isoform remodeling, CNV loss with decreased expression, expressed variants with predicted ORF consequences, SV-linked isoform-to-ORF chains, and CNV gain with increased expression. **g,** Candidate-focused alluvial plot linking DNA mechanism, mechanistic rule, RNA consequence, predicted ORF/protein-fate category, and support tier among high-confidence DNA–RNA and DNA–RNA–ORF chains. Background rows, broader context rows, and DNA-only rows are excluded to emphasize the most interpretable connected events.

We then quantified how the DNA event denominator changed as RNA-awareness became stricter. The broadest denominator contained all DNA events. This was reduced first to DNA events occurring in genes detected by long-read RNA in the cohort, then to DNA events occurring in patient-matched RNA-detected patient-gene pairs, and finally to DNA events in RNA-quantifiable patient-gene pairs (Fig. 6c). Here, a patient-gene pair means that a DNA event and RNA measurement are considered in the same patient and gene context, rather than only at the gene level across the cohort. The stricter RNA-quantifiable denominator further requires sufficient RNA support in that patient-gene context to evaluate RNA remodeling, so it excludes genes that are merely detectable at low or non-informative levels. This denominator structure is important because it prevents silent, non-transcribed, or insufficiently quantified DNA events from diluting DNA-to-RNA interpretation.

Using these denominators, we next estimated how often DNA events were reflected at the RNA and predicted ORF/protein-fate layers. The apparent fraction of DNA events with RNA response, ORF/protein-fate response, or combined RNA plus ORF/protein-fate response depended strongly on the denominator used, with the RNA-quantifiable patient-gene denominator providing the most conservative basis for interpreting DNA-to-RNA reflection (Supplementary Fig. 10a). DNA-layer reflection classes further showed that CNV and DMR/methylation events made up most of the RNA-aware DNA event universe, whereas SV and SNV/indel layers contributed fewer events but more specific DNA–RNA links, such as SV-linked RNA architecture and mutant RNA with predicted ORF/protein consequences (Supplementary Fig. 10b).

We then reversed the direction of analysis and asked how often recurrent RNA remodeling had patient-matched DNA support. DNA support specificity differed substantially by RNA alteration layer (Fig. 6d). Isoform-specific mutant expression and SV-linked RNA architecture showed the strongest high-specificity DNA support, as expected for RNA features that directly encode sequence-variant or structural-variant relationships. Gene-expression change, isoform-usage remodeling, differential transcript usage, ASE/ASTU, and predicted ORF/protein-fate remodeling more often showed directional/layer-concordant or broad DNA-context support rather than direct event-level DNA–RNA evidence (Fig. 6d). Across all RNA-altered patient-gene pairs, most events had directional/layer-concordant DNA support or broad DNA context, whereas high-specificity DNA–RNA/ORF support represented a smaller, more stringent subset (Fig. 6e; Supplementary Fig. 10c). This distinction is important: broad DNA context supports patient-matched genomic co-occurrence, but it should not be interpreted as direct causal evidence.

Finally, we focused on the most interpretable connected events. We used a tiered framework to separate complete DNA–RNA–ORF chains from broader contextual support. Tier 1 represented the strongest class: patient-matched DNA alterations connected to RNA remodeling and a predicted ORF/protein-fate consequence. Tier 2 represented patient-matched, layer-concordant DNA–RNA chains without requiring an ORF/protein-fate consequence. Broader Tier 3 and Tier 4 categories captured patient-matched or gene-level DNA–RNA context but were not treated as high-confidence mechanistic chains (Supplementary Fig. 10d). For the main candidate analysis, we therefore focused on Tier 1 and Tier 2 events.

Within this high-confidence subset, the largest connected classes involved SV-linked RNA architecture and SV-linked isoform remodeling, followed by CNV loss associated with decreased expression, expressed variants with predicted ORF consequences, SV-linked isoform-to-ORF chains, and CNV gain associated with increased expression (Fig. 6f). A candidate-focused alluvial map connected DNA mechanism, mechanistic rule, RNA consequence, predicted ORF/protein-fate category, and support tier (Fig. 6g). Structural variants mainly connected to SV-linked RNA architecture and isoform remodeling, sequence variants connected to mutant RNA and predicted mutant ORF/protein consequences, and copy-number changes connected to directionally concordant expression remodeling (Fig. 6g).

Together, these analyses provide a denominator-aware map of how recurrent glioma DNA remodeling is reflected in the long-read transcriptome. The key result is not that every DNA event produces an RNA consequence, nor that every RNA alteration can be assigned a precise DNA cause. Instead, long-read integration separates global DNA remodeling, RNA-evaluable DNA events, RNA alterations with broad DNA context, and a stringent subset of high-specificity DNA–RNA or DNA–RNA–ORF chains. This framework provides the quantitative foundation for the following layer-specific and gene-set-specific integration analyses.

### Copy-number and expressed sequence variants define broad and high-specificity DNA-to-RNA remodeling routes

Having established a denominator-aware integration framework, we next asked how individual DNA layers contributed to RNA and predicted ORF-level remodeling. Across DNA layer families, copy-number changes were the most abundant source of DNA-RNA coupling, whereas focal non-CNV layers, including sequence variants, DMRs, SVs, LOH, and compound cis-event combinations, contributed fewer but often more specific RNA consequences (Fig. 7a). The layer-to-consequence map showed that CNV gain and CNV loss were most frequently associated with expression up- or down-regulation, while non-CNV layers were more frequently linked to isoform/DTU remodeling, dominant isoform changes, or smaller focused sets of expression-linked events (Fig. 7a). This pattern supports a layered interpretation: CNV provides the broadest dosage-associated RNA signal, whereas focal DNA events provide more specific candidate mechanisms for transcript-architecture and coding-output remodeling. We first focused on CNV because it represented the largest DNA event family and the most abundant DNA-to-RNA connection. CNV-linked RNA remodeling was dominated by CNV gain associated with expression increase, followed by CNV loss associated with expression increase, CNV loss associated with expression decrease, and CNV gain associated with expression decrease (Fig. 7b). Thus, while canonical dosage-concordant relationships were common, CNV state did not deterministically predict expression direction in all patient-gene contexts. Patient-level CNV coupling profiles showed that all DNA-evaluable patients carried large numbers of CNV-linked RNA events, with variable proportions of concordant, discordant, and isoform/DTU/ORF-coupled cases (Fig. 7c). A direct comparison of recurrent-primary CNV delta against recurrent-primary RNA expression changes further showed a mixture of concordant, discordant, isoform/ORF-linked, and non-strictly coupled events across patients (Supplementary Fig. 11a). Compound cis-event combinations also mapped to multiple RNA consequence classes, including expression change, isoform/DTU remodeling, and dominant isoform change, suggesting that some RNA effects occur in genomic contexts where more than one DNA layer is altered in the same gene-patient setting (Supplementary Fig. 11b). Together, these analyses indicate that CNV provides a broad RNA-dosage substrate, but additional regulatory and structural layers are needed to explain the full spectrum of RNA remodeling.

**Figure 7.**
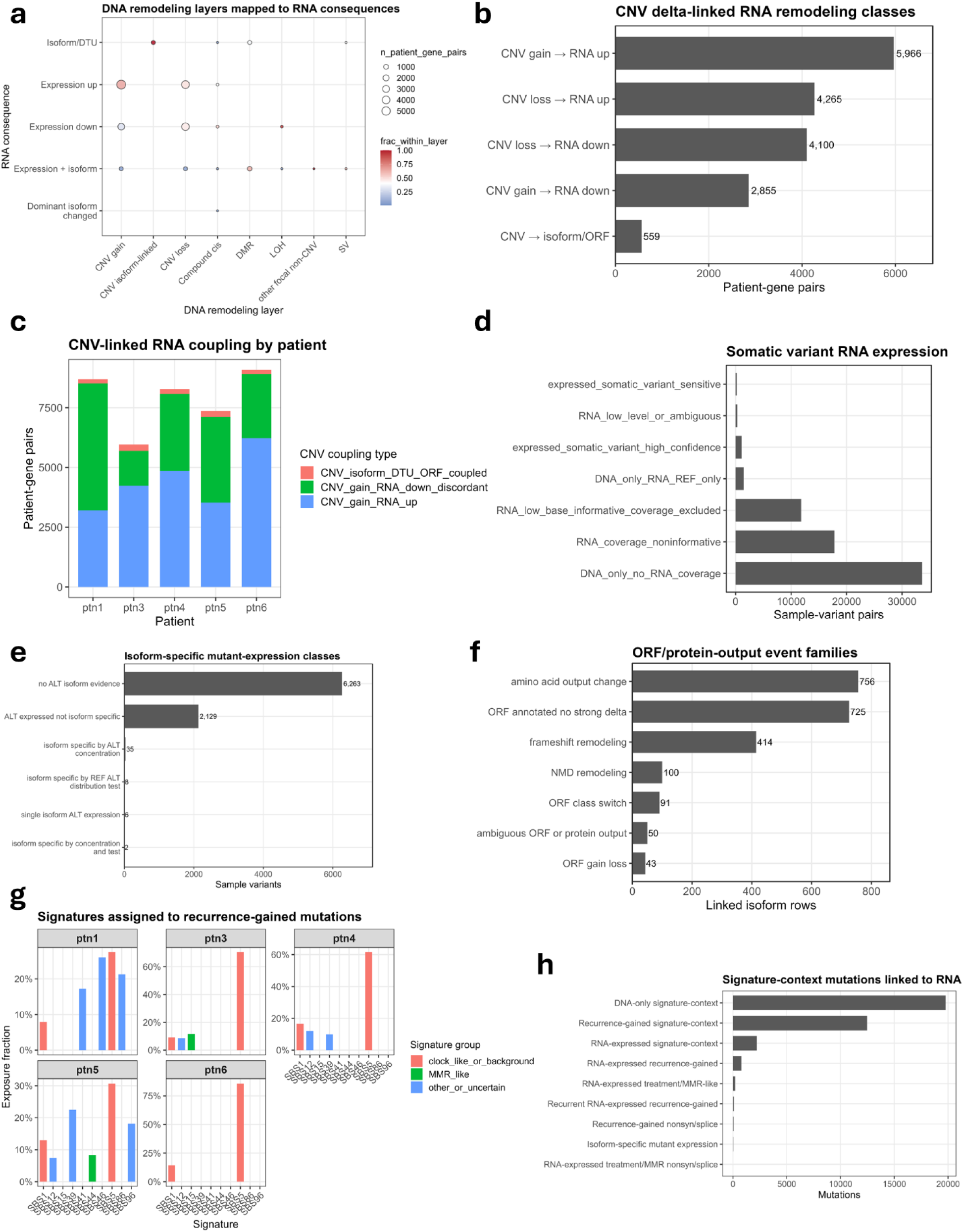
Copy-number and expressed sequence variants connect DNA remodeling to RNA and predicted ORF consequences. **a,** DNA remodeling layers mapped to RNA consequence classes. Point size indicates the number of patient-gene pairs, and color indicates the fraction of events within each DNA layer mapping to the indicated RNA consequence. RNA consequence classes include expression up, expression down, expression plus isoform change, isoform/DTU remodeling, and dominant isoform change. **b,** CNV delta-linked RNA remodeling classes across patient-gene pairs. Categories summarize directional relationships between recurrent-primary CNV change and RNA expression change, including CNV gain with RNA up, CNV loss with RNA up, CNV loss with RNA down, CNV gain with RNA down, and CNV-associated isoform/ORF remodeling. **c,** CNV-linked RNA coupling by patient. Stacked bars summarize patient-level counts of CNV gain/RNA down discordance, CNV gain/RNA up concordance, and CNV-associated isoform/DTU/ORF-coupled events. **d,** Somatic variant RNA-expression classes. DNA variants were stratified according to RNA coverage and ALT RNA evidence, including DNA-only variants without RNA coverage, RNA-covered but non-informative variants, variants excluded for low base-informative RNA coverage, DNA-only/RNA REF-only variants, high-confidence expressed somatic variants, ambiguous low-level RNA evidence, and sensitive expressed-variant calls. **e,** Isoform-specific mutant-expression classes among RNA-informative variants. Categories distinguish variants without ALT isoform evidence from ALT-expressed variants without isoform specificity and smaller subsets with isoform-specific ALT concentration or related isoform-specific mutant-expression patterns. **f,** Predicted ORF/protein-output event families among mutation-linked isoform rows. Events include amino-acid output change, ORF-annotated transcripts without strong ORF delta, transcript remodeling, frameshift remodeling, NMD remodeling, ORF-class switching, ambiguous ORF/protein-output states, and ORF gain or loss. **g,** COSMIC SBS signatures assigned to recurrence-gained mutations by patient. Signature groups are summarized as clock-like/background, MMR-like, or other/uncertain mutational-process contexts. **h,** Signature-context mutations linked to RNA expression. Classes separate DNA-only signature-context mutations from RNA-expressed, recurrence-gained, treatment/MMR-associated, nonsynonymous/splice-linked, and isoform-specific mutant-expression subsets.

We then examined somatic SNVs and indels, which are less frequent than CNV events but can provide more direct molecular links when the mutant allele is expressed. Most DNA variants had no informative RNA evidence because the variant fell in a gene or region without sufficient RNA coverage, or because RNA coverage at the variant base was non-informative (Fig. 7d). Among RNA-informative variants, only a smaller subset showed high-confidence or sensitive evidence of somatic alternative (ALT) RNA expression (Fig. 7d). Nonetheless, the absence of mutant RNA evidence could reflect RNA detectability and base-level coverage constraints rather than true absence of mutant transcript expression. Integrated variant classes further showed that most patient variants remained DNA-only or low RNA-support events, whereas smaller subsets showed variant-associated RNA remodeling, ASE/ASTU support, expressed recurrent variants, or mutant-dominant isoform evidence (Supplementary Fig. 11c). Patient-gene bubble summaries highlighted recurrently altered genes with stronger integrated evidence, including canonical glioma or cancer-associated genes such as IDH1, TP53, ATRX, and CIC-related contexts, but also showed that high-confidence expressed-variant evidence remained selective rather than universal (Supplementary Fig. 11d).

Long-read RNA provided an additional advantage by assigning expressed mutant alleles to isoform context. Most ALT-expressed variants did not show clear isoform-specific enrichment, but a subset showed evidence for ALT expression concentrated in particular isoforms or associated with isoform-specific mutant-expression patterns (Fig. 7e). This distinction is important because mutant RNA expression at the gene level does not necessarily imply that all isoforms carry the mutant allele equally. Mutation-linked isoform rows were then classified by predicted ORF/protein-output event family. The largest groups involved amino-acid output change, ORF-annotated transcripts without a strong ORF delta, transcript remodeling, frameshift remodeling, NMD remodeling, ORF-class switching, ambiguous protein-output states, and ORF gain or loss (Fig. 7f). These results show that expressed somatic variants can extend beyond simple mutant allele detection to candidate isoform-specific and predicted coding-output consequences.

Finally, we asked whether mutational-process context helped organize expressed variants and their RNA consequences. Somatic mutation burden and temporal mutation classes varied across patients, with recurrent-primary burden changes showing patient-specific gains and losses rather than one uniform recurrence pattern (Supplementary Fig. 11e–g). COSMIC SBS signature analysis assigned recurrence-gained mutations to clock-like/background, MMR-like, and other/uncertain signature groups, with different patients showing distinct signature compositions (Fig. 7g; Supplementary Fig. 12a). When signature-context mutations were linked to RNA expression, most remained DNA-only or signature-context events without RNA expression, but smaller subsets were RNA-expressed, recurrence-gained, treatment/MMR-associated, nonsynonymous or splice-linked, or isoform-specific mutant-expression events (Fig. 7h). The most frequent RNA-expressed signature contexts included SBS86, SBS15, SBS5, SBS8, SBS96, and SBS44, and patient-level heatmaps showed that RNA-expressed signature-context mutations were distributed across clock-like/background, MMR-like, and other/uncertain groups (Supplementary Fig. 12b,c). A signature-to-RNA-to-isoform/coding alluvial further showed that only a minority of signature-context mutations propagated into RNA expression, isoform-specific mutant RNA, or predicted coding consequences (Supplementary Fig. 12d). Thus, mutational-process analysis provides context for expressed variants, but the strongest mechanistic interpretation comes from the subset of mutations that can be connected directly to mutant RNA, isoform specificity, and predicted ORF/protein-output consequences.

Overall, these analyses show that DNA-to-RNA coupling is layer-dependent. CNV alterations provide the broadest and most abundant route to RNA remodeling, but their effects are often dosage-associated and sometimes directionally discordant. In contrast, expressed somatic variants and signature-context mutations are less frequent but more mechanistically specific when mutant RNA, isoform-specific ALT expression, and predicted ORF/protein-output consequences can be demonstrated. This separation between broad dosage coupling and focused expressed-variant chains provides the first layer-specific extension of the denominator-aware framework.

### Allele- and haplotype-specific transcript usage reveals recurrence-associated isoform remodeling

We next asked whether recurrence-associated RNA remodeling could be resolved at the allele or haplotype-marker level. We distinguished two related but different signals. Allele-specific expression (ASE) measures whether one allele contributes more total RNA from a gene, usually summarized as a major-allele fraction across RNA-covered heterozygous markers. In contrast, allele- or haplotype-specific transcript usage (ASTU) asks whether reads assigned to different alleles or haplotype-marker groups use different isoforms within the same gene. Therefore, ASE captures allelic dosage imbalance, whereas ASTU captures allele-resolved isoform choice. Importantly, ASTU can occur with or without strong gene-level ASE.

We first evaluated recurrent ASE and its DNA support. RNA-covered heterozygous SNPs were abundant across patients and timepoints, providing sufficient marker coverage for allele-aware RNA analysis (Supplementary Fig. 13a). Most high-confidence recurrent ASE candidates were supported by LOH or allele-specific copy-number evidence, whereas RNA-only ASE candidates without patient-specific DNA support represented a much smaller class (Fig. 8a; Supplementary Fig. 13b). Gene-level major-allele fraction distributions showed broad allelic imbalance spectra across patients, with a subset of genes approaching strong monoallelic or near-monoallelic RNA contribution (Supplementary Fig. 13c). These findings indicate that recurrent ASE is frequently embedded in DNA allelic architecture rather than being only a transcript-level imbalance.

**Figure 8.**
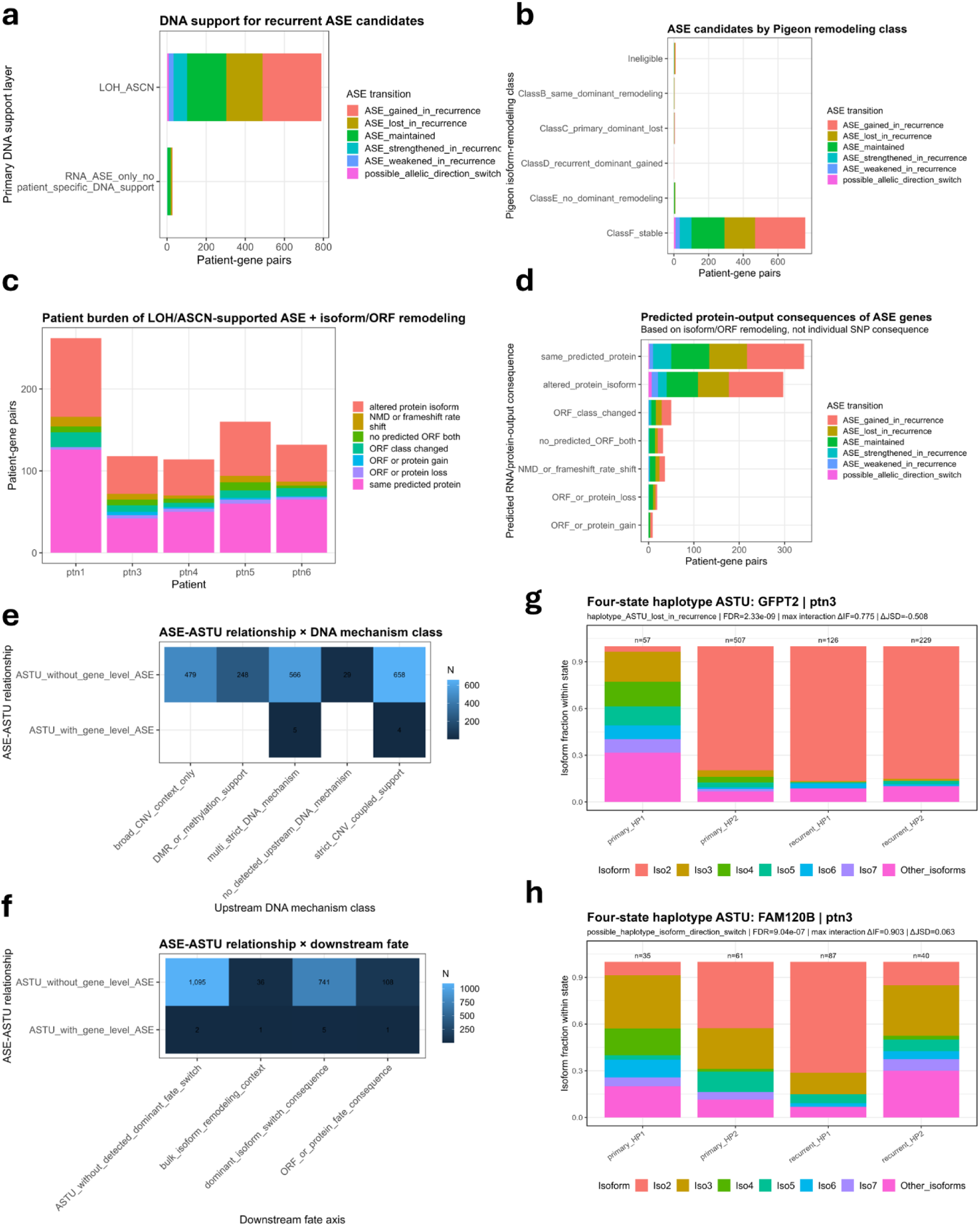
Allele-specific expression and allele/haplotype-specific transcript usage reveal recurrence-associated RNA remodeling. **a,** DNA support for high-confidence recurrent ASE candidates. Bars stratify recurrent ASE transitions by primary DNA support layer, including LOH/allele-specific copy-number support and RNA-only ASE without patient-specific DNA support. **b,** LOH/allele-specific CNV-supported ASE candidates stratified by bulk Pigeon isoform-remodeling class. Class F indicates genes stable or not high-confidence remodeled in the bulk A–F isoform framework. **c,** Patient burden of LOH/allele-specific CNV-supported ASE genes with isoform or predicted ORF/protein-output remodeling. Bars summarize predicted consequence classes per patient. **d,** Predicted protein-output consequences of LOH/allele-specific CNV-supported ASE genes, stratified by ASE transition class. Consequence classes include same predicted protein, altered protein isoform, ORF-class change, ORF/protein gain or loss, no predicted ORF in both groups, and NMD or frameshift-rate shift. **e,** Relationship between ASTU and gene-level ASE across upstream DNA mechanism classes. Counts are stratified by whether ASTU occurred with or without detectable gene-level ASE and by DNA mechanism context, including broad CNV context, DMR/methylation support, multi-strict DNA mechanism, no detected upstream DNA mechanism, and strict CNV-coupled support. **f,** Relationship between ASTU and gene-level ASE across downstream fate axes. Counts are stratified by whether ASTU occurred with or without detectable gene-level ASE and by downstream fate category, including ASTU without detected dominant fate switch, bulk isoform-remodeling context, dominant isoform-switch consequence, and predicted ORF/protein-fate consequence. **g,** Four-state haplotype-marker ASTU example for GFPT2 in patient 3. Bars show isoform fractions for HP1 and HP2 marker-assigned reads in primary and recurrent tumors, illustrating loss of haplotype-specific transcript usage in recurrence. **h,** Four-state haplotype-marker ASTU example for FAM120B in patient 3. Bars show isoform fractions for HP1 and HP2 marker-assigned reads in primary and recurrent tumors, illustrating a possible recurrence-associated direction switch in haplotype-specific isoform preference.

We then asked whether DNA-supported ASE coincided with isoform and predicted ORF remodeling. LOH/allele-specific CNV-supported ASE candidates were dominated by genes classified as bulk-stable in the A–F Pigeon remodeling framework, but smaller subsets overlapped dominant or secondary isoform-remodeling classes (Fig. 8b). This indicates that allele-specific imbalance can occur in genes that do not show a major bulk isoform-class switch, meaning that allele-aware analysis can uncover remodeling hidden from aggregate RNA measurements. At the patient level, LOH/allele-specific CNV-supported ASE genes showed variable burdens of predicted protein-output consequences, including same predicted protein, altered protein isoform, ORF-class change, ORF/protein gain or loss, and NMD or frameshift-rate shifts (Fig. 8c,d). Extended ASE summaries similarly showed that DNA-supported ASE candidates were enriched for isoform/ORF remodeling, while patient-specific recurrent ASE was most often explained by LOH or allele-specific copy-number imbalance, with smaller contributions from multi-layer DNA explanations (Supplementary Fig. 13d,e). Top recurrent ASE candidates showed both ASE gained in recurrence and ASE strengthened in recurrence classes, with recurrent major-allele fractions reaching high values in selected genes (Supplementary Fig. 13f).

We next moved from total allelic expression to allele- or haplotype-marker-resolved isoform usage. Read-level HP1/HP2 assignment confirmed that large numbers of full-length reads could be assigned to haplotype-marker groups across samples, enabling isoform-fraction comparisons between haplotypes (Supplementary Fig. 14a). ASTU effect-size analysis compared two dimensions: Jensen–Shannon divergence between HP1 and HP2 isoform distributions, which measures how different the haplotype-specific isoform repertoires are, and the maximum absolute isoform-fraction shift, which captures the largest isoform-level difference between haplotypes (Supplementary Fig. 14b). Comparing primary and recurrent HP1-versus-HP2 JSD further showed that haplotype-specific isoform divergence could be maintained, gained, lost, weakened, or strengthened across recurrence (Supplementary Fig. 14c). Thus, ASTU provides an isoform-resolution view of allele/haplotype remodeling that is not captured by gene-level ASE alone.

We then integrated ASTU with upstream DNA mechanism classes and downstream RNA/ORF fate axes. Recurrence-associated ASTU candidates were observed across all DNA-evaluable patients and were associated with broad CNV context, DMR/methylation support, LOH or allele-specific CNV, strict CNV-coupled support, and SV/fusion contexts (Supplementary Fig. 14d). Downstream, ASTU candidates overlapped bulk isoform-remodeling context, dominant isoform-switch consequences, and predicted ORF/protein-fate consequences, although many remained in an ASTU-only class without detected dominant fate switching (Supplementary Fig. 14e). When ASTU was cross-tabulated with gene-level ASE, most ASTU events occurred without detectable gene-level ASE, whereas a smaller subset showed both ASTU and gene-level ASE (Fig. 8e,f). Thus, allele-specific isoform usage is distinct from allele-specific expression, and it can reveal allele-resolved transcript architecture even when total RNA output from the two alleles is not strongly imbalanced.

Finally, we examined how ASTU related to bulk Pigeon isoform-remodeling classes. Most recurrence-ASTU candidates occurred in genes classified as Class F/stable in bulk isoform analysis, with smaller contributions from dominant switching, same-dominant remodeling, primary-dominant loss, recurrent-dominant gain, and no-dominant remodeling classes (Supplementary Fig. 14f–h). This shows that marker-resolved isoform usage can expose a hidden layer of remodeling that is averaged out in bulk gene-level or isoform-class analyses. Four-state examples in patient 3 illustrated this behavior. In GFPT2, haplotype-specific isoform usage was lost in recurrence, with recurrent HP1 and HP2 converging toward a more similar isoform composition (Fig. 8g). In FAM120B, the primary and recurrent haplotype-marker groups showed a possible direction switch in isoform preference, demonstrating that recurrence can alter not only the strength but also the direction of allele/haplotype-specific isoform usage (Fig. 8h).

Together, these analyses show that recurrence-associated transcript remodeling includes an allele-aware layer that is not captured by gene-level expression, bulk isoform usage, or ORF analyses alone. ASE identifies genes in which one allele contributes more total RNA, often in the setting of LOH or allele-specific copy-number imbalance. ASTU identifies genes in which allele- or haplotype-marker groups use different isoforms, frequently without detectable gene-level ASE. This distinction supports a model in which recurrent astrocytomas remodel not only RNA abundance and isoform composition, but also the allele-specific architecture of transcript usage.

### DNA methylation remodeling links regulatory DNA changes to isoform and predicted ORF consequences

We next asked whether recurrence-associated DNA methylation changes were connected to RNA architecture and predicted coding consequences. Differentially methylated regions (DMRs) were first stratified by regulatory context and temporal status. Across UTR, promoter, intronic, gene-annotated, and exonic DMR classes, many patient-gene DMR relationships were shared between primary and recurrent tumors, but both primary-only and recurrent-only DMRs were also detected across regulatory categories (Fig. 9a). DMR directionality showed that hypermethylated and hypomethylated DMRs were distributed across regulatory contexts, with intronic and gene-annotated DMRs forming the largest absolute classes (Supplementary Fig. 15a). The recurrent-primary methylation effect distribution was broad and centered near zero, with positive and negative tails, indicating that most DMR-associated patient-gene contexts showed modest shifts while a smaller subset showed stronger recurrent hypermethylation or hypomethylation (Supplementary Fig. 15b).

**Figure 9.**
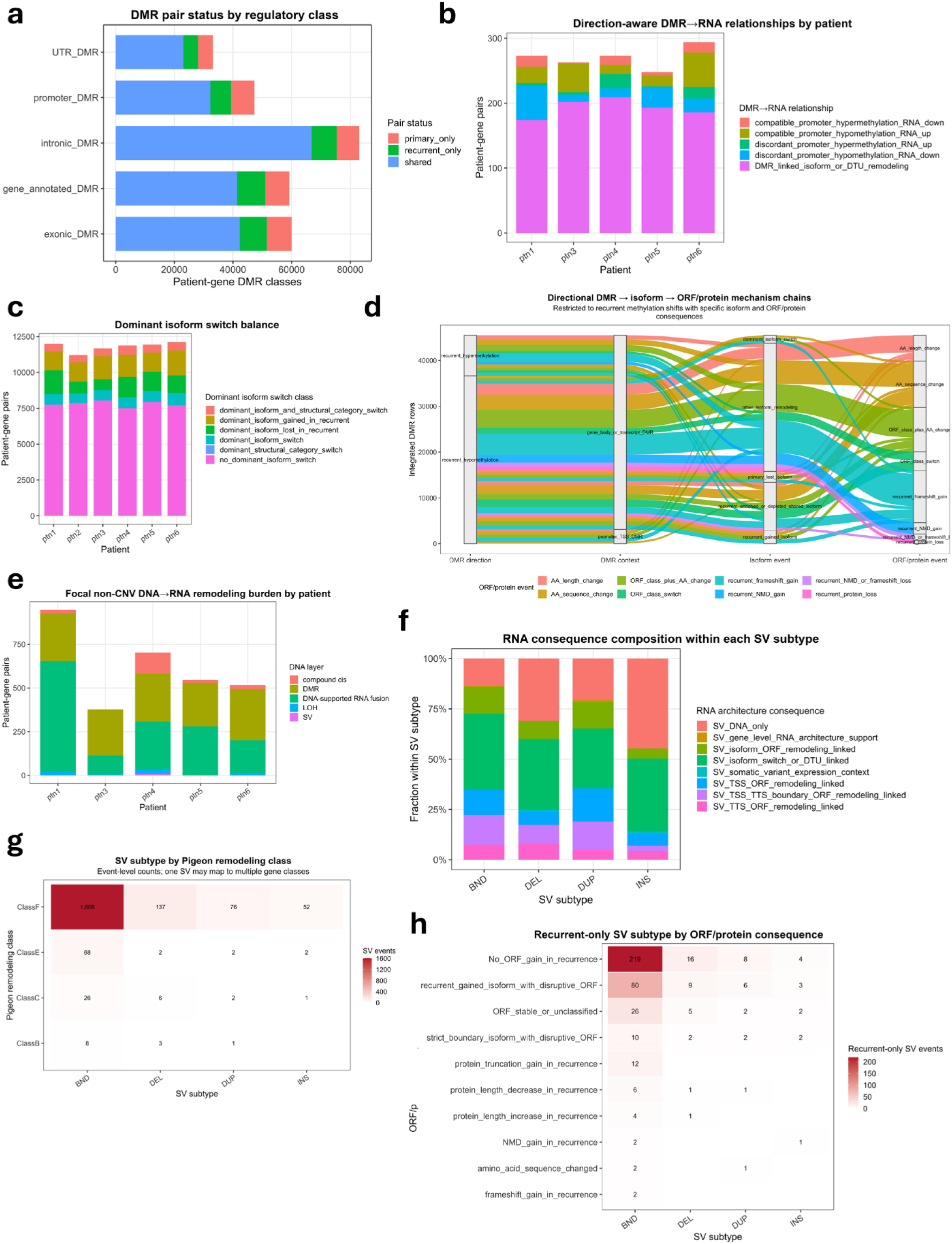
DMRs and structural variants connect focal non-CNV DNA remodeling to RNA architecture and predicted ORF consequences. **a,** DMR pair status by regulatory class. Patient-gene DMR classes are stratified by genomic context, including UTR, promoter, intronic, gene-annotated, and exonic DMRs, and by temporal status as shared, primary-only, or recurrent-only. **b,** Direction-aware DMR-RNA relationships by patient. Patient-gene pairs are classified by whether DMR direction and RNA change are compatible with promoter hypermethylation/RNA down, promoter hypomethylation/RNA up, discordant promoter relationships, isoform or UTR remodeling, or other DMR-linked contexts. **c,** Dominant isoform switch classes among DMR-linked patient-gene pairs. Bars summarize whether genes show no dominant isoform switch, dominant structural-category switch, recurrent dominant isoform gain, primary dominant isoform loss, or dominant isoform switching. **d,** Directional DMR-to-isoform-to-predicted ORF/protein alluvial. Flows connect recurrent methylation direction, DMR context, isoform event class, and predicted ORF/protein event class among DMR-linked rows with specific isoform and ORF/protein consequence annotations. **e,** Focal non-CNV DNA-RNA remodeling burden by patient. Stacked bars summarize patient-level counts across focal DNA layers, including compound cis events, DMRs, DNA-supported RNA fusions, LOH, and SVs. **f,** RNA consequence composition within each SV subtype. Fractions are shown separately for breakend-like, deletion, duplication, and insertion SV classes and stratified by RNA consequence class. **g,** SV subtype by Pigeon bulk isoform-remodeling class. Heatmap values indicate SV event counts across SV subtype and A–F Pigeon remodeling classes. **h,** Recurrent-only SV subtype by predicted ORF/protein consequence. Heatmap values indicate recurrent-only SV-linked event counts across SV subtype and ORF/protein consequence class.

We then connected DMRs to gene expression and isoform remodeling. Promoter/TSS methylation changes showed both concordant and discordant relationships with gene expression, rather than a simple one-directional methylation-expression pattern across all genes (Supplementary Fig. 15c). We therefore treated methylation-RNA relationships as regulatory associations that require genomic and transcript context, not as direct proof of repression or activation. Direction-aware DMR-RNA classification showed that most DMR-linked patient-gene pairs were associated with isoform or UTR remodeling rather than simple promoter methylation-expression concordance, although compatible promoter hypermethylation/RNA down and promoter hypomethylation/RNA up relationships were present in subsets of cases (Fig. 9b). Extended coupling analysis similarly showed that many DMRs had RNA architecture context, while smaller subsets linked gene-body DMRs to ORF consequences, isoform switching, or DTU-like remodeling (Supplementary Fig. 15d).

Because long-read RNA captures complete isoform structures, we next asked whether DMR-associated RNA remodeling reached dominant isoform and predicted ORF/protein-fate levels. DMR-linked dominant isoform analysis showed that most gene-patient pairs did not undergo a dominant isoform switch, but subsets showed dominant structural-category switching, recurrent dominant isoform gain, primary dominant isoform loss, or dominant isoform switching (Fig. 9c). Multilayer and confounder-aware methylation summaries showed that many methylation-RNA candidates occurred without detected LOH/CNV context, while a smaller subset occurred together with LOH/CNV, SV, or haplotype-context support (Supplementary Fig. 15e). Directional DMR-to-isoform-to-ORF alluvial analysis further connected recurrent hypermethylation and hypomethylation, promoter/TSS and gene-body/transcript DMR contexts, isoform events, and predicted ORF/protein consequences (Fig. 9d). In this restricted set of complete DMR–isoform–ORF chains, gene-body or transcript-associated DMRs contributed prominently to coding-impact classes, including protein remodeling without simple gain or loss and ORF/protein gain or loss categories, whereas promoter/TSS DMRs contributed a smaller but more directly interpretable expression-linked component (Fig. 9d; Supplementary Fig. 15f,g). These results indicate that methylation remodeling in recurrent astrocytoma is not only a promoter-expression phenomenon. A subset of DMRs is linked to isoform architecture and predicted coding-output changes, especially when DMRs occur in gene-body or transcript-associated contexts.

### Structural variants link focal non-CNV DNA remodeling to RNA architecture and predicted coding consequences

We next examined focal non-CNV DNA alterations, focusing on structural variants because they provide a direct architectural route from DNA rearrangement to RNA structure. Across patients, focal non-CNV DNA-RNA remodeling included DMRs, DNA-supported RNA fusions, SVs, LOH, and compound cis events, with different patients showing different dominant non-CNV layers (Fig. 9e). This patient-level heterogeneity indicates that focal DNA-to-RNA remodeling is not driven by one universal event type, but by patient-specific combinations of regulatory, allelic, and structural mechanisms.

SVs were then analyzed in more detail. Most detected SVs were primary-only or recurrent-only rather than shared, indicating substantial temporal remodeling of the structural-variant landscape (Supplementary Fig. 16a). Breakend-like events represented the largest SV class, followed by deletions, insertions, and duplications (Supplementary Fig. 16b). Breakpoints were enriched in intragenic and near-gene contexts, with additional promoter-proximal, TSS-proximal, and TTS-proximal subsets (Supplementary Fig. 16c). Many SV breakpoints were located within several kilobases of linked isoform boundaries, and breakpoint-to-isoform-boundary links were particularly common near novel isoform TSS/TTS positions, observed isoform boundaries, and recurrently gained isoform boundaries (Supplementary Fig. 16d,e). These patterns support a structural model in which SVs can remodel RNA architecture by altering, disrupting, or lying near transcript boundary regions.

We next asked whether SVs were supported by RNA junction evidence and whether they mapped to specific RNA consequence classes. A substantial fraction of recurrent and recurrence-gained SV breakpoint-gene links showed gene-level RNA junction support, although this should not be interpreted as base-pair-resolution fusion validation for every event (Supplementary Fig. 16f). RNA consequence composition differed by SV subtype, with breakend-like, deletion, duplication, and insertion events showing different fractions of SV-DNA-only calls, SV-linked gene-level RNA architecture support, isoform remodeling, somatic-variant expression context, and TSS/TTS or ORF remodeling classes (Fig. 9f). At the transcript-annotation level, most SV-linked events mapped to bulk-stable Class F genes, but recurrent SVs also intersected genes with Class E, Class C, and Class B remodeling patterns, indicating that SV-associated RNA effects can occur both with and without a large bulk isoform-class transition (Fig. 9g; Supplementary Fig. 16g). This mirrors the allele/haplotype-specific transcript-usage findings: focal DNA events can affect transcript architecture in ways that are partly hidden by bulk remodeling classes.

Finally, recurrent-only SVs were connected to predicted ORF/protein consequences. Breakend-like events contributed the largest number of recurrent-only SV-linked ORF/protein consequence rows, partly reflecting their higher event burden, but deletions, duplications, and insertions also contributed to recurrent isoform gain, disruptive ORF gain, protein truncation gain, protein length changes, frameshift gain, NMD gain, and amino-acid sequence change categories (Fig. 9h). SV-linked RNA architecture consequence classes further showed that SVs were associated with isoform switch or DTU-like remodeling, DNA-only SVs, TSS/TTS-boundary ORF remodeling, isoform-ORF remodeling, gene-level RNA architecture support, and somatic-variant expression context (Supplementary Fig. 16h). A top-gene candidate analysis identified genes with recurrent SV breakpoints linked to junction remodeling, with NLGN1 among the strongest candidates (Supplementary Fig. 17a). Representative IGV DNA tracks and RNA sashimi plots for NLGN1 from patient 4 show structural complexity at the NLGN1 locus and the remodeling observed in the recurrent tumor, together with altered primary-to-recurrent junction architecture at the RNA level. (Supplementary Fig. 17b,c). Together, these analyses show that SVs provide one of the clearest focal DNA-to-RNA architectural links in the cohort, with a subset extending to predicted ORF/protein-fate consequences.

### Targeted biological gene sets reveal convergence of DNA–RNA–ORF remodeling on recurrence-relevant programs

After defining layer-specific DNA–RNA mechanisms, we asked whether these events converged on biological gene sets relevant to astrocytoma recurrence. We first examined splicing and RNA-binding protein regulators, which are relevant to the extensive isoform remodeling observed. Within this gene set, integrated DNA–RNA–ORF flows showed that many gene-patient pairs had either CNV-dominant or no clear DNA event context, but a subset connected DNA remodeling to expressed somatic ALT RNA, major dominant-isoform switching, methylation–RNA coupling, SV-linked RNA architecture, and predicted mutant RNA or ORF/protein consequences (Fig. 10a). Consistent with this, paired expression analysis showed recurrent shifts in multiple splicing/RBP regulators, including increases in several RBM, SRSF, SF3, HNRNP, and RNA-processing genes, while others decreased in recurrence (Supplementary Fig. 18a). These results suggest that splicing-regulatory genes are themselves remodeled at DNA and RNA levels, providing a plausible upstream context for the widespread isoform remodeling observed across the cohort.

**Figure 10.**
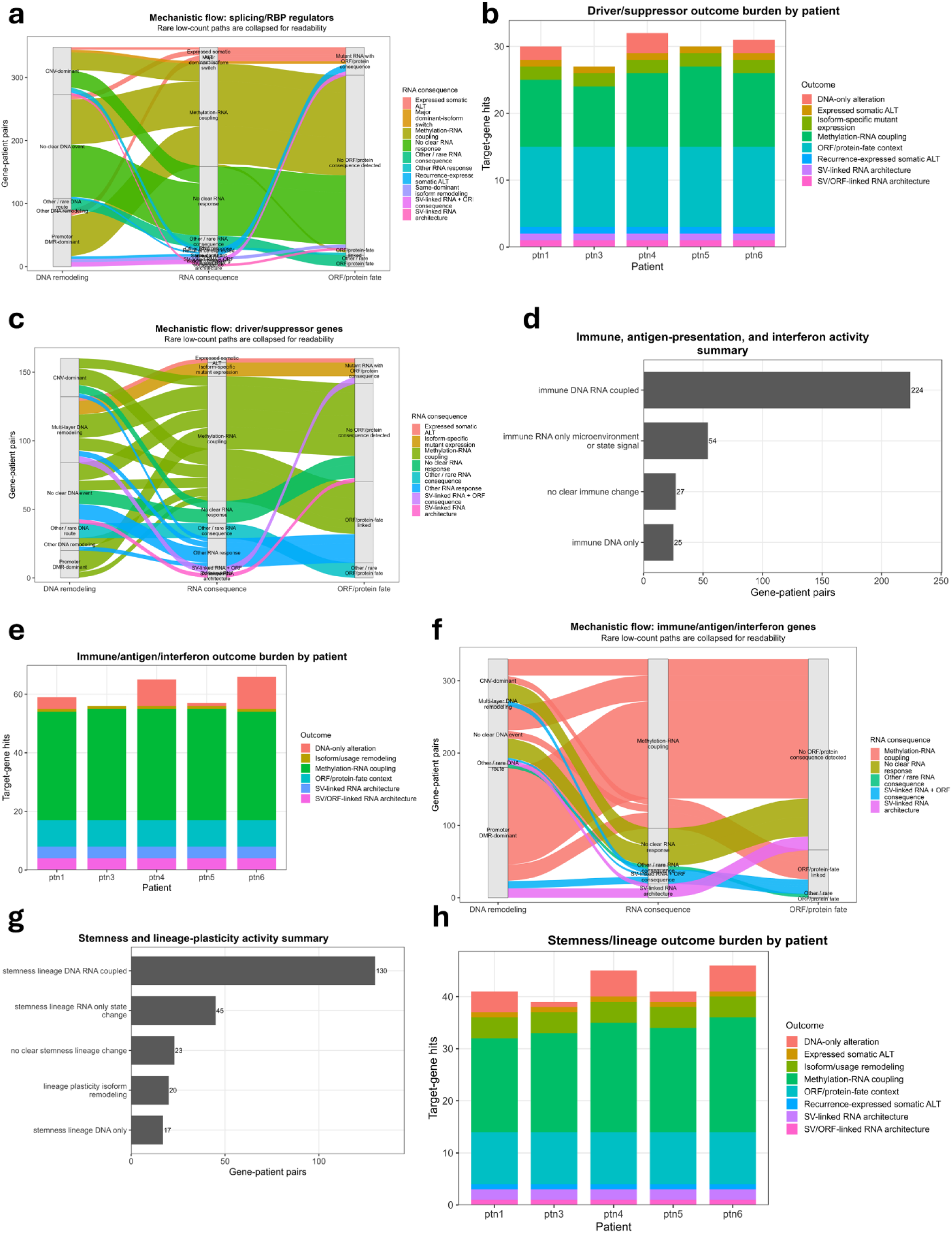
Targeted biological gene-set integration reveals convergence of DNA–RNA–ORF remodeling on recurrence-relevant programs. **a,** Collapsed mechanistic flow for splicing and RNA-binding protein regulator genes. Flows connect DNA remodeling class, RNA consequence class, and predicted ORF/protein-fate class across splicing/RBP gene-patient pairs. Rare low-count paths are collapsed for readability. **b,** Driver and tumor-suppressor outcome burden by patient. Stacked bars summarize target-gene hits across outcome classes, including DNA-only alteration, expressed somatic ALT, isoform-specific mutant expression, methylation–RNA coupling, ORF/protein-fate context, recurrence-expressed somatic ALT, SV-linked RNA architecture, and SV/ORF-linked RNA architecture. **c,** Collapsed mechanistic flow for driver and tumor-suppressor genes. Flows connect DNA remodeling class, RNA consequence class, and predicted ORF/protein-fate class, with rare low-count paths collapsed for readability. **d,** Immune, antigen-presentation, and interferon activity summary. Bars show gene-patient pair counts classified as DNA–RNA coupled, RNA-only microenvironment or state signal, DNA-only alteration, or no clear immune change. **e,** Immune, antigen-presentation, and interferon outcome burden by patient. Stacked bars summarize outcome classes among immune-related target genes, including DNA-only alteration, isoform/usage remodeling, methylation–RNA coupling, ORF/protein-fate context, SV-linked RNA architecture, and SV/ORF-linked RNA architecture. **f,** Collapsed mechanistic flow for immune, antigen-presentation, and interferon genes. Flows connect DNA remodeling class, RNA consequence class, and predicted ORF/protein-fate class, with rare low-count paths collapsed for readability. **g,** Stemness and lineage-plasticity activity summary. Bars show gene-patient pair counts classified as DNA–RNA coupled, RNA-only state change, isoform-remodeling context, DNA-only alteration, or no clear stemness/lineage change. **h,** Stemness and lineage-plasticity outcome burden by patient. Stacked bars summarize outcome classes among stemness/lineage target genes, including DNA-only alteration, expressed somatic ALT, isoform/usage remodeling, methylation–RNA coupling, ORF/protein-fate context, recurrence-expressed somatic ALT, SV-linked RNA architecture, and SV/ORF-linked RNA architecture.

We next focused on glioma drivers and tumor suppressors. Patient-level outcome summaries showed that driver/suppressor genes were frequently altered across multiple consequence classes, with large contributions from methylation–RNA coupling, ORF/protein-fate context, DNA-only alterations, isoform-specific mutant expression, expressed somatic ALT, recurrence-expressed somatic ALT, and SV-linked RNA architecture (Fig. 10b). Mechanistic flow analysis showed that CNV-dominant, promoter DMR-dominant, multi-layer DNA remodeling, expressed somatic ALT, and SV-linked mechanisms could each connect to distinct RNA consequences and, in selected cases, to predicted ORF/protein-fate outcomes (Fig. 10c). Exemplar summaries highlighted recurrently relevant driver/suppressor genes, including ATRX, PTEN, BCOR, MET, and EGFR, illustrating how different genes entered the integration framework through different DNA mechanisms and RNA/ORF consequence routes (Supplementary Fig. 18b). Thus, canonical glioma genes were not only altered at the DNA level; subsets showed evidence of propagation into RNA remodeling or predicted coding consequences.

Immune, antigen-presentation, and interferon-related genes showed a related but distinct pattern. Most immune-related gene-patient pairs with detectable activity fell into a DNA–RNA coupled category, while smaller subsets showed RNA-only microenvironment or state-associated signal, DNA-only change, or no clear immune change (Fig. 10d). Patient-level outcome burdens showed that immune/antigen/interferon genes were dominated by methylation–RNA coupling and ORF/protein-fate context, with additional contributions from isoform/usage remodeling, SV-linked RNA architecture, SV/ORF-linked RNA architecture, expressed somatic ALT, and DNA-only alterations (Fig. 10e). Mechanistic flow analysis again emphasized promoter DMR-dominant and multi-layer DNA remodeling routes, with smaller SV-linked RNA and ORF-linked paths (Fig. 10f). Exemplar immune summaries included genes such as HLA-A, ISG15, JAK2, CD47, NLRC5, and ITGAM, showing that immune-related remodeling was heterogeneous across patients and mechanisms rather than captured by a single immune-evasion axis (Supplementary Fig. 18c).

Finally, we examined stemness and lineage-plasticity genes. Similar to the immune set, stemness/lineage gene-patient pairs were most often classified as DNA–RNA coupled, followed by RNA-only state change, no clear lineage change, isoform-remodeling context, and DNA-only change (Fig. 10g). Patient-level burdens showed recurrent involvement of methylation–RNA coupling, ORF/protein-fate context, isoform/usage remodeling, expressed somatic ALT, recurrence-expressed somatic ALT, SV-linked RNA architecture, SV/ORF-linked RNA architecture, and DNA-only alterations (Fig. 10h). The corresponding mechanistic flow showed promoter DMR-dominant, CNV-dominant, multi-layer DNA remodeling, expressed somatic ALT, and SV-linked RNA architecture routes converging on methylation–RNA coupling, isoform/usage remodeling, recurrence-expressed somatic ALT, and predicted ORF/protein-fate categories (Supplementary Fig. 18d). Exemplar summaries included lineage and plasticity-associated genes such as TEAD1, TEAD2, MYC, CD44, JAG1, AQP4, PROM1, and STK3, further illustrating patient-specific routes into lineage-state remodeling (Supplementary Fig. 18e).

Together, these targeted analyses show that the DNA–RNA–ORF integration framework converges on biologically interpretable systems rather than producing a random catalog of events. Splicing/RBP regulators provide a direct link to the isoform-remodeling phenotype, driver/suppressor genes anchor the analysis in canonical astrocytoma biology, immune/antigen/interferon genes highlight microenvironment and immune-state remodeling, and stemness/lineage genes connect genomic remodeling to cellular plasticity. The dominant conclusion remains patient-specific: recurrent astrocytomas use different combinations of CNV, methylation, expressed variants, SV-linked RNA architecture, isoform remodeling, and predicted ORF/protein-fate changes to remodel biologically relevant programs.

## Discussion

In this study, we used paired long-read DNA and RNA sequencing to show that recurrent astrocytoma remodeling is not only a genetic or transcriptional process, but a multi-layer DNA-to-RNA-to-isoform-to-ORF remodeling trajectory. The central finding is that genetic and epigenetic alterations acquired or retained during recurrence can be traced to distinct RNA consequences, including gene-expression shifts, isoform redistribution, allele- or haplotype-specific transcript usage, expressed mutant RNA, SV-linked transcript architecture, and predicted ORF/protein-fate changes. Importantly, these effects were not uniform across patients or DNA layers. CNV provided the broadest dosage-associated RNA signal, SVs provided the clearest architectural DNA-to-RNA links, expressed SNVs/indels provided high-specificity mutant RNA and isoform-specific mutant-expression events, and DMRs showed context-dependent relationships with expression, isoform usage, and predicted coding output. Thus, the main contribution of this study is not simply that long-read sequencing detects more events, but that matched long-read DNA and full-length RNA can connect recurrence-associated genome and epigenome remodeling to patient-specific transcript and predicted coding consequences.

Glioma recurrence has been extensively studied using short-read genomic, transcriptomic, and epigenomic profiling, but most prior studies have described recurrence through molecular layers that remain partly disconnected. Paired glioblastoma studies showed that primary and recurrent tumors are clonally related yet often diverge through patient-specific genetic evolution, therapy-associated selection, expression-state switching, and altered copy-number structure^4,18^. Large-scale diffuse glioma studies extended this framework by showing that many founding driver alterations are retained at recurrence, whereas recurrent tumors acquire heterogeneous changes in aneuploidy, subclonal architecture, hypermutation, cell-cycle alterations, transcriptional state, and microenvironmental composition^5,6,19^. More recent longitudinal single-cell and multi-omic studies further emphasized that glioma progression cannot be understood from DNA alone, because genetic evolution is accompanied by changes in malignant-cell state, DNA methylation, immune and stromal composition, proteomic state, and treatment-associated ecosystem remodeling^7,20,21^. Several studies have also interrogated IDH-mutant astrocytoma recurrence using multi-omics approaches. Vallentgoed et al.^8^ identified cell cycling, altered differentiation, extracellular-matrix remodeling, and DNA methylation loss as convergent features of malignant progression. Rodriguez Almaraz et al.^9^ showed that recurrent high-grade tumors can acquire diverse RAS–MAPK pathway alterations associated with inferior survival, emphasizing that progression can converge at the pathway level despite heterogeneity in the individual genomic event. Rautajoki et al.^10^ reported increased rearrangements and deletions during progression to grade 4, together with recurrent CDKN2A/RB1 inactivation, activating PDGFRA/MET alterations, DNA-repair changes, and NRG3 loss. Tang et al.^11^ showed that recurrent IDH-mutant astrocytomas increasingly adopt proliferative/progenitor and immune/mesenchymal-enriched proteomic states, with the latter characterized by gemistocytic differentiation and immune infiltration. Our findings fit an emerging disease-specific model in which IDH-mutant astrocytomas retain their founding molecular identity while acquiring heterogeneous progression-associated changes. Further, despite smaller cohort, our findings complement and support these previous studies. However, the previous approaches using in the glioma or astrocytoma cohorts remain constrained by the read structure of short-read technologies. Short-read DNA sequencing is powerful for SNVs, indels, copy-number alterations, and mutational signatures, but has limited ability to resolve complex rearrangements, long-range haplotypes, repeat-associated structures, phased methylation states, and allele-specific structural context. Short-read RNA sequencing captures gene expression and local splice junctions, but generally infers transcript structures indirectly and cannot routinely assign full-length isoforms, fusion architecture, mutant alleles, haplotype-specific transcript usage, and predicted ORF consequences to the same RNA molecule. Thus, prior short-read multi-omic studies have been powerful for defining recurrence-associated layers, but less able to determine how DNA remodeling is expressed through full-length RNA and coding-output remodeling within the same patient.

The current approach differs also from classical long-read sequencing fundamentally. Many cancer cohorts long-read studies have used long-read DNA or RNA as powerful single-layer discovery tools^14,22^. Long-read DNA studies have resolved complex rearrangements, haplotypes, methylation landscapes, viral integrations, ecDNA, and repeat-associated structural events that are difficult to interpret with short reads^14,15^. Long-read transcriptome studies have catalogued extensive cancer-associated isoform diversity, including unannotated transcripts and isoform-level programs with clinical or biological relevance^12,13^. These studies establish the value of long-read sequencing, but they usually ask either a genome-architecture question or an isoform-discovery question. Here, the long-read sequencing was used to establish a direct DNA-RNA-ORF/proteome potential causal chain. To our knowledge, this paired longitudinal long-read DNA and full-length RNA integration has not yet been applied to glioma or to other cancer cohorts.

An important advantage of this study is the introduction of new analytical modalities to resolve complex isoform remodeling and to connect the DNA and RNA layers. To date, standard long-read RNA analysis workflows often stops at transcript discovery, structural-category annotation, presence/absence analysis, or differential isoform abundance^12,13,23^. Those approaches are useful, but they cannot resolve whether there is remodeling in isoform architecture without large gene-expression change. We therefore introduce complementary approaches to quantify isoform remodeling that can detect significant changes even within apparently stable genes. The class A–F framework separated dominant isoform switching, same-dominant secondary remodeling, primary-dominant isoform loss, recurrent-dominant isoform gain, no-dominant remodeling, and stable genes. Jensen–Shannon divergence and maximum absolute delta isoform fraction quantified the overall distributional shift and the largest isoform-level usage change, while Shannon entropy measured whether recurrence made isoform usage more diffuse or more focused. This entropy component is important because two genes can show similar divergence but opposite changes in transcript complexity. In parallel, Dirichlet-multinomial modeling with DRIMSeq provided statistical evidence of differential transcript usage^16^. These analyses are not redundant: JSD/dIF/entropy prioritize effect size and architecture, whereas DTU prioritizes statistical redistribution of isoform proportions. Together with ORF prediction and transcript-to-genome coordinate lifting^17^, this framework converts long-read RNA from an isoform catalog into a consequence-aware model of isoform and predicted coding-output remodeling. Another important advantage of this work is the demonstration of the power of long read multiomics in resolving how genomic events can be reflected at the gene, isoform, or even protein level. A common problem in multi-omics is that overlap is mistaken for mechanism. If all DNA events are used as the denominator, many silent or non-quantifiable events dilute DNA-to-RNA reflection. If only RNA-altered genes are considered, broad DNA context can be overinterpreted as causal support. We therefore separated global DNA events from RNA-detected genes, patient-matched RNA-detected patient-gene pairs, and RNA-quantifiable patient-gene pairs. We also distinguished high-specificity DNA–RNA/ORF support from directional or layer-concordant support and broad DNA context. Such a framework allowed us to understand the complex and multilayered remodeling of glioma recurrence to an extent that cannot be equally revealed using standard short read sequencing methods.

Biologically, the study supports a model in which recurrent gliomas preserve core driver identity while remodeling how that identity is expressed through RNA architecture and predicted coding output. This patient-specific structure is consistent with previous longitudinal glioma studies, but the long-read design reveals an additional layer: recurrence can be expressed through isoform redistribution, altered transcript-boundary usage, allele- or haplotype-specific isoform choice, mutant isoform expression, and predicted ORF/protein-fate changes. Targeted gene-set analyses further showed that integrated DNA–RNA–ORF chains converge on biologically interpretable systems, including splicing and RNA-binding proteins, glioma drivers and tumor suppressors, immune and interferon-related genes, and stemness or lineage-plasticity programs. These analyses do not prove pathway causality, and they should not be read as functional validation. Their value is that they prioritize coherent, patient-specific remodeling chains that can be tested experimentally. Expanding this framework to larger cohorts will therefore be essential for identifying novel targets and gene sets that can be exploited as glioma vulnerabilities.

Several limitations should be acknowledged directly. The cohort is small, DNA-dependent recurrence analyses were limited to five evaluable patient pairs and predicted ORF/protein-fate consequences are computational rather than direct proteomic measurements. 2 patients had lower purity of either primary or recurrent tumor samples, indicating lower tumor coverage. While these 2 patients did not appear to be major outliers in different analyses, the results should be interpreted carefully. Bulk tumor profiling cannot fully separate malignant-cell-intrinsic remodeling from changes in tumor purity, lineage composition, or microenvironmental admixture. Despite these limitations, this study provides a proof-of-principle framework for longitudinal long-read multi-omics in recurrent glioma. Future work should extend this approach to larger molecularly stratified cohorts, integrate proteomics or Ribo-seq where tissue permits, and functionally validate selected SV-, DMR-, ASTU-, and mutant-isoform-linked candidates. The broader implication is that precision oncology in recurrent glioma may benefit from moving beyond lists of DNA alterations or gene-expression changes toward patient-specific DNA-to-RNA-to-isoform-to-ORF remodeling chains.

## Methods

### Patient cohort

Among participants enrolled in a study approved by the Ethics Committee of Tohoku University Hospital (2025-1-1029), six patients with IDH-mutant astrocytoma were selected based on the availability of paired tumor tissues resected at initial diagnosis and recurrence. Tumor specimens were flash-frozen in liquid nitrogen immediately after surgical resection and stored at −80°C until nucleic acid extraction. Peripheral blood samples were collected before the initial surgery in EDTA-2Na blood collection tubes, and whole blood was stored at −80°C until DNA extraction. Written informed consent was obtained from all participants. Cohort information are presented in Supplementary table 1. The study was approved by Tohoku University Hospital’s ethical review board (# 2023-1-446).

### DNA extraction

High-molecular-weight genomic DNA was extracted using the Nanobind PanDNA kit (PacBio, Cat. No. 103-260-000) according to the manufacturer’s protocol. For tumor specimens, approximately 10 mg of frozen tissue was dissected using a scalpel and homogenized by 10 strokes using a Dounce homogenizer. The homogenate was then transferred to a new tube, and the remaining extraction steps were performed according to the kit protocol. For matched blood samples, high-molecular-weight genomic DNA was extracted from 400 µL of whole blood using the whole-blood RBC lysis protocol of the Nanobind PanDNA kit. Briefly, whole blood was subjected to red blood cell lysis, followed by white blood cell pelleting and genomic DNA extraction using the Nanobind-based workflow. DNA was assessed for quantity and quality using NanoDrop spectrophotometry, Qubit fluorometry, and the Agilent Femto Pulse system with the Genomic DNA 165 kb kit (Agilent, Cat. No. FP-1002-0275). All genome quality numbers (GQN) at 10k values were greater than 7.0, with a median of 9.9.

### RNA extraction

Total RNA was extracted using the RNeasy Mini kit (QIAGEN, Cat. No. 74104) according to the manufacturer’s protocol. Briefly, approximately 10 mg of each frozen tumor sample was homogenized in QIAzol Lysis Reagent (QIAGEN, Cat. No. 79306), followed by phase separation with chloroform. The aqueous phase was transferred to a new tube, and the remaining purification steps were performed according to the RNeasy Mini kit protocol. Extracted RNA was assessed for quantity and quality using NanoDrop spectrophotometry, Qubit fluorometry, and an Agilent 2100 Bioanalyzer with the RNA 6000 Nano kit (Agilent, Cat. No. 5067-1511). All samples used had RNA integrity number (RIN) values were greater than 7.0 with a median of 7.3.

### Preparation of whole-genome sequencing libraries for long-read sequencing

Long-read whole-genome sequencing libraries were prepared from high-molecular-weight genomic DNA using the SMRTbell Prep Kit 3.0 (PacBio, Cat. No. 102-182-700) according to the manufacturer’s protocol. Briefly, short DNA fragments were depleted from genomic DNA samples using the Short Read Eliminator kit (PacBio, Cat. No. 102-208-300), followed by DNA shearing using the Hamilton Microlab Prep system at Haplo Pharma Inc., Sendai, Japan. Up to 3,000 ng of genomic DNA was processed using the high-mass workflow, depending on sample yield, and samples with limited DNA yield were processed using the low-mass workflow. If shearing was insufficient, additional shearing was performed until the average fragment size reached approximately 20–25 kb. Sheared DNA was then converted into SMRTbell libraries using the SMRTbell Prep Kit 3.0. The final libraries were evaluated using the Agilent Femto Pulse system at Haplo Pharma Inc., Sendai, Japan, and libraries with an average size of 15–20 kb were used for sequencing.

Libraries were sequenced on the PacBio Revio sequencing platform at PacBio, Menlo Park, CA, USA. Sequencing was performed with target coverages of 60× for tumor tissue samples, corresponding to 1.5 SMRT Cell equivalents per sample, and 30× for blood samples, corresponding to 0.75 SMRT Cell equivalents per sample.

### Preparation of Kinnex libraries for long-read RNA sequencing

Full-length RNA long-read sequencing libraries were prepared using the Iso-Seq Express 2.0 Kit (PacBio, Cat. No. 103-071-500) and the Kinnex PCR 8-fold kit (PacBio, Cat. No. 103-071-600) according to the manufacturer’s protocol. Briefly, 300 ng of total RNA was used for cDNA synthesis, followed by cDNA amplification and Kinnex PCR. The Kinnex PCR products were then used to generate Kinnex arrays. For multiplex sequencing, each sample was assigned a unique barcode combination. The final libraries were assessed using the Agilent Femto Pulse system. Libraries were sequenced on the PacBio Revio system with a target output of 10 million HiFi reads per sample.

### Long Read RNA data analysis

Long read RNA sequencing were analyzed using the isoseq-pigeon workflow (https://isoseq.how/). However, for Allele specific transcript usage (ASTU), we mapped the long read RNA reads after the Refine step in the isoseq workflow. All other analysis was performed using the classical workflow and outputs. All downstream analysis was performed in *R studio*.

### Long read DNA data analysis

Long read DNA sequencing was analyzed using the standard HiFi-somatic-WDL workflow (https://github.com/PacificBiosciences/HiFi-somatic-WDL). All downstream analysis using outputs from the HiFi-somatic workflow and was performed in *R studio*.

### Long-read RNA annotation and full-length count matrix construction

Pigeon filtered classification and junction outputs were parsed for each Iso-Seq sample and merged into a sample-level transcript annotation table. Isoforms were harmonized across samples using Pigeon/PacBio isoform identifiers together with junction-chain signatures to reduce sample-specific identifier ambiguity. For each transcript, we retained the associated gene, Pigeon structural category, full-length read count, transcript coordinates, coding annotation when available, predicted NMD status when available, and sample metadata. Gene-level full-length counts were calculated by summing full-length reads across all isoforms assigned to the same gene in each sample. Isoform-level analyses used the harmonized isoform identifier as the primary transcript unit.

Gene-level differential expression was performed using full-length read counts summed per gene and modeled with DESeq2^24^ using a paired design where timepoint compared recurrent versus primary tumors. Genes with fewer than 10 total full-length reads across all samples were excluded before model fitting. Isoform-level differential abundance was similarly modeled with DESeq2 using full-length isoform counts and the same paired design. Isoforms were retained if they were detected in at least four patients in both primary and recurrent groups, had at least 10 total full-length reads, and mapped to genes passing a relaxed gene-level differential-expression screen. Gene-set activity was estimated from variance-stabilized gene-level counts using ssGSEA across Hallmark, Reactome, oncogenic, and immunologic gene-set collections, followed by paired recurrent-primary testing.

### Isoform gain/loss and gene-level isoform remodeling

Isoform gain and loss were evaluated using support-filtered isoforms. An isoform was considered present in a sample if it had at least five full-length reads and accounted for at least 10% of the gene’s full-length read support in that sample. High-confidence recurrent-gained or primary-lost isoforms required support in at least four patients.

For gene-level isoform remodeling, we calculated isoform fractions within each gene. For gene *g*, isoform *i*, and group *t*where *t* ∈ {primary,recurrent}, the mean isoform fraction was defined as:

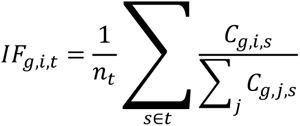

where *C_g_*_,*i*,*s*_is the full-length read count for isoform *i*of gene *g*in sample *s*, and the denominator is the total full-length read count for all isoforms of gene *g*in that sample.

For each gene, the primary-to-recurrent isoform-fraction shift for isoform *i*was:

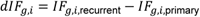

and the largest isoform-level usage change was:

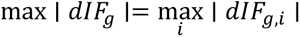

Distributional divergence between primary and recurrent isoform usage was measured using Jensen–Shannon divergence. For the primary and recurrent isoform-fraction vectors *P_g_*and *Q_g_*:

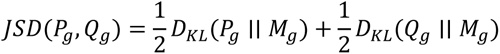

where:

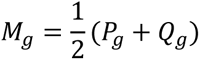

and:

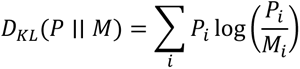

A small pseudocount was added before normalization to avoid undefined logarithms. Genes were eligible for remodeling analysis if they had at least 20 total full-length reads and at least two expressed isoforms with at least five total full-length reads. High-confidence isoform remodeling required *JSD* ≥ 0.10, max ∣ *dIF* ∣≥ 0.20, and patient-level support in at least four patients.

Dominant isoforms were defined using both group-level and patient-level support. A dominant isoform required mean isoform fraction ≥0.35, margin over the second-ranked isoform ≥0.10, and support in at least four patients. Genes were then assigned to six remodeling classes: Class A, dominant isoform switching; Class B, same dominant isoform with secondary isoform remodeling; Class C, primary dominant isoform lost; Class D, recurrent dominant isoform gained; Class E, no-dominant remodeling with strong distributional change; and Class F, stable or not high-confidence remodeled.

### Shannon entropy analysis of isoform complexity

To determine whether recurrence made isoform usage more diffuse or more focused, Shannon entropy was calculated for each gene’s primary and recurrent isoform-fraction distributions:

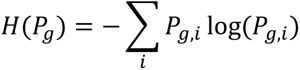

and:

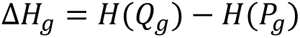

where *P_g_*and *Q_g_*are the primary and recurrent isoform-fraction vectors. Positive Δ*H*indicates more diffuse isoform usage in recurrence, whereas negative Δ*H*indicates more focused isoform usage. A normalized entropy was also calculated as:

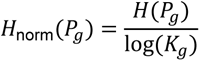

where *K_g_*is the number of expressed isoforms. For visualization, high-divergence entropy classes used *JSD* ≥ 0.20and ∣ Δ*H* ∣≥ 0.20.

### Differential transcript usage analysis

Differential transcript usage was tested using DRIMSeq^16^. Full-length isoform counts were modeled within genes using a Dirichlet-multinomial framework with patient pairing. The design matrix included patient and timepoint. Only genes with at least two retained isoforms were tested. Isoforms were required to reach at least 1% within-gene proportion in at least four samples before DRIMSeq filtering. Additional DRIMSeq filters required gene expression in at least four samples, minimum gene count of 20, isoform expression in at least four samples, and minimum isoform count of 10. Gene-level and isoform-level DTU statistics were extracted, and stageR was used for two-stage adjustment of gene-level screening and isoform-level confirmation. DTU results were interpreted as statistical evidence of transcript-usage redistribution, complementary to the JSD/dIF/entropy framework, which prioritizes effect size and transcript-architecture remodeling.

### Predicted ORF and protein-output annotation

Genemark-hmm^17^ was used to predict ORFs from long read transcripts. First, per sample long read transcript sequences were exported, and Genemark-hmm was used to predict the ORFs, limiting the predicted ORFs to maximum of one per transcript to reduce complexity, and using standard settings. Predicted ORFs were parsed from GeneMarkS-T/GeneMark transcript-coordinate GFF outputs. ORF coordinates were lifted from transcript space to genome coordinates using PacBio/Pigeon isoform exon models. Lifted ORFs were compared with canonical GENCODE v49 coding sequence annotations, prioritizing MANE Select transcripts where available, then Ensembl canonical transcripts, and otherwise the longest coding transcript.

ORFs were classified relative to the canonical coding sequence as annotated/canonical, alternative canonical, upstream ORF, downstream ORF, upstream-overlapping ORF, downstream-overlapping ORF, internal ORF, N-terminal truncation, C-terminal truncation, span-both, CDS-overlap other, no canonical CDS, no ORF, or unknown/unlifted ORF. NMD potential was estimated using an exon-junction-complex heuristic, where predicted stop codons located at least 55 nucleotides upstream of the last exon-exon junction were flagged as NMD-compatible. Frameshift potential was annotated for CDS-overlapping ORF classes by comparing the lifted ORF start frame with the canonical CDS start frame. ORF/protein-output consequences were treated as computational predictions and not as direct proteomic measurements.

### Denominator aware DNA–RNA integration framework

DNA event calls were imported from the established HiFi-somatic analysis outputs, including somatic SNV/indel, SV, CNV/LOH, DMR/methylation, mutational-signature, and haplotype/allelic-context tables. The integration analysis did not re-call DNA events. Instead, DNA events were standardized to common patient, timepoint, gene, event-class, and recurrence-status fields and then linked to long-read RNA features.

To avoid denominator ambiguity, DNA events were summarized using progressively RNA-aware denominators: all DNA events; DNA events occurring in RNA-detected genes; DNA events occurring in patient-matched RNA-detected patient-gene pairs; and DNA events occurring in RNA-quantifiable patient-gene pairs. A patient-gene pair required DNA and RNA information from the same patient and gene context. RNA-quantifiable patient-gene pairs required sufficient RNA support to evaluate gene expression, isoform usage, DTU, ASE/ASTU, or ORF/protein-fate features.

RNA alterations were grouped into core consequence layers, including gene-expression remodeling, isoform-usage remodeling, DTU, SV-linked RNA architecture, expressed somatic ALT RNA, isoform-specific mutant expression, ASE/ASTU, and predicted ORF/protein-fate consequences. RNA alterations were then stratified by DNA support specificity: high-specificity DNA–RNA/ORF support, directional or layer-concordant support, broad DNA context, no available DNA, or no detected DNA support. High-specificity support included direct relationships such as expressed somatic ALT RNA, isoform-specific mutant expression, SV-linked RNA architecture, or DNA–RNA–ORF chains. Directional/layer-concordant support included relationships such as CNV gain with increased RNA expression, CNV loss with decreased RNA expression, or compatible promoter methylation-expression patterns. Broad DNA context indicated patient-matched DNA alteration in the same gene without a specific directional or event-level RNA mechanism.

### Layer-specific DNA–RNA analyses

For CNV-RNA coupling, recurrent-primary CNV delta was calculated for each patient-gene pair using gene-level copy-number summaries. Absolute copy-number-like changes of at least 0.5 were classified as CNV gain or loss. CNV-RNA relationships were classified as concordant when CNV gain associated with RNA upregulation or CNV loss associated with RNA downregulation, discordant when the opposite pattern was observed, or isoform/DTU/ORF-coupled when copy-number context overlapped RNA architecture remodeling without simple expression concordance.

For somatic variant expression, somatic SNVs from the HiFi-somatic pipeline were scanned in genome-aligned Iso-Seq RNA BAM files. Each RNA read overlapping a somatic SNV was classified as supporting the reference allele, somatic ALT allele, other allele, deletion/gap, or no-call. High-confidence RNA ALT expression required at least three ALT-supporting reads, at least five total informative reads, and ALT fraction ≥0.10. Sensitive ALT expression required at least two ALT-supporting reads and ALT fraction ≥0.05. RNA reads were then linked back to Pigeon isoforms to evaluate isoform-specific mutant allele expression and predicted ORF/protein-output consequences.

For mutational-signature integration, temporal somatic mutation catalogs were exported for SigProfiler-based SBS signature assignment^25,26^. Signature-context mutations were then overlaid with RNA ALT expression, recurrence-gained mutation status, isoform-specific mutant expression, nonsynonymous/splice annotation, and predicted ORF/protein-output consequences.

For ASE, high-confidence germline heterozygous SNPs were selected from matched normal DNA and intersected with RNA reads. At each RNA-covered heterozygous marker, reference and alternate allele counts were obtained. Gene-level ASE was calculated by aggregating informative SNPs within each gene and sample. A gene-level ASE call required at least three informative SNPs, at least 30 total ref+alt RNA counts, gene-level FDR ≤0.05, and weighted major-allele fraction ≥0.65. Strong ASE used weighted major-allele fraction ≥0.75. High-confidence recurrent ASE candidates required at least five informative SNPs and at least 50 ref+alt RNA counts in each compared timepoint.

For haplotype-resolved ASTU, Iso-Seq reads were assigned to HP1 or HP2 using normal HiPhase germline haplotype markers. HP1 and HP2 are technical haplotype labels and were not interpreted as maternal or paternal alleles. Haplotype-assigned reads were linked to Pigeon isoforms, and isoform fractions were calculated separately for each haplotype-marker group. Within-sample ASTU was summarized using HP1-versus-HP2 JSD and maximum absolute isoform-fraction difference. Recurrence-specific ASTU was tested per patient-gene using a Poisson model comparing:

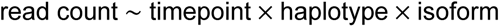

against a reduced model without the three-way interaction:

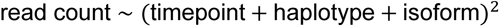

The three-way interaction tested whether haplotype-specific isoform usage changed between primary and recurrent tumors. Recurrence-ASTU candidates required sufficient reads in both timepoints and both haplotypes, at least two isoforms, FDR ≤0.10 or a recurrence-associated effect-size change, and maximum interaction ∣ *dIF* ∣≥ 0.25.

For DMR-RNA coupling, DMRs were grouped by regulatory context, including promoter, TSS-proximal, gene-body/transcript, intronic, exonic, UTR, and proximal classes. Paired recurrent-primary methylation direction was summarized per patient-gene-DMR class. Promoter/TSS DMRs were compared with patient-level gene expression direction and classified as concordant, discordant, stable, or insufficiently expressed. DMRs in other regulatory contexts were overlaid with isoform switching, DTU, TSS/TTS remodeling, junction remodeling, and predicted ORF/protein-fate consequences. Because methylation-RNA relationships are context-dependent, they were interpreted as associations unless supported by additional RNA architecture or ORF evidence.

For SV-RNA architecture integration, SV breakpoints were annotated relative to genes, promoters, TSS/TTS windows, and Pigeon isoform boundaries. Breakpoints were linked to nearby isoform TSS/TTS or junction features, recurrently gained isoforms, SV-linked RNA junction evidence, fusion-support evidence where available, isoform remodeling, and predicted ORF/protein-fate consequences. SV-linked RNA architecture support was interpreted as evidence of structural proximity or gene-level RNA architectural remodeling, not as base-pair-resolution fusion validation unless direct fusion support was present.

### Targeted biological gene-set integration

Targeted biological gene sets were analyzed using the same standardized DNA–RNA integration framework. For each gene set, patient-gene pairs were annotated for DNA events, RNA consequences, expressed somatic ALT RNA, isoform-specific mutant expression, SV-linked RNA architecture, methylation-RNA coupling, isoform/DTU remodeling, and predicted ORF/protein-fate consequences. Gene-patient pairs were classified as DNA-only, RNA-only, DNA–RNA coupled, or multi-layer DNA–RNA coupled. Candidate prioritization used a weighted score incorporating target priority, recurrence-gained DNA events, number of DNA layers, splice-site/splice-region variants, SVs, CNVs, promoter DMRs, mutational-signature context, gene-expression change, isoform/DTU remodeling, somatic ALT RNA expression, isoform-specific mutant expression, SV-linked RNA architecture, methylation-RNA coupling, predicted ORF/protein-fate evidence, and recurrence across patients. These scores were used only to rank and summarize candidate patterns, not as validation of causality.

The targeted gene sets used for the biological integration figure were:

### Splicing regulators and RNA-binding proteins

SF3B1, SRSF2, U2AF1, U2AF2, ZRSR2, PRPF8, SNRNP200, SF3A1, SF3A2, SF3A3, SF3B2, SF3B3, SF3B4, SNRPA1, SNRPB, SNRPC, SNRPD1, SNRPD2, SNRPD3, SNRPE, SNRPF, SNRPG, SRSF1, SRSF3, SRSF6, SRSF7, SRSF9, TRA2A, TRA2B, HNRNPA1, HNRNPA2B1, HNRNPC, HNRNPK, HNRNPM, PTBP1, PTBP2, QKI, RBFOX1, RBFOX2, ESRP1, ESRP2, MBNL1, MBNL2, CELF1, NOVA1, NOVA2, RBM10, RBM15, RBM39, DDX5, DDX17, DHX9, DHX15, DHX16, DHX38, EFTUD2, BUD13, SNW1.

### Glioma drivers and tumor suppressors

IDH1, IDH2, TP53, ATRX, EGFR, PTEN, NF1, RB1, CDKN2A, CDKN2B, PDGFRA, MET, MDM2, MDM4, CDK4, CDK6, PIK3CA, PIK3R1, TERT, BRAF, CIC, FUBP1, NOTCH1, NOTCH2, SETD2, ARID1A, SMARCA4, BCOR, KMT2D, KDM6A, MYC, MYCN.

### Immune, antigen-presentation, and interferon-related genes

HLA-A, HLA-B, HLA-C, HLA-DRA, HLA-DRB1, HLA-DPA1, HLA-DPB1, B2M, TAP1, TAP2, TAPBP, NLRC5, CIITA, IFNGR1, IFNGR2, JAK1, JAK2, STAT1, STAT2, IRF1, IRF7, IRF9, ISG15, IFIT1, IFIT2, IFIT3, MX1, OAS1, CXCL9, CXCL10, CXCL11, CD274, PDCD1LG2, CTLA4, LAG3, HAVCR2, TIGIT, LGALS9, CD47, SIRPA, VTCN1, IDO1, IDO2, CSF1, CSF1R, AIF1, ITGAM, CD68, CD163, MRC1, LST1, TYROBP, TREM2, FCGR1A, FCGR2A, CX3CR1, CD3D, CD3E, CD8A, CD8B, GZMB, PRF1, NKG7, GNLY, CXCR3, CCR5.

### Stemness and lineage-plasticity genes

SOX2, POU3F2, SALL2, OLIG2, NES, PROM1, CD44, BMI1, MYC, MYCN, ASCL1, OLIG1, DLL3, SOX10, PDGFRA, BCAN, DCX, NKX2-2, CSPG4, GFAP, AQP4, ALDH1L1, VIM, LCN2, SLC1A3, FABP7, CLU, NOTCH1, NOTCH2, NOTCH3, DLL1, JAG1, JAG2, HES1, HES5, HEY1, YAP1, WWTR1, TEAD1, TEAD2, TEAD3, TEAD4, LATS1, LATS2, MST1, STK4, STK3.

## Acknowledgments

This work was supported by Biomedical Research Core of Tohoku University Graduate School of Medicine. A part of this study was supported by the Support system for young researchers to use research equipment, instruments, and devices in Tohoku University. The authors report no conflict of interest regarding this work. A large language model (LLM; ChatGPT) was used for language editing and code development.

## Funding

This work was funded by PacBio SMRT Cancer grant and the Japan Society for Promotion of Science (JSPS) grant (#23H02741) for **S.R.**

## Author contributions

**S.R.:** Conception, Funding, Formal analysis & Bioinformatics, writing. **S.R., D.A., M.K.:** Study design. **D.A., S.Y., M.K.:** Sample/cohort collection. **D.A.:** Sequencing libraries preparation. **D.A., S.Y., M.K., H.E., K.N.:** review and editing.

## Data availability

Raw long read RNA and DNA sequencing data were deposited in sequencing read archive (SRA) under project number:

## Notes

### Competing Interest Statement

The authors have declared no competing interest.

## References

1 Price, M. et al. CBTRUS Statistical Report: Primary Brain and Other Central Nervous System Tumors Diagnosed in the United States in 2017-2021. Neuro Oncol 26, vi1-vi85, doi:10.1093/neuonc/noae145 (2024).

2 Stupp, R. et al. Radiotherapy plus concomitant and adjuvant temozolomide for glioblastoma. N Engl J Med 352, 987–996, doi:10.1056/NEJMoa043330 (2005).

3 Louis, D. N. et al. The 2021 WHO Classification of Tumors of the Central Nervous System: a summary. Neuro Oncol 23, 1231–1251, doi:10.1093/neuonc/noab106 (2021).

4 Kim, J. et al. Spatiotemporal Evolution of the Primary Glioblastoma Genome. Cancer Cell 28, 318–328, doi:10.1016/j.ccell.2015.07.013 (2015).

5 Ceccarelli, M. et al. Molecular Profiling Reveals Biologically Discrete Subsets and Pathways of Progression in Diffuse Glioma. Cell 164, 550–563, doi:10.1016/j.cell.2015.12.028 (2016).

6 Barthel, F. P. et al. Longitudinal molecular trajectories of diffuse glioma in adults. Nature 576, 112–120, doi:10.1038/s41586-019-1775-1 (2019).

7 Varn, F. S. et al. Glioma progression is shaped by genetic evolution and microenvironment interactions. Cell 185, 2184–2199.e2116, doi:10.1016/j.cell.2022.04.038 (2022).

8 Vallentgoed, W. R. et al. Evolutionary trajectories of IDH-mutant astrocytoma identify molecular grading markers related to cell cycling. Nat Cancer 6, 1693–1713, doi:10.1038/s43018-025-01023-z (2025).

9 Rodriguez Almaraz, E., et al. Longitudinal profiling of IDH-mutant astrocytomas reveals acquired RAS-MAPK pathway mutations associated with inferior survival. Neurooncol Adv 7, vdaf024, doi:10.1093/noajnl/vdaf024 (2025).

10 Rautajoki, K. J. et al. Genomic characterization of IDH-mutant astrocytoma progression to grade 4 in the treatment setting. Acta Neuropathol Commun 11, 176, doi:10.1186/s40478-023-01669-9 (2023).

11 Tang, J. et al. Protein-based classification reveals an immune-hot subtype in IDH mutant astrocytoma with worse prognosis. Cancer Cell 43, 2136–2155.e2114, doi:10.1016/j.ccell.2025.08.006 (2025).

12 Shi, X. et al. Transcript isoform diversity defines molecular subtypes and prognosis in acute myeloid leukemia through long-read sequencing. Cell Rep 44, 116216, doi:10.1016/j.celrep.2025.116216 (2025).

13 Dondi, A. et al. Detection of isoforms and genomic alterations by high-throughput full-length single-cell RNA sequencing in ovarian cancer. Nat Commun 14, 7780, doi:10.1038/s41467-023-43387-9 (2023).

14 Zumalave, S. et al. Concurrent L1 retrotransposition events promote reciprocal translocations in human tumorigenesis. Science 392, eaee4513, doi:10.1126/science.aee4513 (2026).

15 O’Neill, K. et al. Long-read sequencing of an advanced cancer cohort resolves rearrangements, unravels haplotypes, and reveals methylation landscapes. Cell Genom 4, 100674, doi:10.1016/j.xgen.2024.100674 (2024).

16 Nowicka, M. & Robinson, M. D. DRIMSeq: a Dirichlet-multinomial framework for multivariate count outcomes in genomics. F1000Res 5, 1356, doi:10.12688/f1000research.8900.2 (2016).

17 Lukashin, A. V. & Borodovsky, M. GeneMark.hmm: new solutions for gene finding. Nucleic Acids Res 26, 1107–1115, doi:10.1093/nar/26.4.1107 (1998).

18 Wang, J. et al. Clonal evolution of glioblastoma under therapy. Nat Genet 48, 768–776, doi:10.1038/ng.3590 (2016).

19 Brat, D. J. et al. Comprehensive, Integrative Genomic Analysis of Diffuse Lower-Grade Gliomas. N Engl J Med 372, 2481–2498, doi:10.1056/NEJMoa1402121 (2015).

20 Spitzer, A. et al. Deciphering the longitudinal trajectories of glioblastoma ecosystems by integrative single-cell genomics. Nat Genet 57, 1168–1178, doi:10.1038/s41588-025-02168-4 (2025).

21 Dekker, L. J. M. et al. Multiomics profiling of paired primary and recurrent glioblastoma patient tissues. Neurooncol Adv 2, vdaa083, doi:10.1093/noajnl/vdaa083 (2020).

22 Veiga, D. F. T. et al. A comprehensive long-read isoform analysis platform and sequencing resource for breast cancer. Sci Adv 8, eabg6711, doi:10.1126/sciadv.abg6711 (2022).

23 Shi, Q., Li, X., Liu, Y., Chen, Z. & He, X. FLIBase: a comprehensive repository of full-length isoforms across human cancers and tissues. Nucleic Acids Res 52, D124–d133, doi:10.1093/nar/gkad745 (2024).

24 Love, M. I., Huber, W. & Anders, S. Moderated estimation of fold change and dispersion for RNA-seq data with DESeq2. Genome Biol 15, 550, doi:10.1186/s13059-014-0550-8 (2014).

25 Khandekar, A. et al. Visualizing and exploring patterns of large mutational events with SigProfilerMatrixGenerator. BMC Genomics 24, 469, doi:10.1186/s12864-023-09584-y (2023).

26 Bergstrom, E. N. et al. SigProfilerMatrixGenerator: a tool for visualizing and exploring patterns of small mutational events. BMC Genomics 20, 685, doi:10.1186/s12864-019-6041-2 (2019).

