## Supplementary Figures for "Long read multi-omics sequencing reveals DNA-to-RNA evolutionary remodeling trajectories of recurrent astrocytoma"

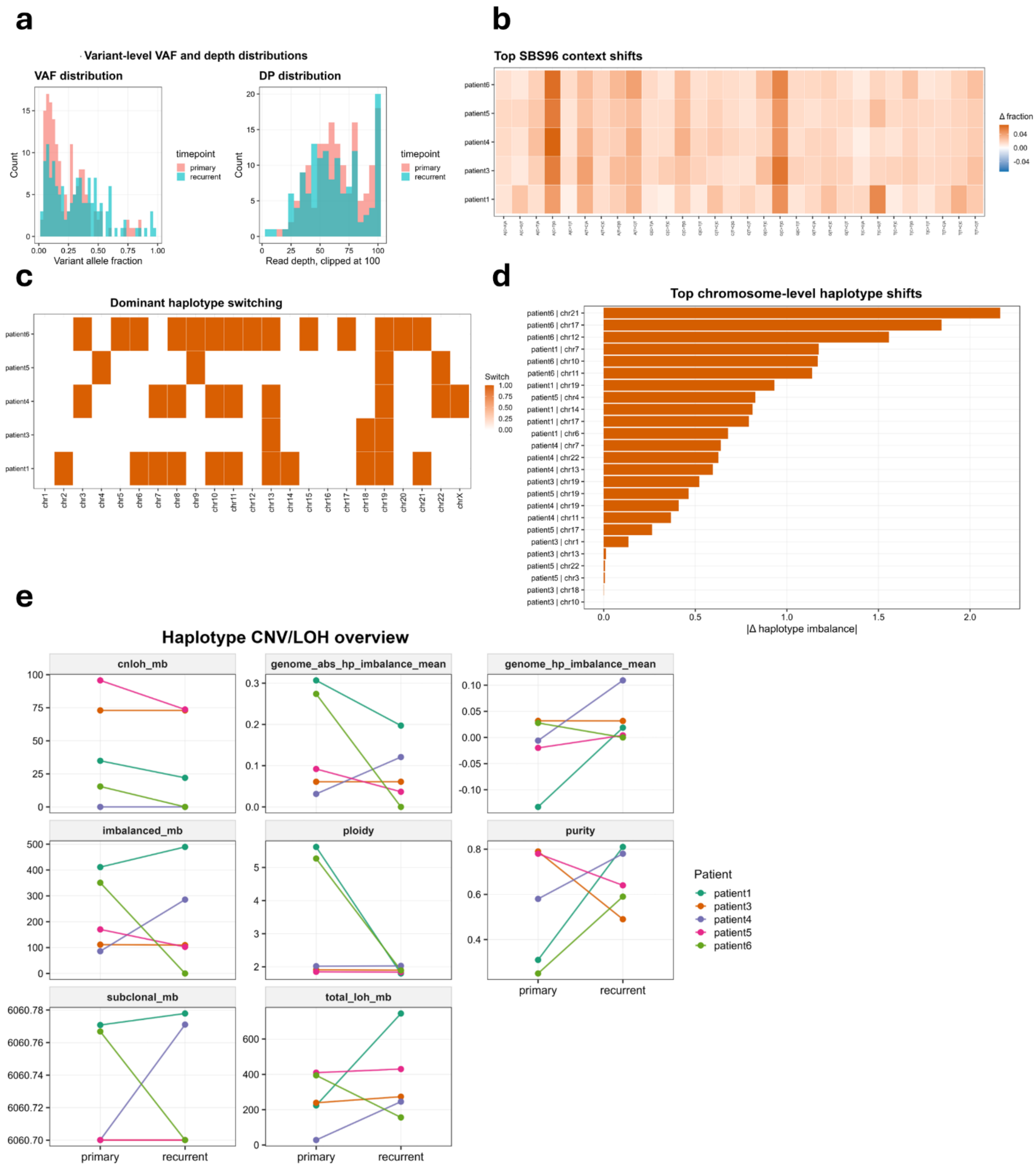


**Supplementary Figure 1. Supporting small-variant, mutational-context, and haplotype analyses.**

**a,** Variant-level SNV/indel VAF and read-depth distributions across primary and recurrent tumors.
**b,** Top SBS96 mutational-context shifts between primary and recurrent tumors, summarized by patient.
**c,** Chromosome-level dominant haplotype switching between primary and recurrent tumors.
**d,** Ranked chromosome-level haplotype imbalance shifts across patient-chromosome pairs.

**e,** Haplotype CNV/LOH overview showing paired changes in copy-neutral LOH, genome-wide haplotype imbalance, mean haplotype imbalance, imbalanced Mb, ploidy, purity, subclonal burden, and total LOH.

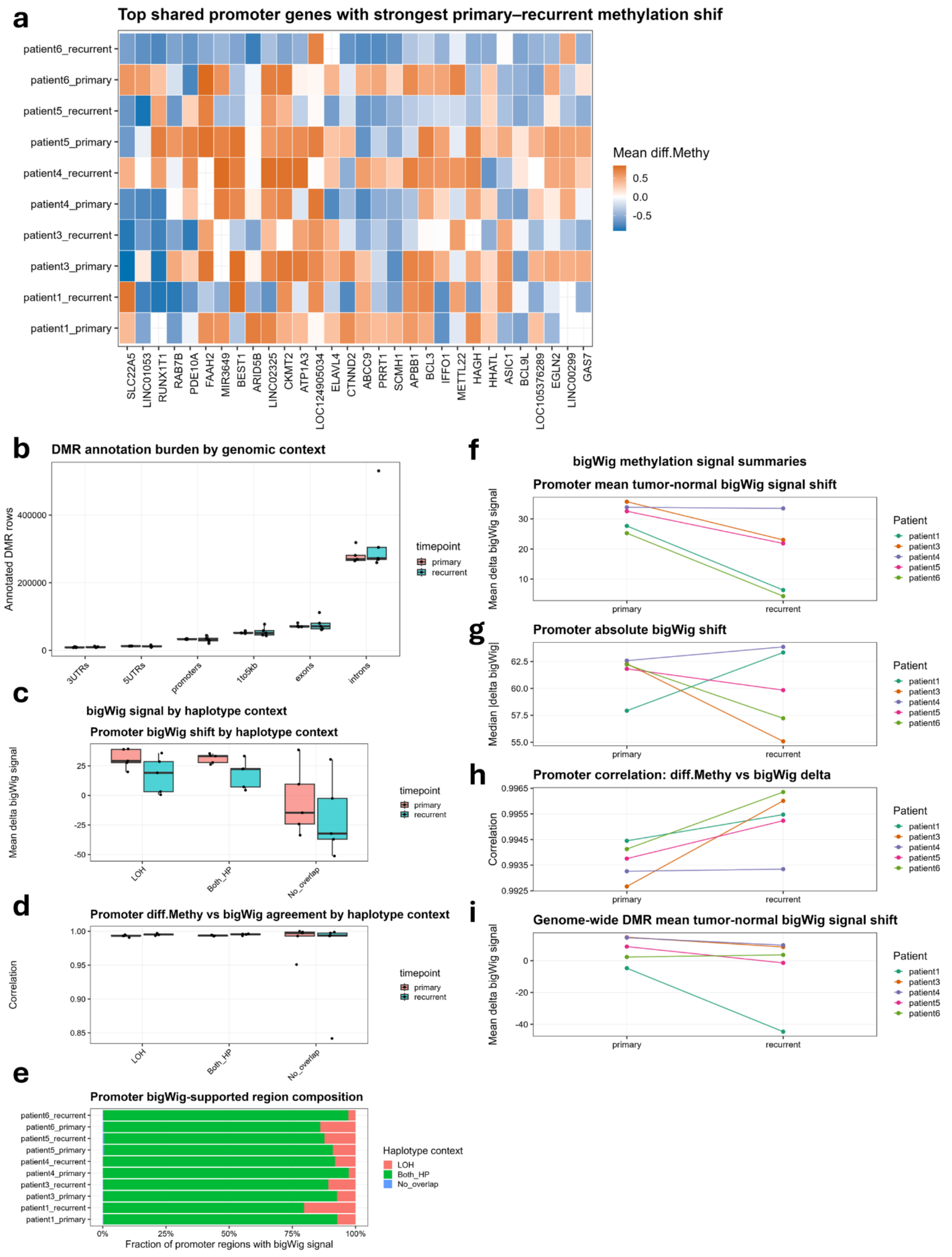


**Supplementary Figure 2. Extended DMR and methylation signal analyses.**

**a,** Heatmap of shared promoter genes with the strongest primary-recurrent methylation shifts.
**b,** DMR annotation burden by genomic context, comparing primary and recurrent tumors across 3′UTR, 5′UTR, promoter, proximal, exon, and intron annotations.
**c,** Promoter bigWig signal shift stratified by haplotype context.
**d,** Agreement between promoter differential methylation estimates and bigWig signal by haplotype context.
**e,** Composition of promoter regions with bigWig methylation signal support across LOH, both-haplotype, and no-overlap contexts.
**f,** Promoter mean tumor-normal bigWig signal shift across paired samples.
**g,** Promoter absolute bigWig methylation signal shift across paired samples.
**h,** Correlation between promoter differential methylation and bigWig methylation delta across paired samples.
**i,** Genome-wide DMR mean tumor-normal bigWig signal shift across primary and recurrent tumors.

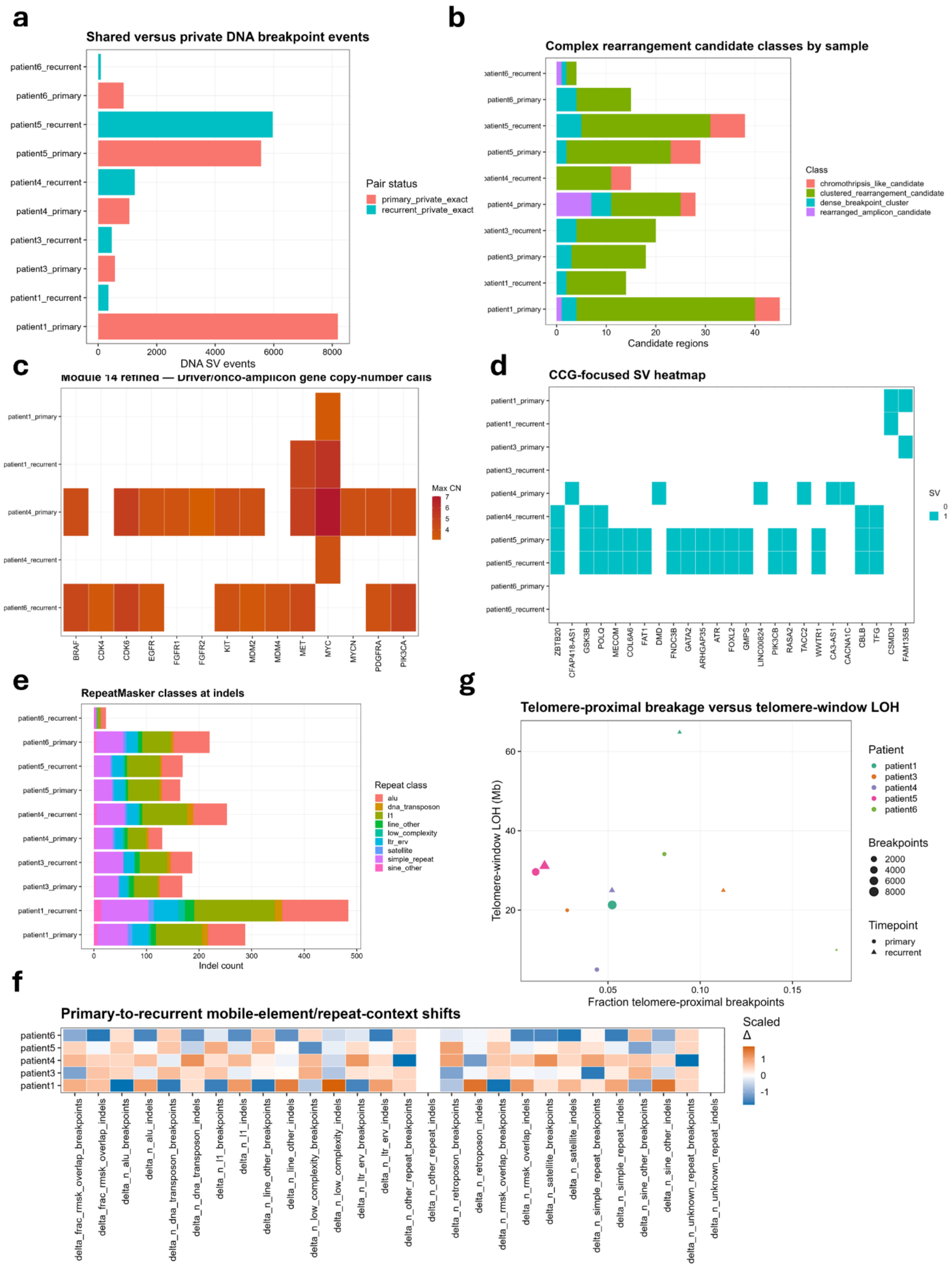


**Supplementary Figure 3. Extended SV, complex rearrangement, amplicon, repeat, and telomere-context analyses.**

**a,** Shared versus private DNA breakpoint events across paired primary and recurrent tumors.
**b,** Complex rearrangement candidate classes by sample, including chromothripsis-like, clustered rearrangement, dense breakpoint cluster, and rearranged amplicon candidate classes. These labels represent structural proxy classes rather than validated mechanistic calls.
**c,** Refined driver/onco-amplicon gene copy-number calls across selected oncogenic loci and samples.
**d,** Cancer-gene-census-focused SV heatmap showing presence or absence of SV overlap with selected cancer-associated genes.
**e,** RepeatMasker classes intersecting somatic indels across samples.
**f,** Primary-to-recurrent mobile-element/repeat-context shifts summarized by patient and repeat-context feature.
**g,** Relationship between telomere-proximal breakpoint fraction and telomere-window LOH, with point size indicating breakpoint burden and shape indicating timepoint.

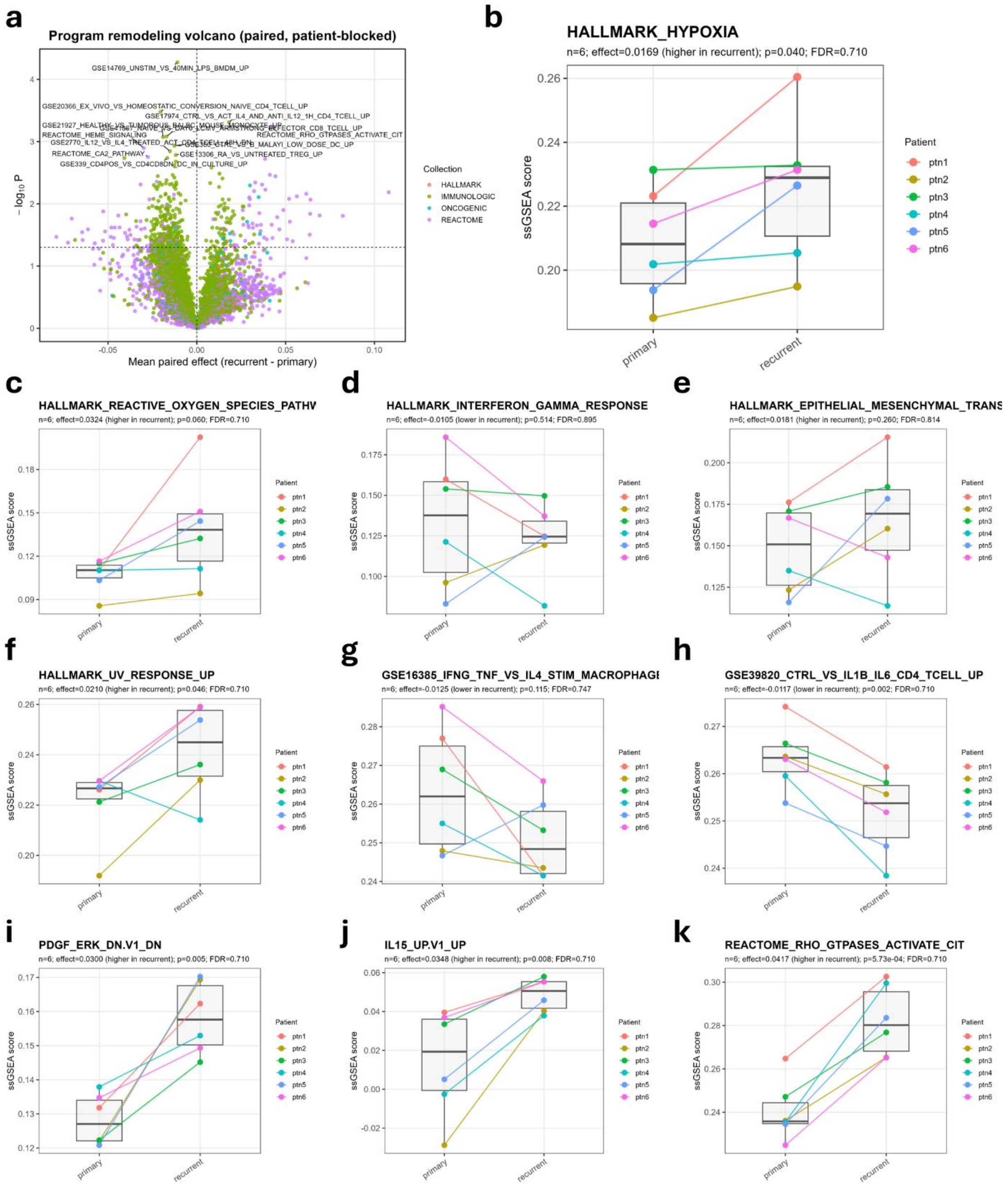


**Supplementary Figure 4. Recurrence-associated gene-program remodeling from paired ssGSEA analysis.**

**a,** Program remodeling volcano plot showing paired recurrent-primary ssGSEA score changes across Hallmark, Reactome, Oncogenic, and Immunologic gene-set collections.
**b,** Paired ssGSEA trajectory for HALLMARK_HYPOXIA.

**c,** Paired ssGSEA trajectory for HALLMARK_REACTIVE_OXYGEN_SPECIES_PATHWAY.
**d,** Paired ssGSEA trajectory for HALLMARK_INTERFERON_GAMMA_RESPONSE.
**e,** Paired ssGSEA trajectory for HALLMARK_EPITHELIAL_MESENCHYMAL_TRANSITION.
**f,** Paired ssGSEA trajectory for HALLMARK_UV_RESPONSE_UP.
**g,** Paired ssGSEA trajectory for GSE16385_IFNG_TNF_VS_IL4_STIM_MACROPHAGE.
**h,** Paired ssGSEA trajectory for GSE39820_CTRL_VS_IL1B_IL6_CD4_TCELL_UP.
**i,** Paired ssGSEA trajectory for PDGF_ERK_DN.V1_DN.
**j,** Paired ssGSEA trajectory for IL15_UP.V1_UP.
**k,** Paired ssGSEA trajectory for REACTOME_RHO_GTPASES_ACTIVATE_CIT.

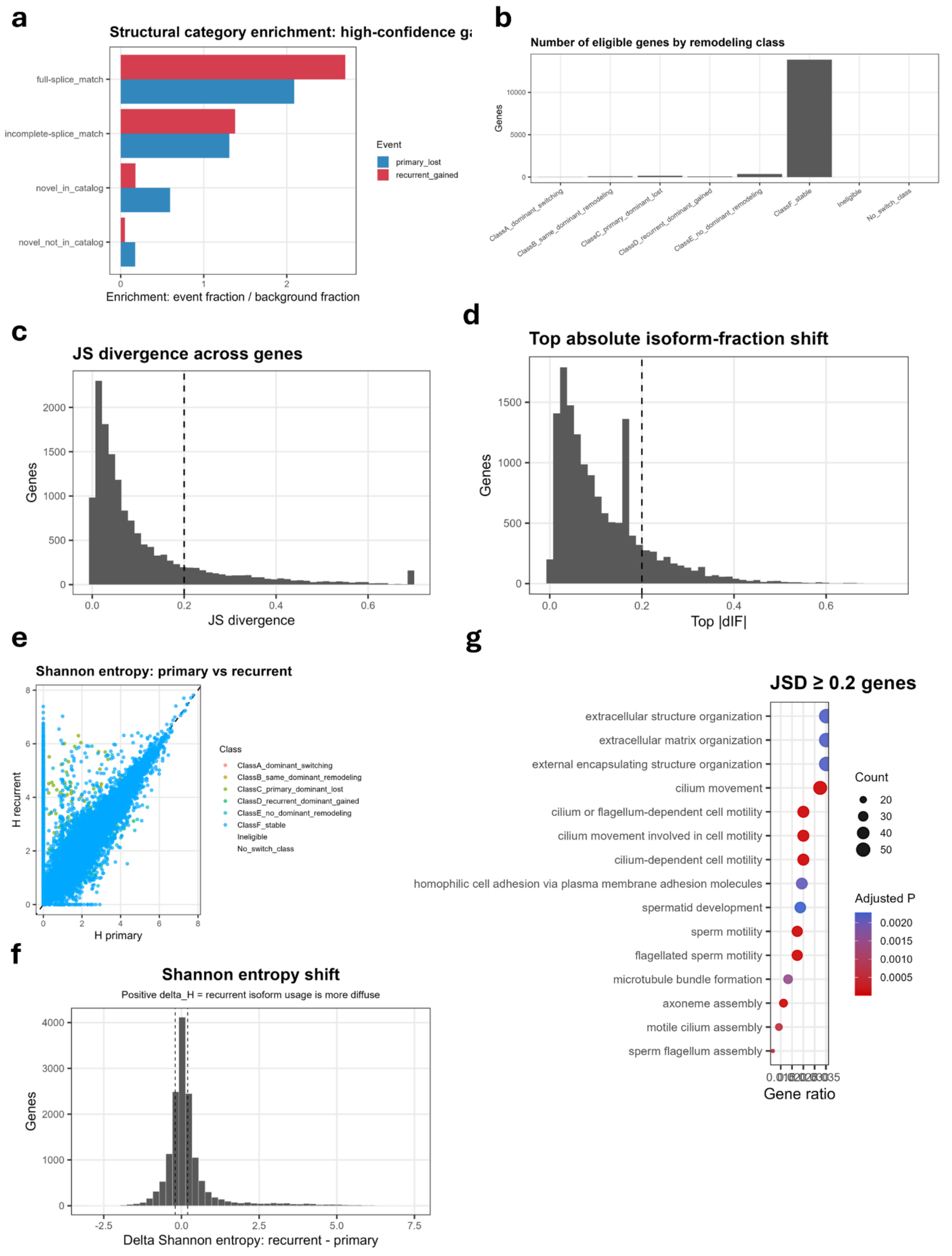


**Supplementary Figure 5. Isoform gain/loss, A–F remodeling classes, JSD/dIF thresholds, and entropy behavior.**

**a,** Structural-category enrichment among high-confidence recurrently gained and primary-lost isoforms relative to background expressed isoforms.
**b,** Number of eligible genes assigned to each A–F isoform remodeling class. Class A indicates dominant isoform switching; Class B, same dominant isoform with secondary remodeling; Class C, primary dominant isoform lost; Class D, recurrent dominant isoform gained; Class E, no-dominant remodeling; and Class F, stable or not high-confidence remodeled.
**c,** Distribution of gene-level Jensen–Shannon divergence values, with the high-confidence threshold marked.
**d,** Distribution of gene-level maximum absolute delta isoform fraction values, with the high-confidence threshold marked.
**e,** Scatter plot comparing primary and recurrent Shannon entropy for gene-level isoform usage distributions.
**f,** Distribution of recurrent-primary Shannon entropy shifts; positive values indicate more diffuse recurrent isoform usage, whereas negative values indicate more focused recurrent isoform usage.
**g,** Gene Ontology biological process over-representation analysis of high-JSD genes, highlighting extracellular matrix, extracellular structure, cilium, motility, microtubule, and axoneme-associated processes.

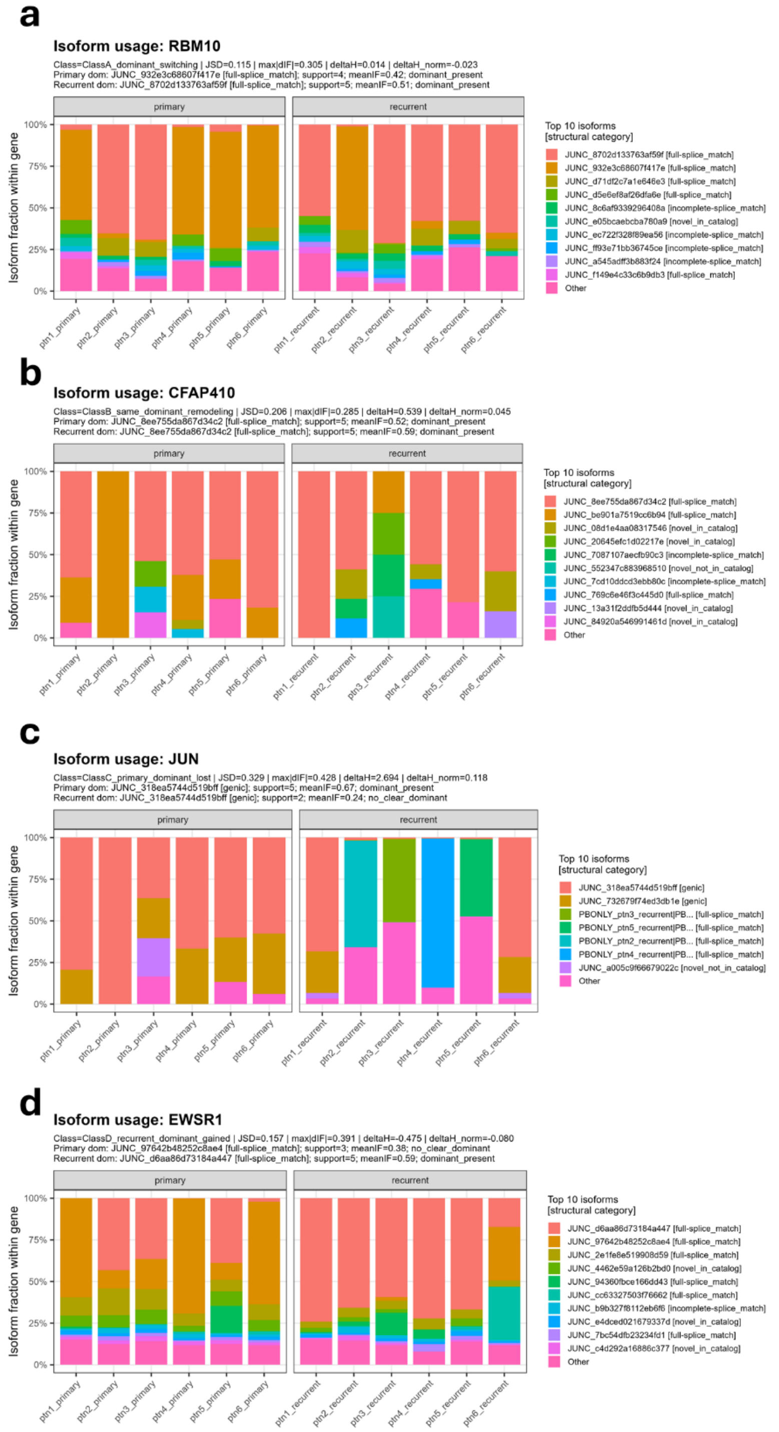


**Supplementary Figure 6. Representative isoform-usage examples from the JSD/dIF/entropy remodeling framework.**

**a,** Isoform-fraction trajectories for RBM10 across primary and recurrent tumors, illustrating remodeling of major and secondary isoforms within an expressed gene.
**b,** Isoform-fraction trajectories for CFAP410, showing patient-paired redistribution among multiple isoforms.

**c,** Isoform-fraction trajectories for JUN, showing recurrence-associated shifts in dominant or high-abundance isoforms.
**d,** Isoform-fraction trajectories for EWSR1, showing patient-paired isoform redistribution across primary and recurrent tumors.
Each plot shows the fraction of gene-level full-length reads assigned to the top isoforms, stratified by timepoint and patient.

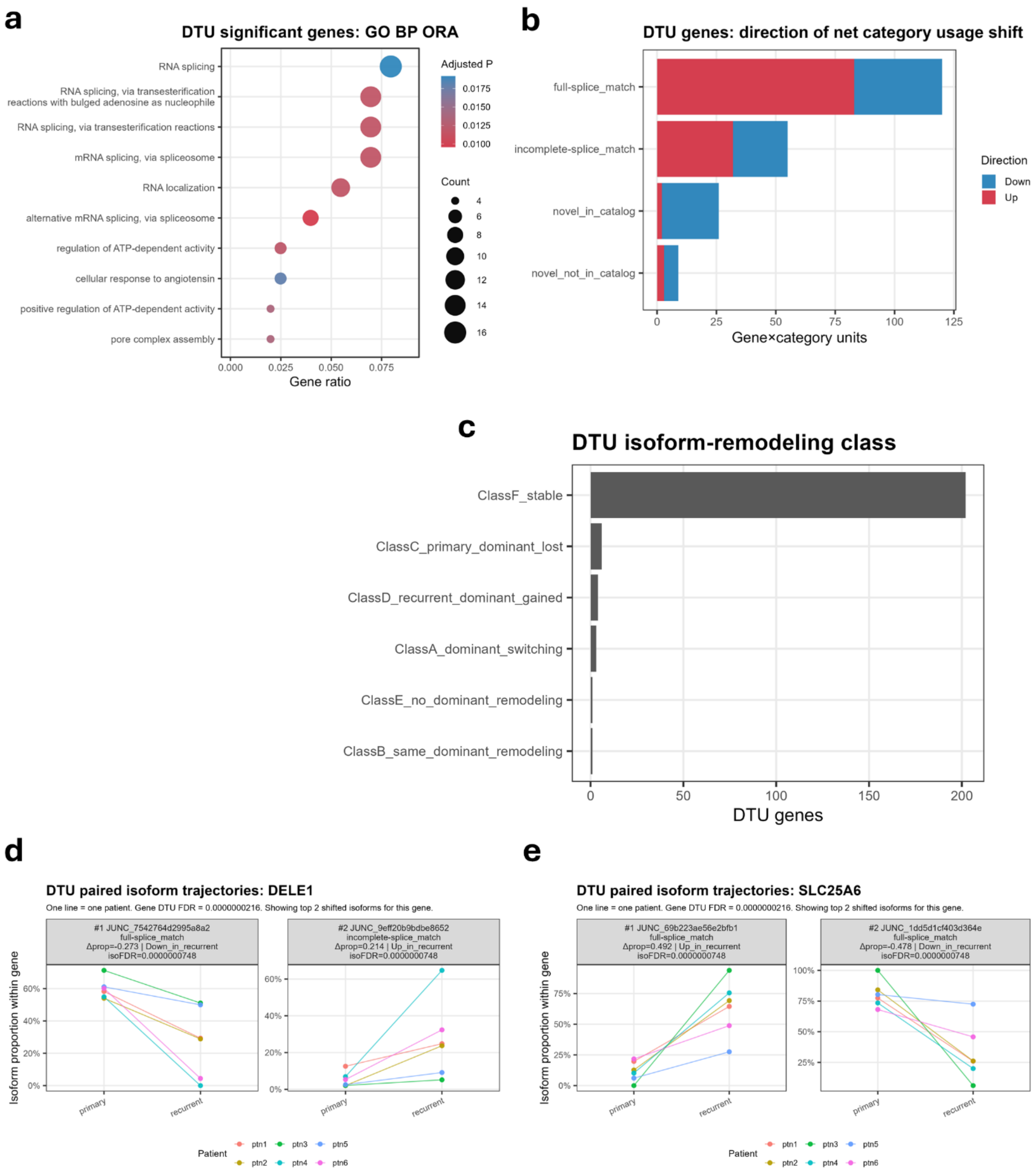


**Supplementary Figure 7. Differential transcript usage analysis and representative DTU isoform trajectories.**

**a,** Gene Ontology biological process over-representation analysis of DTU-significant genes, highlighting RNA splicing, mRNA splicing via spliceosome, RNA localization, and related RNA-processing programs.
**b,** Direction of DTU-associated isoform usage shifts by Pigeon structural category.
**c,** DTU-significant genes stratified by A–F isoform-remodeling class, showing the relationship

between statistical DTU and the effect-size/dominance-based remodeling framework.
**d,** Paired isoform trajectories for DELE1, showing reciprocal primary-to-recurrent changes in the top shifted isoforms across patients.
**e,** Paired isoform trajectories for SLC25A6, showing recurrent gain of one isoform and loss of another across patients.

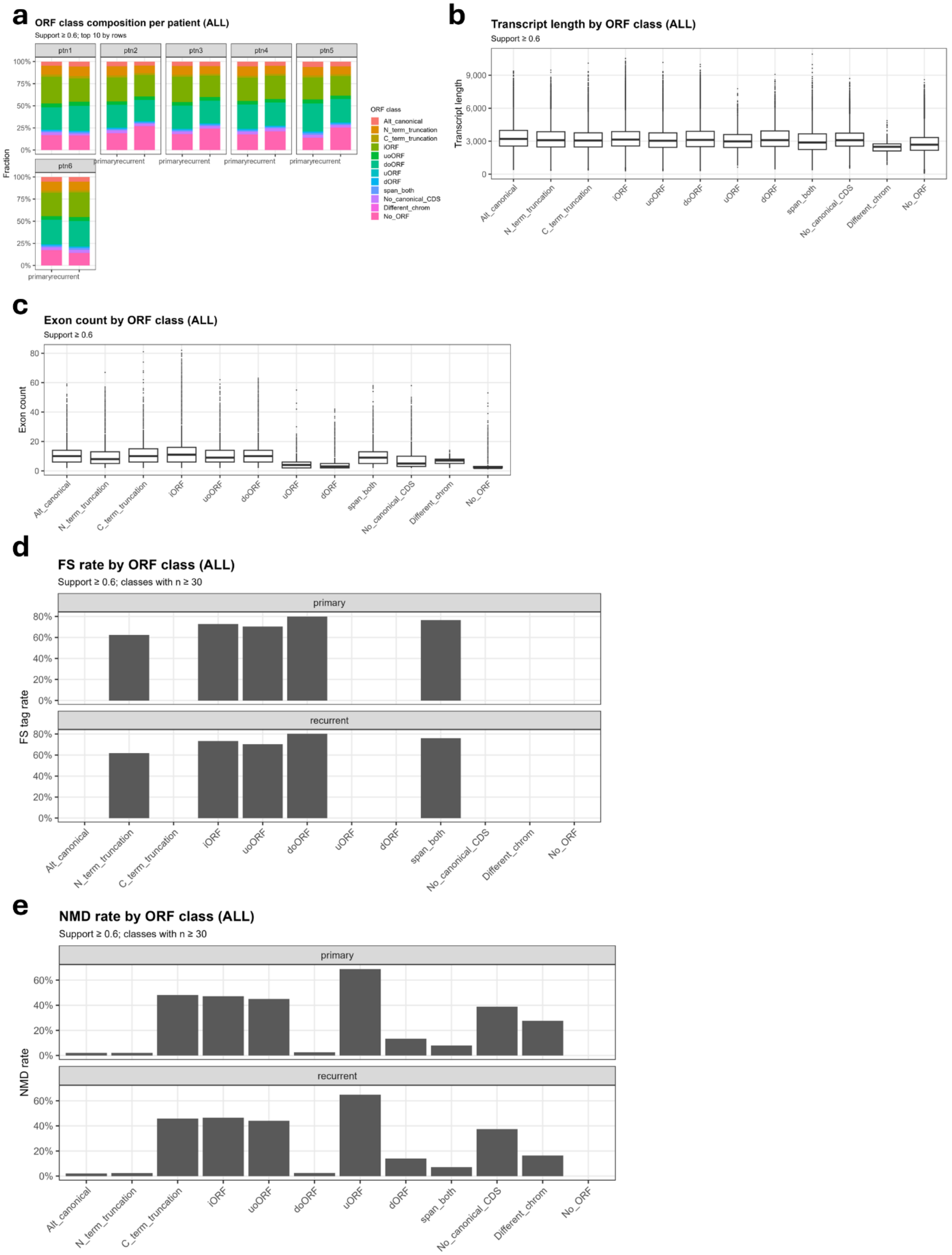


**Supplementary Figure 8. Supporting ORF annotation features across patients and ORF classes.**

**a,** Patient-level ORF-class composition across primary and recurrent tumors.
**b,** Transcript length distribution stratified by predicted ORF class.
**c,** Exon count distribution stratified by predicted ORF class.
**d,** Frameshift-rate summary across predicted ORF classes in primary and recurrent tumors.
**e,** NMD-rate summary across predicted ORF classes in primary and recurrent tumors.
These panels support the internal consistency of the ORF annotation framework and show that predicted ORF classes differ in expected transcript-structure and consequence features.

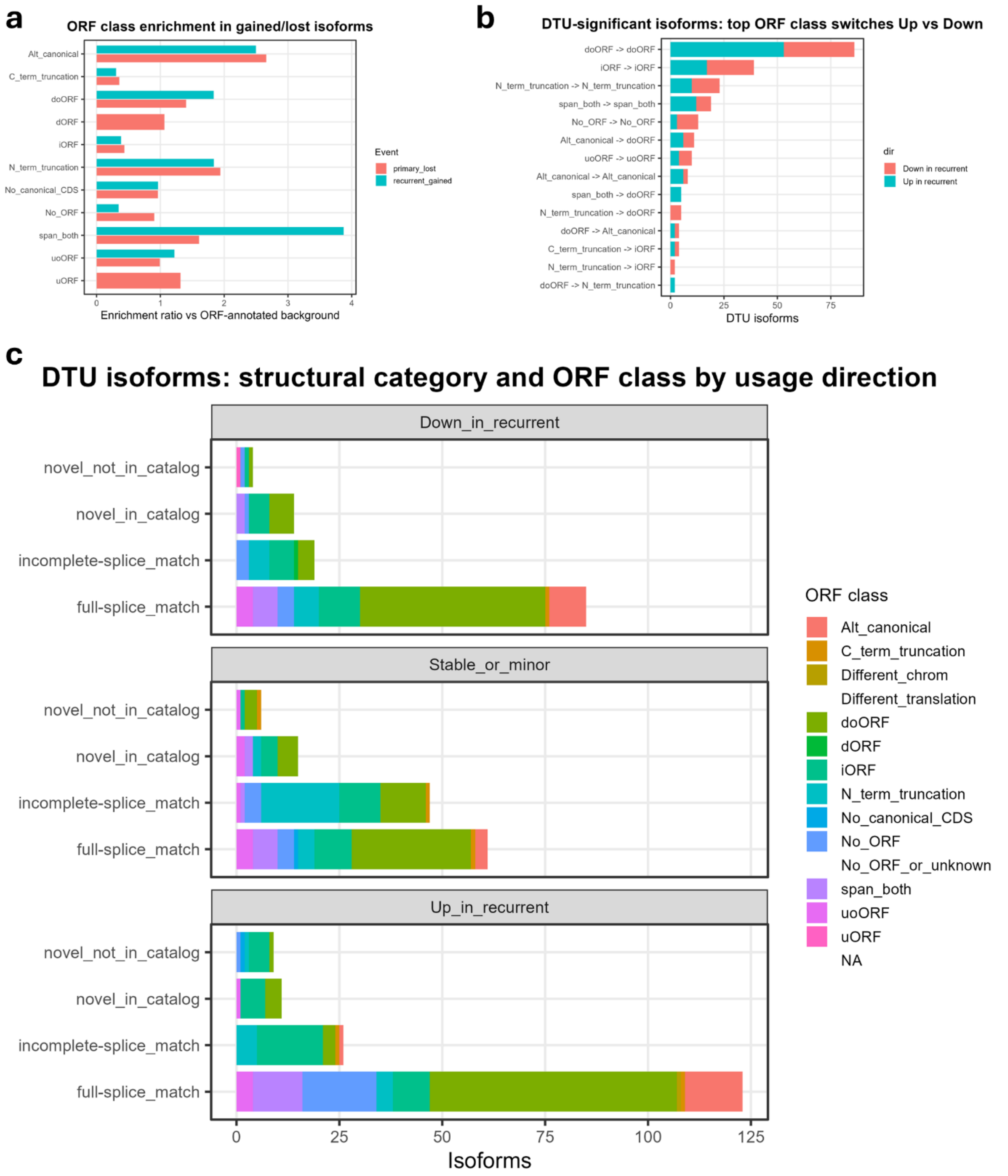


**Supplementary Figure 9. ORF enrichment and DTU-linked ORF remodeling.**

**a,** ORF-class enrichment among high-confidence primary-lost and recurrent-gained isoforms relative to the ORF-annotated background.
**b,** Top predicted ORF-class switches among DTU-significant isoforms, stratified by whether isoform usage increased or decreased in recurrence.
**c,** DTU-associated isoforms stratified jointly by Pigeon structural category, predicted ORF class, and

recurrent usage direction.
These panels show that recurrence-associated isoform gain/loss and DTU events are associated with selective predicted ORF-state remodeling rather than uniform shifts across all transcript classes.

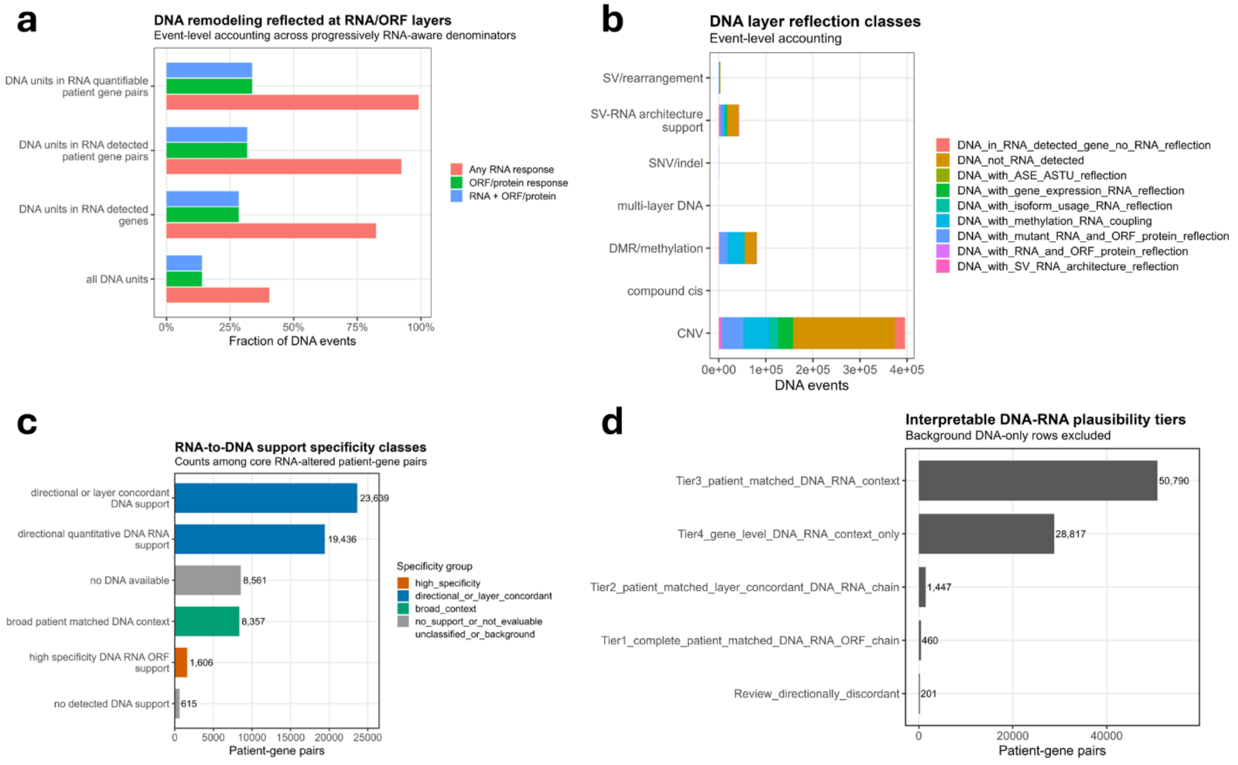


**Supplementary Figure 10. Extended denominator, reflection, support-specificity, and plausibility summaries for DNA–RNA integration.**

**a,** DNA remodeling reflected at RNA and predicted ORF/protein-fate layers. Fractions are recalculated within progressively RNA-aware DNA denominators and indicate DNA events with any RNA response, predicted ORF/protein-fate response, or combined RNA plus ORF/protein-fate response.

**b,** DNA-layer reflection classes across DNA event families, showing how different DNA layers map to RNA-aware and predicted ORF/protein-fate-linked reflection classes.

**c,** RNA-to-DNA support specificity classes shown as patient-gene pair counts among core RNA-altered patient-gene pairs. Classes separate high-specificity DNA–RNA/ORF support, directional/layer-concordant support, broad patient-matched DNA context, no DNA available, and no detected DNA support.

**d,** Interpretable DNA–RNA plausibility classes after excluding background DNA-only rows. Complete patient-matched DNA–RNA–ORF chains and layer-concordant DNA–RNA chains are shown alongside broader patient-matched DNA–RNA context, gene-level DNA–RNA context only, and directionally discordant review rows.

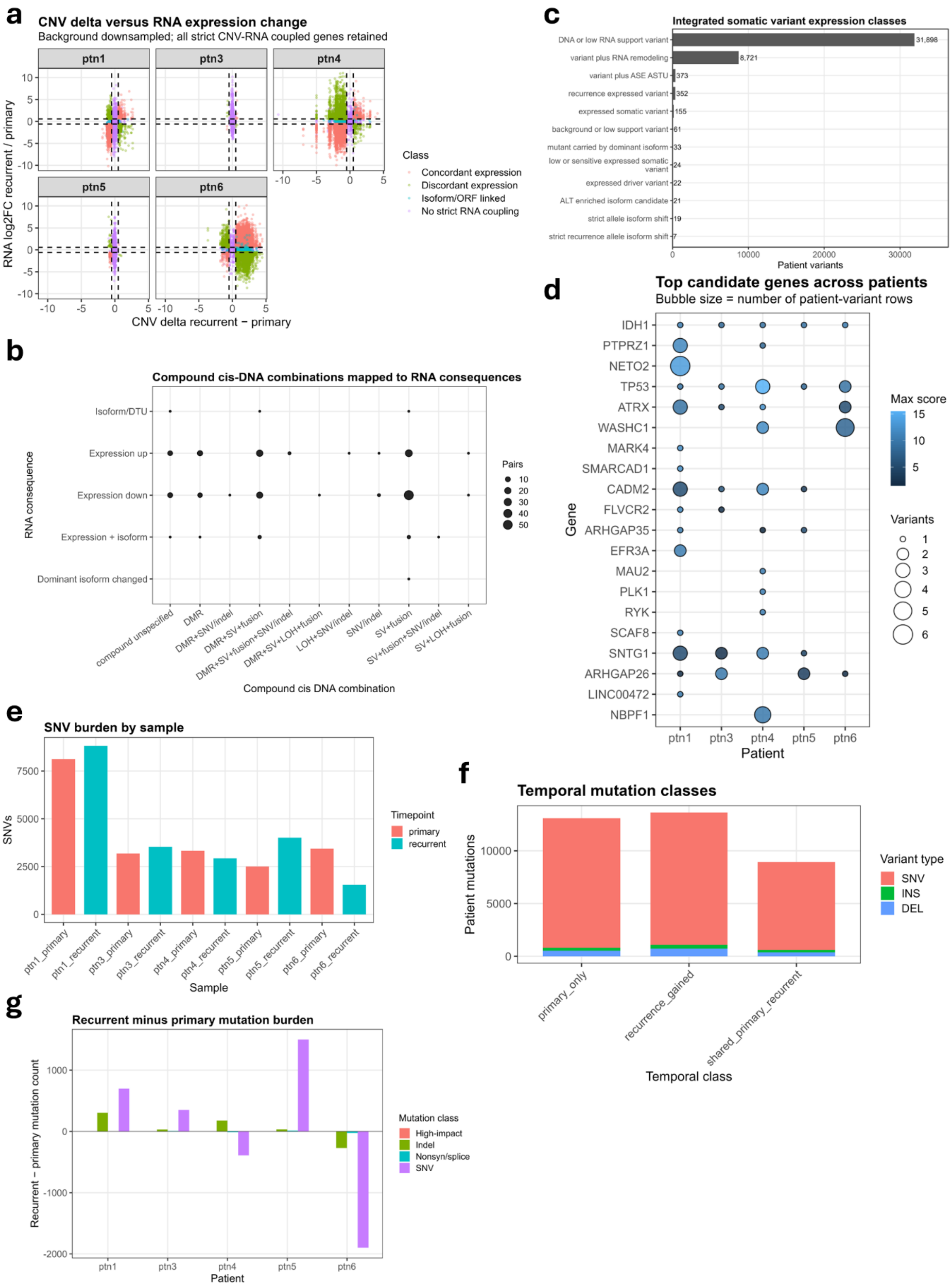


**Supplementary Figure 11. Extended CNV, compound cis-event, somatic-variant, and mutation-burden integration analyses.**

**a,** Recurrent-primary CNV delta versus recurrent-primary RNA expression change by patient. Points are colored by RNA-coupling class, including concordant expression, discordant expression, isoform/ORF-linked events, and events without strict RNA coupling.

**b,** Compound cis-DNA combinations mapped to RNA consequence classes. Point size indicates the number of patient-gene pairs for each compound DNA-layer combination and RNA consequence category.

**c,** Integrated somatic variant expression classes across patient variants, including DNA-only or low RNA-support variants, variant plus RNA remodeling, variant plus ASE/ASTU, expressed recurrent variants, expressed somatic variants, mutant-dominant isoforms, ALT-enriched isoform candidates, and strict recurrence allele/isoform-shift classes.

**d,** Top candidate genes across patients from the integrated somatic variant analysis. Bubble size indicates the number of patient-variant rows and fill intensity indicates maximum integrated variant score.

**e,** SNV burden by sample, stratified by timepoint.

**f,** Temporal mutation classes across patient mutations, including primary-only, recurrent-only, and shared primary-recurrent mutation classes.

**g,** Recurrent minus primary mutation burden by patient and mutation class, including SNV, indel, nonsynonymous/splice, and high-impact-like mutation categories.

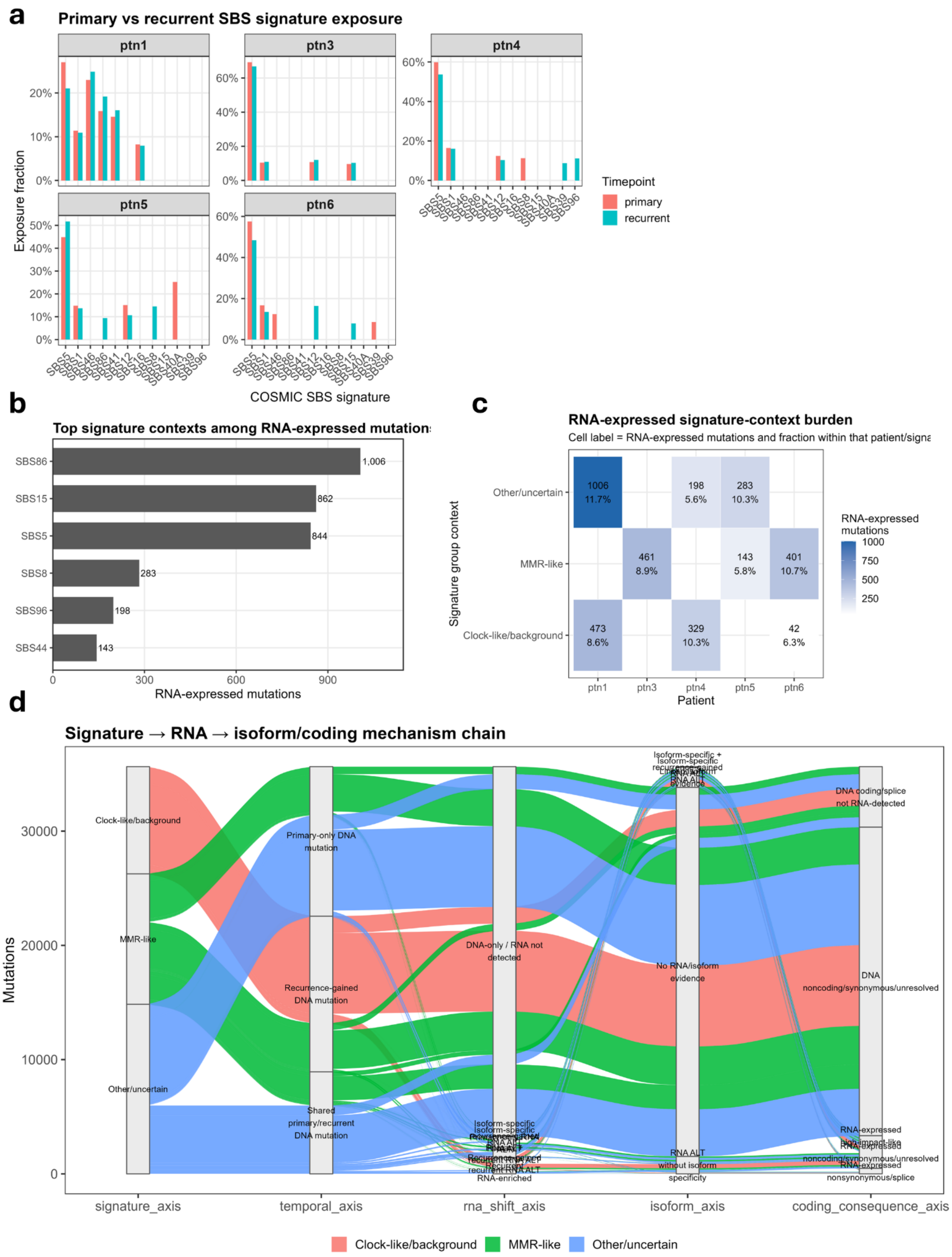


**Supplementary Figure 12. Mutational-signature context for RNA-expressed somatic variants.**

**a,** Primary versus recurrent COSMIC SBS signature exposure by patient and timepoint.

**b,** Top COSMIC SBS signature contexts among RNA-expressed mutations.

**c,** Patient-level burden of RNA-expressed signature-context mutations stratified by signature group, including clock-like/background, MMR-like, and other/uncertain groups. Cell labels indicate RNA-expressed mutation counts and within-patient/signature-group fractions.

**d,** Signature-to-RNA-to-isoform/coding mechanism alluvial. Flows connect signature group, temporal mutation class, RNA shift class, isoform-level mutant RNA evidence, and predicted consequence class.

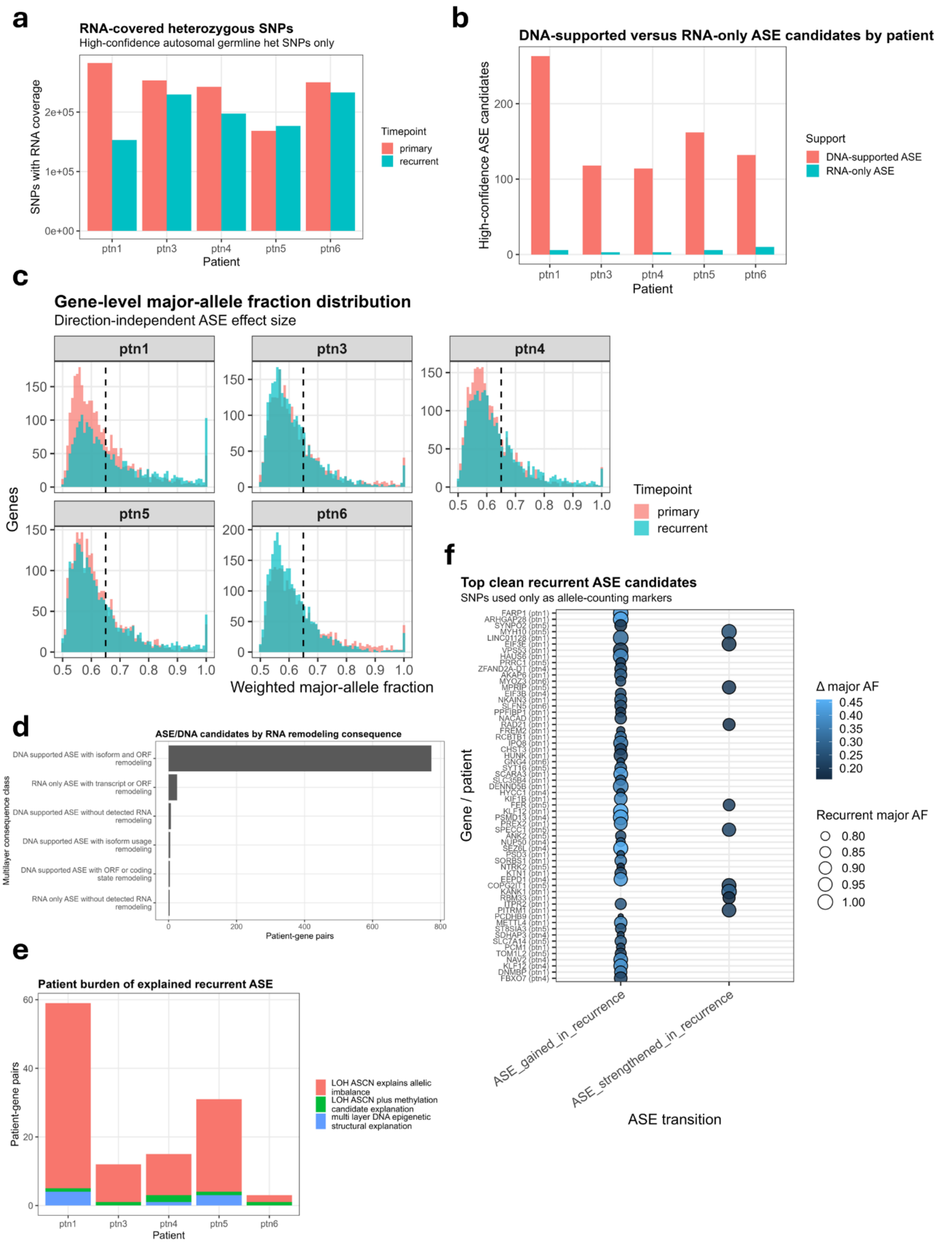


**Supplementary Figure 13. Extended ASE marker coverage, DNA support, and consequence analyses.**

**a,** RNA-covered heterozygous SNPs by patient and timepoint, restricted to high-confidence autosomal germline heterozygous SNP markers.

**b,** DNA-supported versus RNA-only ASE candidates by patient. DNA-supported ASE indicates ASE candidates with patient-matched DNA support, whereas RNA-only ASE lacks detected patient-specific DNA support.

**c,** Gene-level weighted major-allele fraction distributions by patient and timepoint. The dashed line marks the major-allele fraction threshold used to identify direction-independent ASE effect size.

**d,** ASE/DNA candidates stratified by RNA remodeling consequence, including DNA-supported ASE with isoform and ORF remodeling, RNA-only ASE with transcript or ORF remodeling, and other lower-frequency consequence classes.

**e,** Patient burden of explained recurrent ASE, stratified by primary explanatory mechanism, including LOH/allele-specific CNV, unstable or candidate LOH/allele-specific CNV, and multi-layer DNA explanation.

**f,** Top clean recurrent ASE candidates. Bubble size indicates recurrent major-allele fraction, and color indicates change in major-allele fraction between primary and recurrent tumors.

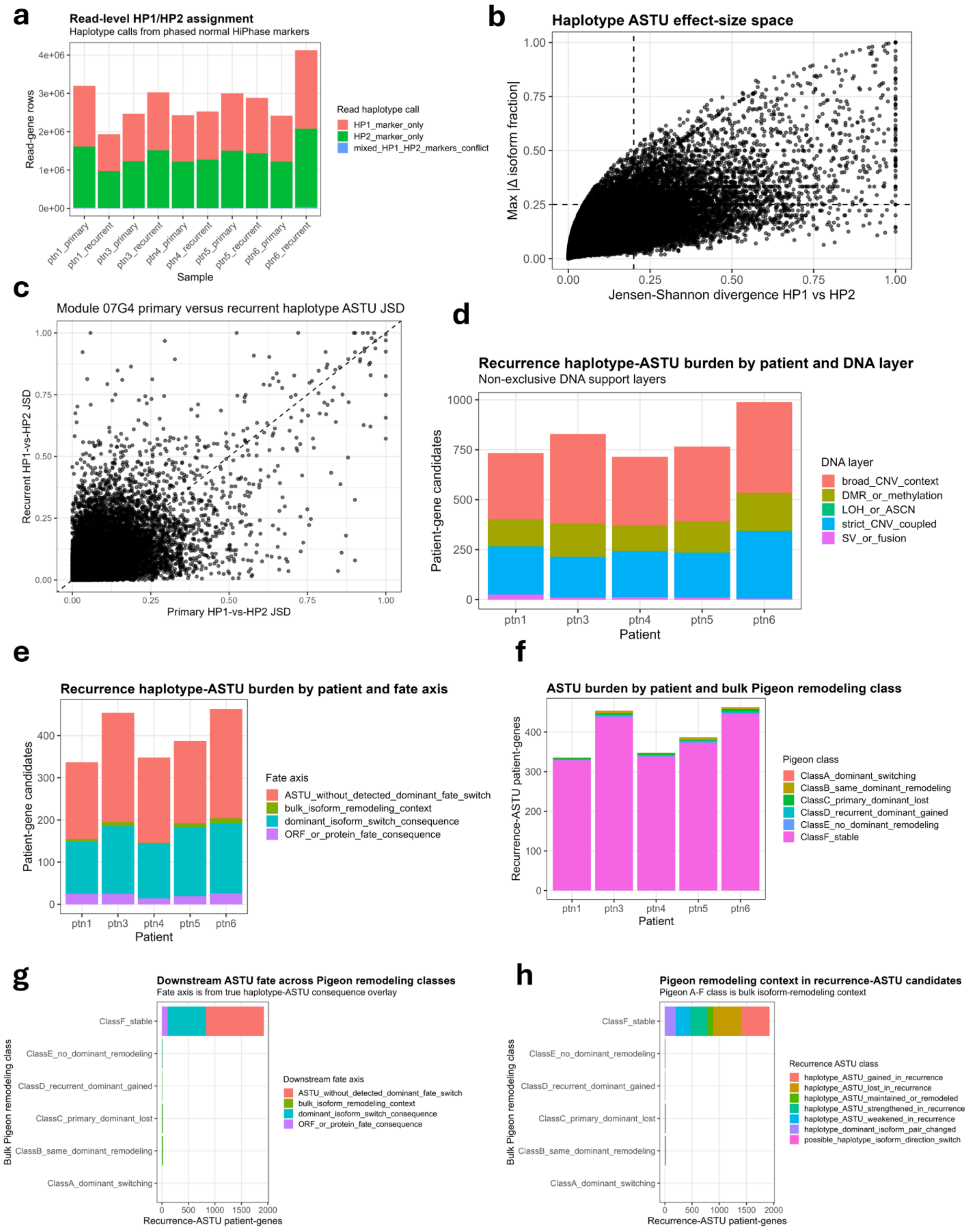


**Supplementary Figure 14. Extended ASTU marker assignment, effect-size, DNA-support, and fate-axis analyses.**

**a,** Read-level HP1/HP2 marker assignment across samples using phased normal haplotype markers. Bars show reads assigned to HP1-only, HP2-only, or conflicting HP1/HP2 marker categories.

**b,** Haplotype-marker ASTU effect-size space. Each point represents a gene-patient context, with Jensen–Shannon divergence comparing HP1 versus HP2 isoform distributions and maximum absolute isoform-fraction shift capturing the largest haplotype-specific isoform difference.

**c,** Primary versus recurrent haplotype-marker ASTU JSD. Points above or below the diagonal indicate patient-gene contexts in which haplotype-specific isoform divergence increased or decreased at recurrence.

**d,** Recurrence-associated haplotype-marker ASTU burden by patient and DNA support layer, including broad CNV context, DMR/methylation support, LOH/allele-specific CNV, strict CNV-coupled support, and SV/fusion support.

**e,** Recurrence-associated haplotype-marker ASTU burden by patient and downstream fate axis, including ASTU without detected dominant fate switch, bulk isoform-remodeling context, dominant isoform-switch consequence, and predicted ORF/protein-fate consequence.

**f,** Recurrence-associated ASTU burden by patient and bulk Pigeon isoform-remodeling class.

**g,** Downstream ASTU fate across bulk Pigeon isoform-remodeling classes, showing how ASTU-only, bulk isoform-remodeling, dominant isoform-switch, and predicted ORF/protein-fate categories distribute across A–F remodeling classes.

**h,** Bulk Pigeon remodeling context among true recurrence-ASTU candidates. Recurrence-ASTU classes include ASTU gained, lost, maintained or remodeled, strengthened, weakened, possible haplotype isoform direction switch, and possible direction switch.

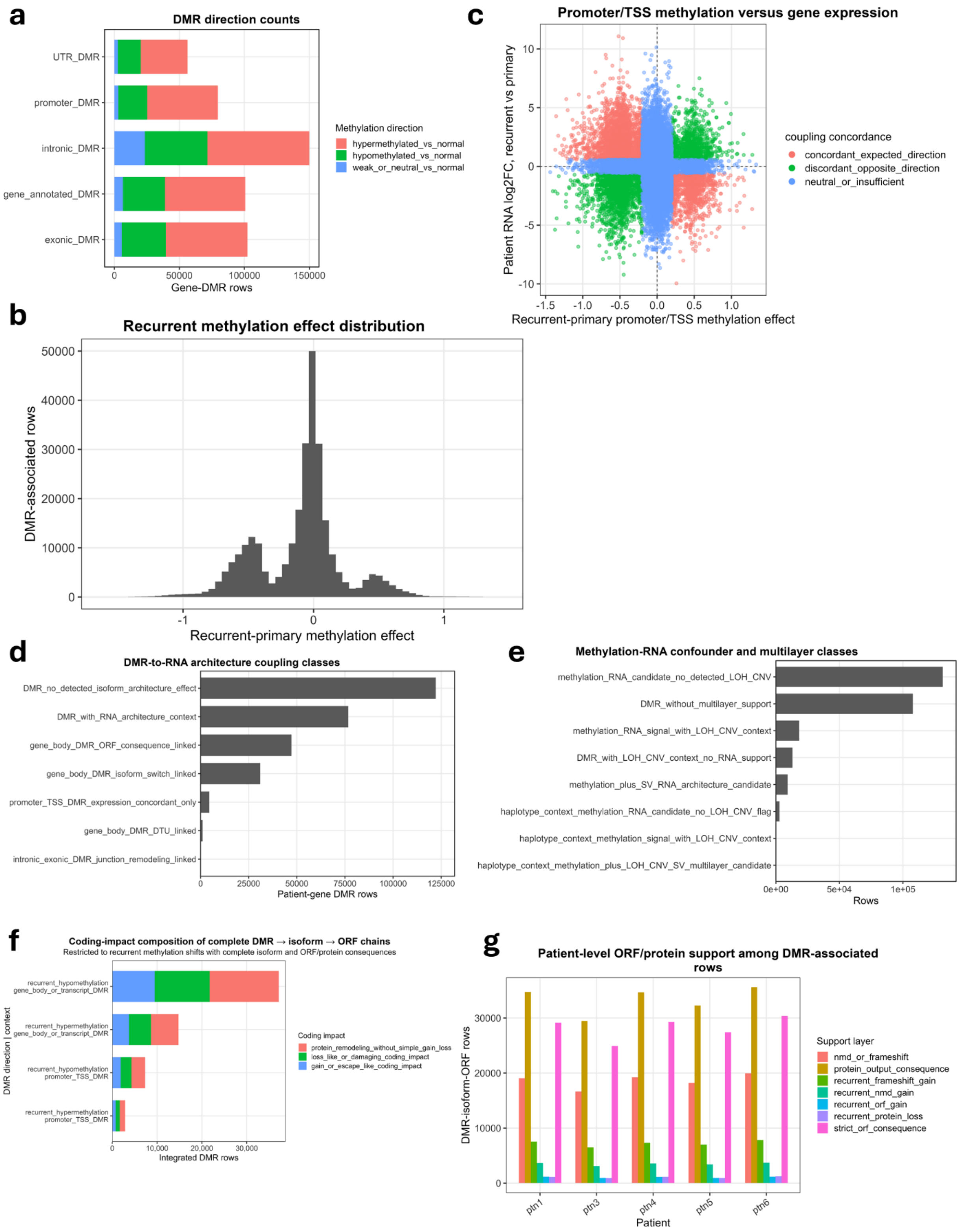


**Supplementary Figure 15. Extended DMR direction, methylation-RNA coupling, and coding-impact analyses.**

**a,** DMR direction by regulatory class. Patient-gene DMR rows are stratified by DMR context and direction relative to normal, including hypermethylated, hypomethylated, and weak or neutral methylation shifts.

**b,** Distribution of recurrent methylation effect used for RNA coupling. Positive values indicate stronger methylation in recurrence relative to primary; negative values indicate lower methylation in recurrence.

**c,** Promoter/TSS methylation effect versus recurrent-primary gene expression change. Each point represents a promoter/TSS DMR-associated patient-gene context.

**d,** DMR-to-RNA architecture coupling classes. Bars summarize DMR-associated patient-gene rows with no detected isoform architecture effect, RNA architecture context, gene-body DMR with ORF consequence, gene-body DMR with isoform switch, promoter/TSS DMR with expression concordance, gene-body DMR with DTU link, or intronic/exonic DMR with junction remodeling.

**e,** Methylation-RNA confounder and multilayer support classes. Bars separate methylation-RNA candidates without detected LOH/CNV context from DMRs with or without multilayer support and methylation-RNA signals occurring together with LOH/CNV, SV, or haplotype-context support.

**f,** Coding-impact composition of complete DMR–isoform–ORF chains, stratified by recurrent methylation direction and DMR context. Coding-impact classes include ORF/protein gain or loss, coding remodeling without simple gain or loss, and protein remodeling without simple amino-acid gain or loss.

**g,** Patient-level ORF/protein consequence support among DMR-associated rows. Bars summarize support layers such as DMR-associated frameshift, recurrent frameshift gain, NMD gain, ORF gain, protein loss, protein-output consequence, and strict ORF consequence.

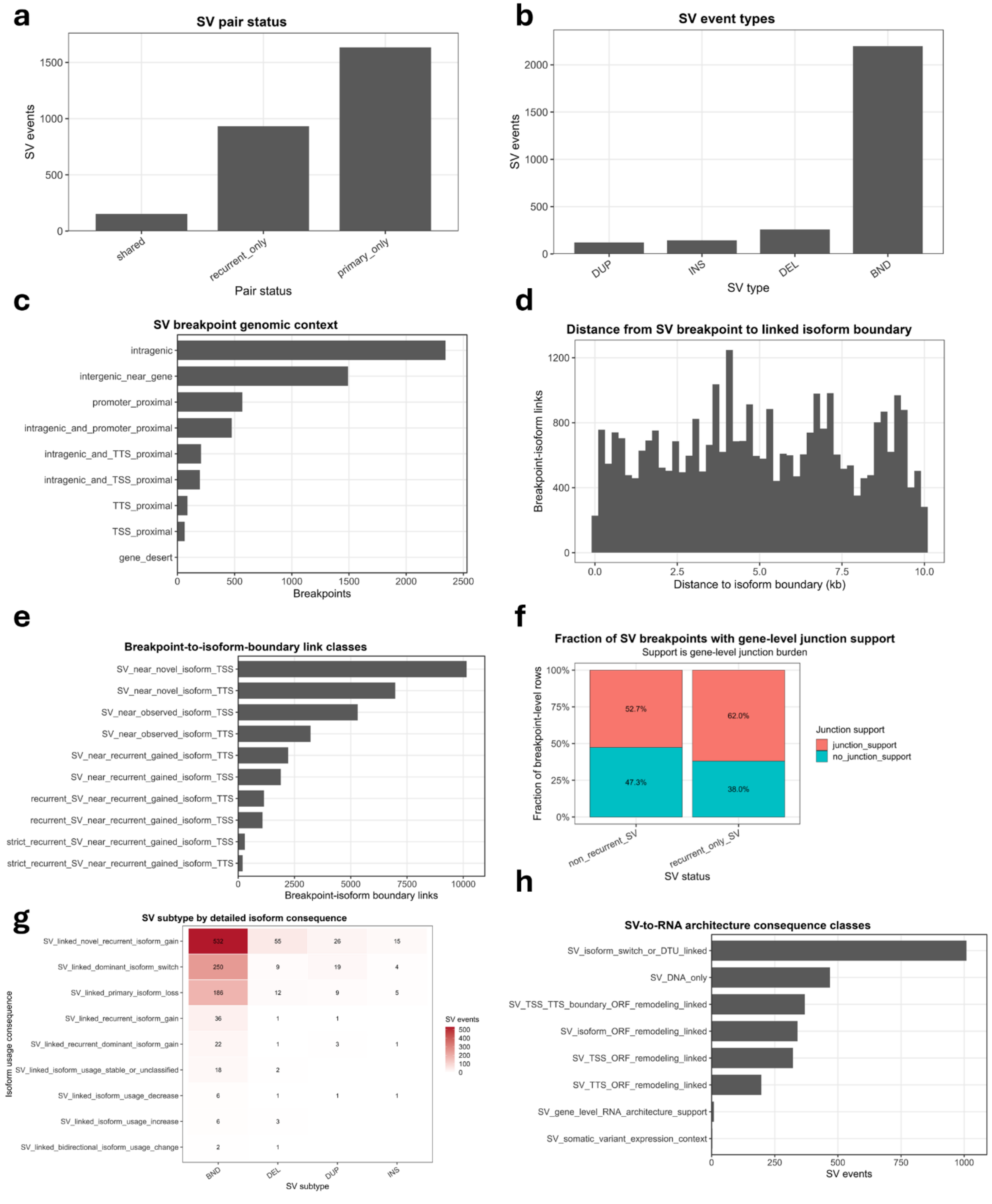


**Supplementary Figure 16. Extended structural-variant RNA architecture analyses.**

**a,** SV pair status across primary and recurrent tumors, stratified as primary-only, recurrent-only, or shared.

**b,** SV event types across the integrated SV set, including breakend-like, deletion, insertion, and duplication classes.

**c,** SV breakpoint genomic context, including intragenic, near-gene, promoter-proximal, TSS/TTS-proximal, and gene-desert categories.

**d,** Distance from SV breakpoints to linked isoform boundaries, restricted to candidate breakpoint-isoform links within the plotted distance range.

**e,** SV breakpoint-to-isoform-boundary link classes, including links near novel isoform TSS/TTS, observed isoform boundaries, and recurrently gained isoform boundaries.

**f,** Fraction of SV breakpoint-gene links with gene-level RNA junction support among recurrent and recurrence-gained SVs.

**g,** SV subtype by detailed isoform consequence. Heatmap values indicate SV event counts across SV subtype and linked isoform usage consequence class.

**h,** SV-to-RNA architecture consequence classes, including SV-linked isoform switch or DTU-like remodeling, DNA-only SVs, TSS/TTS-boundary ORF remodeling, isoform-ORF remodeling, gene-level RNA architecture support, and somatic-variant expression context.

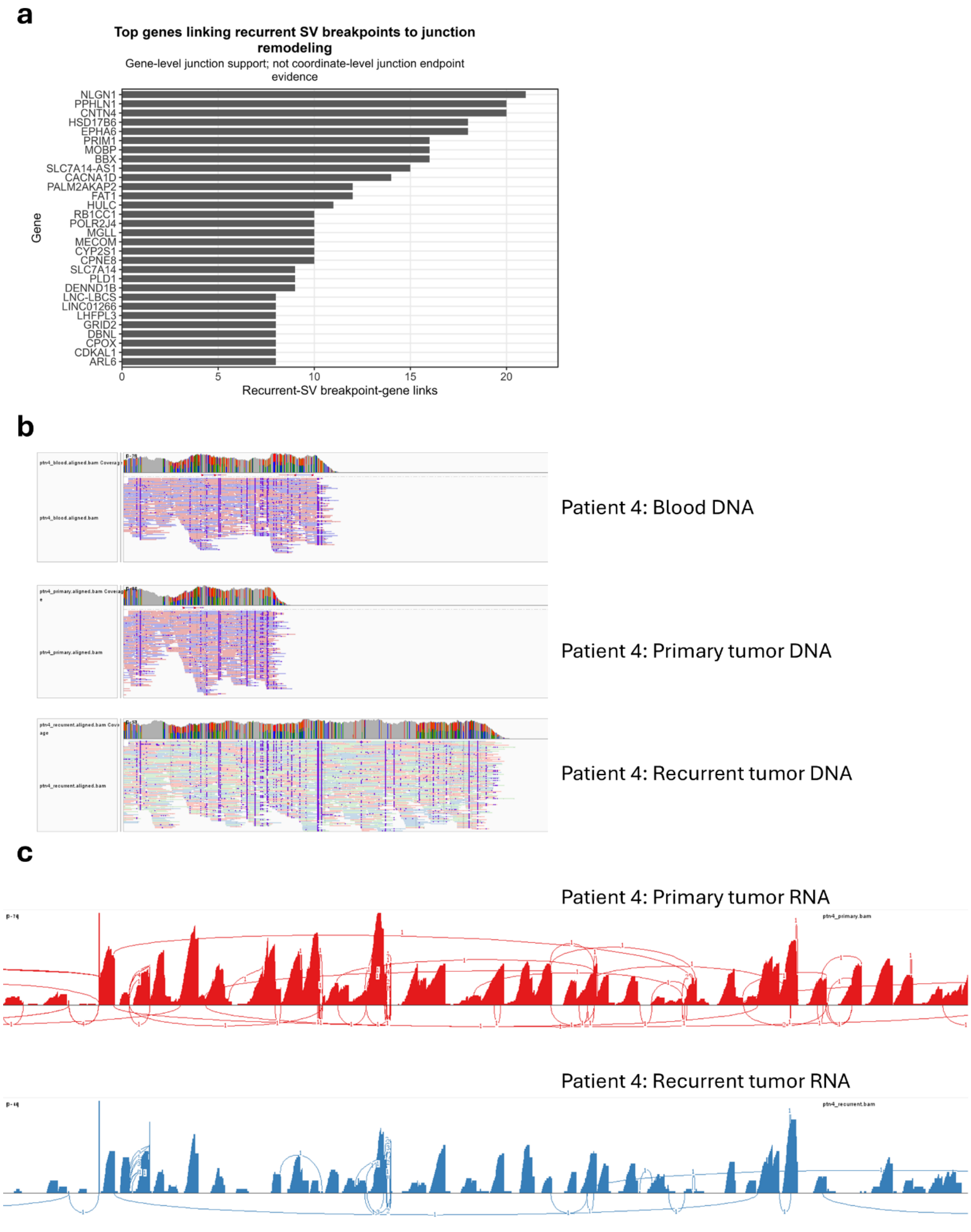


**Supplementary Figure 17. Representative SV-linked transcript-architecture remodeling at NLGN1.**

**a,** Top genes linking recurrent SV breakpoints to junction remodeling. Bars show the number of recurrent SV breakpoint-gene links with gene-level junction support.

**b,** IGV DNA tracks at the NLGN1 locus in patient 4, showing blood, primary tumor, and recurrent tumor DNA read alignments across the candidate SV-remodeled region.

**c,** RNA sashimi plots at the NLGN1 locus in patient 4, comparing primary and recurrent RNA junction architecture across the candidate SV-remodeled region.

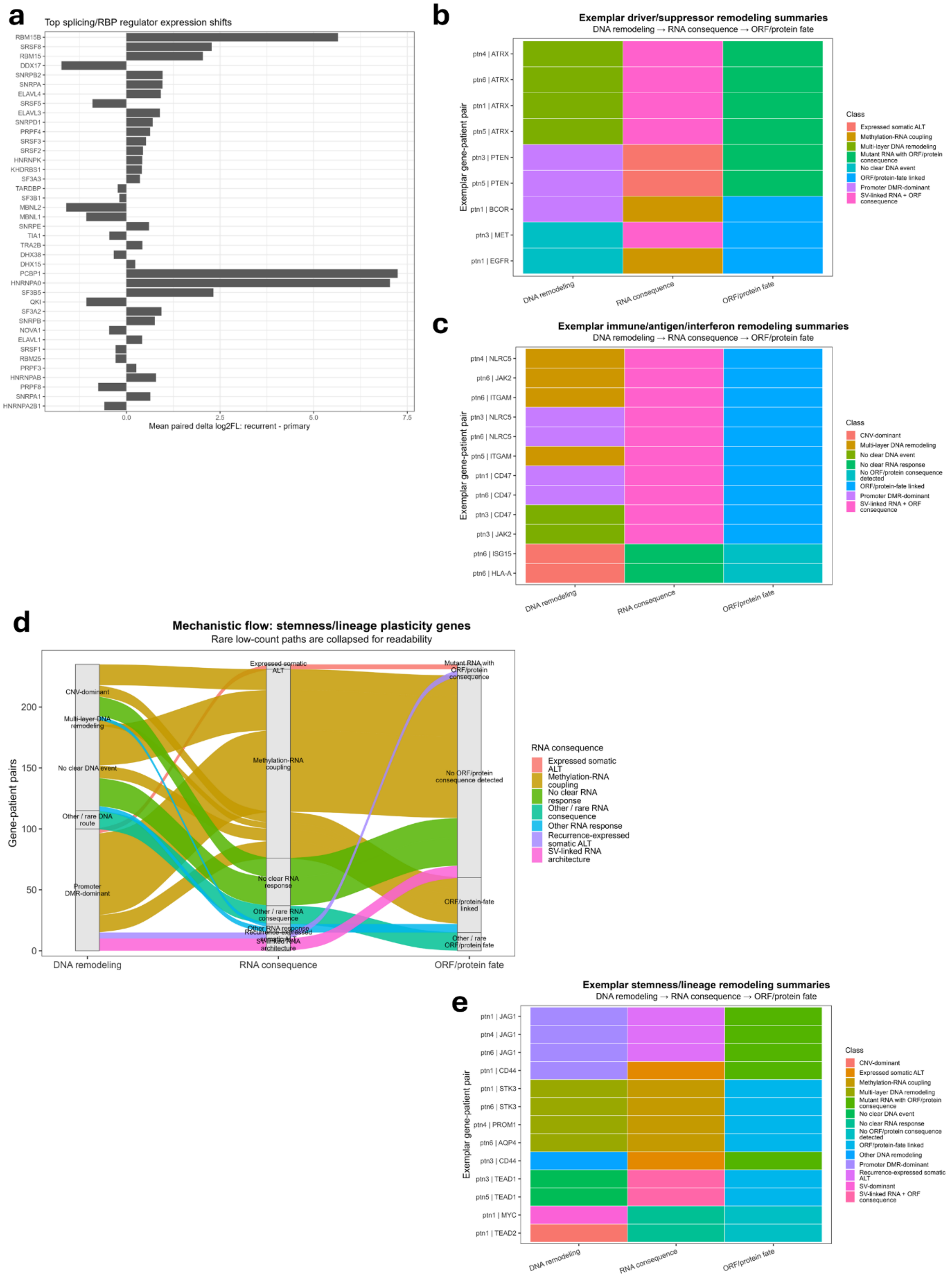

**Supplementary Figure 18. Exemplar biological gene-set summaries for splicing/RBP, driver/suppressor, immune, and stemness/lineage programs.**

**a,** Top splicing and RNA-binding protein regulator expression shifts. Bars show mean paired recurrent-primary log2 expression change for selected splicing/RBP regulators.

**b,** Exemplar driver/suppressor remodeling summaries. Selected gene-patient pairs are summarized across DNA mechanism class, RNA consequence class, and predicted ORF/protein-fate class.

**c,** Exemplar immune, antigen-presentation, and interferon remodeling summaries. Selected gene-patient pairs are summarized across DNA mechanism class, RNA consequence class, and predicted ORF/protein-fate class.

**d,** Collapsed mechanistic flow for stemness and lineage-plasticity genes. Flows connect DNA remodeling class, RNA consequence class, and predicted ORF/protein-fate class, with rare low-count paths collapsed for readability.

**e,** Exemplar stemness and lineage-plasticity remodeling summaries. Selected gene-patient pairs are summarized across DNA mechanism class, RNA consequence class, and predicted ORF/protein-fate class.
